# SPECTRAM: three-dimensional full-vector simulation of coherence-tunable microscopy images

**DOI:** 10.64898/2026.11.24.768465

**Authors:** Jiaqi Yao, Xi Chen

## Abstract

We present SPECTRAM, a physics-based framework for three-dimensional simulation of coherence-tunable microscopy image formation. Combining finite-difference time-domain solutions of Maxwell’s equations with a coherence-tunable, polarization-dependent image-formation model, SPECTRAM captures full-vector light scattering and modality-specific optical processing in brightfield, differential interference contrast, and phase-contrast microscopy. It supports arbitrary three-dimensional sample geometries and generates volumetric image stacks under configurable illumination and objective parameters. Validated against Mie theory and experimental images of polystyrene beads, SPECTRAM further reproduces imaging features of an iPSC-derived stem cell using a refractive-index distribution estimated by gradient light interference microscopy. SPECTRAM provides a versatile, physics-based tool for quantitative three-dimensional simulation of phase-sensitive microscopy.

## 1. Introduction

Phase-sensitive optical microscopy techniques, including brightfield, differential interference contrast (DIC), and phase-contrast microscopy, are widely used for label-free visualization of biological specimens [1–3]. These techniques convert variations in the optical field induced by a specimen’s refractive index distribution into image intensity. However, the detected intensity arises from the interference of optical fields and depends on the illumination source, optical system configuration, and specimen properties [4–8]. These dependencies can introduce image artifacts and complicate the interpretation of image contrast, most pronounced in the axial direction [9–12]. Consequently, identifying specific cellular components and quantitatively characterizing cellular properties from these images can be challenging, especially when high measurement accuracy is required. In thick and heterogeneous specimens, multiple scattering further modifies the propagating electromagnetic field, making the relationship between specimen properties and image contrast increasingly complex [13–17].

These challenges motivate accurate forward modeling of phase-sensitive optical microscopy that captures both the propagation of complex optical fields through the specimen and their subsequent transformation by the microscope’s optical system. Early approaches to optical image formation were largely based on scalar diffraction theory, including coherent point-spread-function (PSF) and transfer-function formulations [18–23]. For modeling three-dimensional (3D) specimen structure, diffraction tomography established a foundation for recovering refractive-index distributions from complex scattered fields under weak-scattering assumptions [24,25]. Subsequent approaches employed the first-order Born and Rytov approximations to describe wave scattering while retaining the complex amplitude and phase of the optical field [26,27]. Although widely used in diffraction tomography, these approximations have limited validity for strongly scattering specimens and do not fully account for higher-order multiple scattering.

To extend phase-image modeling to thick specimens, a product-of-convolutions (POC) model was developed that represents axial phase contributions by dividing the specimen into slices and combining their contributions through PSF convolutions [28]. Other wave-based approaches have subsequently addressed multiple scattering through beam propagation and multislice methods, which iteratively propagate complex optical fields through successive specimen layers to account for multiple-scattering interactions [29,30]. However, conventional multislice methods rely on a paraxial approximation and do not adequately capture highly oblique scattering or backward-scattered fields. The multilayer Born model extended the first Born approximation across multiple finite-thickness specimen layers, enabling the modeling of both forward- and backward-scattered fields [31]. Nevertheless, it retains the first Born approximation within each layer and therefore does not fully capture repeated multiple-scattering interactions within individual layers. Nonlinear integral-equation approaches based on the Lippmann–Schwinger formulation overcome this limitation by accounting for higher-order scattering interactions, in which the scattered fields contribute to subsequent scattering events [32]. However, their implementation for 3D specimens involves substantial computational and memory requirements, while scalar formulations do not capture the full vectorial nature of electromagnetic light–sample interactions. Conventional intensity-based light-transport methods, such as radiative transfer and Monte Carlo simulations, are also used to model multiply scattering thick specimens but generally do not preserve the optical phase required for direct simulation of interference-based microscopy [33,34]. Despite these advances, many wave-propagation approaches employ scalar-field approximations and therefore do not fully capture vectorial electromagnetic interactions and polarization-dependent effects, particularly at high numerical apertures (NAs).

Full-vector electromagnetic solvers, including finite-difference time-domain (FDTD), pseudo-spectral time-domain (PSTD), and finite-element methods (FEM), provide rigorous solutions of Maxwell’s equations and can account for electromagnetic scattering, interference, and polarization effects [35–38]. Beyond specimen-level scattering calculations, these solvers have also been incorporated into phase-sensitive microscope image-formation models. Full-vector FDTD was applied to broadband microscopy under partially coherent and unpolarized illumination [39]. The broader electromagnetic microscopy framework incorporated electromagnetic scattering, optical collection, and refocusing to synthesize images for multiple microscopy configurations, including brightfield, darkfield, and phase-contrast microscopy [40]. This approach was further extended to dispersive biological media [41]. However, its phase-contrast demonstration relied on an idealized reference-wave phase shift. A dedicated FDTD model explicitly incorporated annular illumination and phase-ring modulation, but approximated the illumination using sparse angular sampling at a fixed polar angle [42]. FDTD-based DIC imaging was also demonstrated by incorporating polarization-dependent beam shearing and recombination, although its use of a single axial illumination direction omitted the angular spread associated with a finite condenser NA [43]. Overall, these FDTD demonstrations primarily considered geometrically idealized samples and did not present volumetric z-stacks spanning an extended axial range. These methods also lacked tunable control over illumination coherence and did not investigate the influence of illumination NA on image formation. Although Fresnel-Kirchhoff diffraction has also enabled synthetic z-stacks [44,45], its scalar formulations neglect vectorial effects that become important at high collection NAs, while paraxial approximation can additionally introduce propagation-phase errors for high-angle components that accumulate over axial distance.

To address these limitations, we present SPECTRAM (**S**patial-frequency and **P**artial-coherence **E**lectromagnetic **C**omputation for 3D **R**econstruction **A**lgorithm in **M**icroscopy), a unified framework for simulating 3D image formation in brightfield, phase-contrast, and DIC microscopy. SPECTRAM combines full-vector FDTD simulations of light–sample interactions with a model of the complete microscope imaging system. SPECTRAM incorporates the source’s spatial-frequency distribution and experimentally informed coherence properties to account for the polarization, angular, and temporal spectra of the illumination, while combining contributions from different illumination angles according to their probability of mutual coherence. It accommodates arbitrary 3D specimen geometries and heterogeneous refractive-index distributions, while explicitly representing the refractive indices of the illumination-side, specimen, and collection-side media to account for reflections at their interfaces. The framework further incorporates modality-specific optical processing for brightfield, phase-contrast, and DIC microscopy. It calculates the detected optical signal from the normal component of the time-averaged Poynting vector, providing a more physically realistic representation of optical signal detection. Importantly, we establish a physically consistent model of objective-scanned volumetric image formation, enabling the generation of arbitrary images over extended axial ranges, allowing us to evaluate the accuracy along axial dimension with experimental data.

We validate SPECTRAM by comparing simulated and experimentally acquired 3D images of polystyrene beads across all three modalities and under different illumination NAs. We further demonstrate its applicability to an iPSC-derived stem cell, using a refractive-index distribution estimated by gradient light interference microscopy (GLIM). In summary, SPECTRAM enables systematic investigation of how illumination conditions, multiple scattering, and modality-specific optical processing shape 3D image formation, providing a physics-based foundation for evaluating imaging configurations and developing advanced phase-sensitive microscopy methods.

## 2. Methods

We first construct the simulation model for brightfield microscopy, which also serves as the common image pathway for other phase-sensitive modalities. The model employs Köhler illumination, a widely used configuration that provides uniform specimen illumination and minimizes illumination-induced intensity variations, thereby improving image contrast consistency (Fig. 1a). The collector lens establishes optical conjugation between the LED source plane and the condenser back focal plane, while the condenser front focal plane coincides with the specimen plane. Consequently, each source point in the condenser aperture plane generates a plane wave that illuminates the specimen, with its position determining the spatial frequency. For a sufficiently extended LED source, the partially coherent illumination model treats different source points as having random relative phases and therefore as mutually incoherent [46]. Upon interacting with the specimen, each incident plane wave generates a corresponding scattered field, which coherently combines with the incident field to form an interference pattern. The electromagnetic field is collected by the objective and mapped to its aperture plane in the spatial-frequency domain. The tube lens then relays it to the camera plane to form the detected image, with the magnification given by the ratio of the tube-lens focal length to the objective focal length [47].

**Fig 1.**
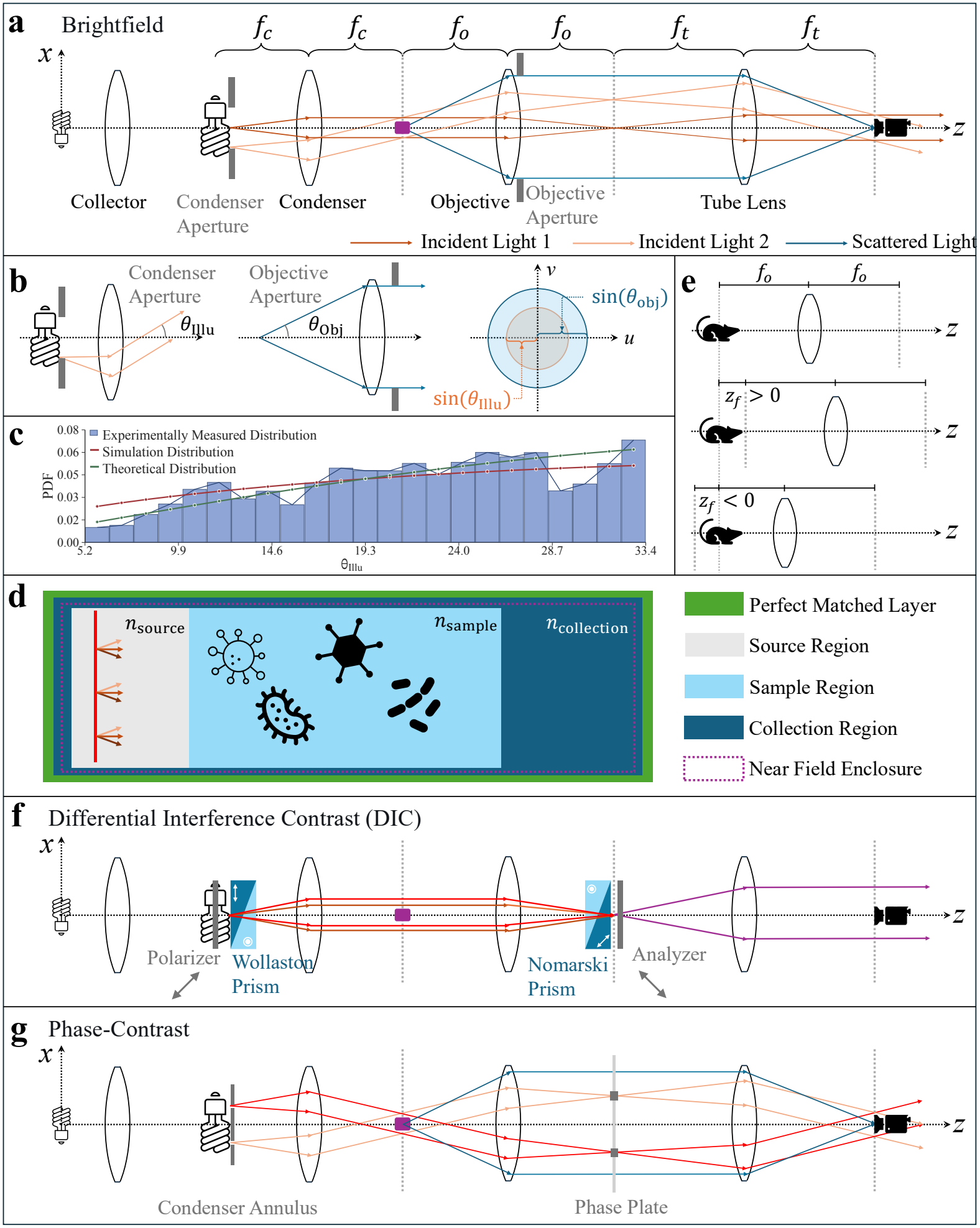
Optical configurations and computational framework of SPECTRAM. **(a)** Brightfield microscopy under Köhler illumination, showing the illumination, collection, and image-formation pathways. **(b)** Condenser and objective apertures defining the angular supports for illumination and collection, respectively. **(c)** Comparison of the experimentally measured angular distribution, the theoretical distribution assuming uniform condenser-aperture illumination, and the modified distribution adopted in the simulations. **(d) *x*** − ***z*** cross-section of the FDTD computational domain, comprising the source, specimen, and collection regions, surrounded by perfectly matched layers and enclosed by the near-field surface. **(e)** Objective scanning used to generate a *z*-stack by shifting the focal position relative to the fixed FDTD domain. **(f)** DIC microscopy configuration incorporating a polarizer, Wollaston prism, Nomarski prism, and analyzer. **(g)** Phase-contrast microscopy configuration incorporating a condenser annulus and an objective phase plate.

The broadband LED illumination is first decomposed into monochromatic components characterized by the angular frequency ω, with their normalized spectral distribution described by *p*_ω_(ω). At each fixed ω, the incident field is further represented as an angular spectrum of plane waves parameterized by the polar angle *θ* and azimuthal angle *ψ*. As shown in Fig. 1b, the optically conjugate condenser and objective apertures define circular angular supports for illumination and collection, respectively. Their angular ranges are determined by NA_Illu_ = *n*_source_ sin *θ*_Illu_ and NA_Obj_ = *n*_collection_ sin *θ*_obj_ where *n*_source_ and *n*_collection_ are the refractive indices of the corresponding regions, and *θ*_Illu_ and *θ*_obj_ are the maximum illumination and collection polar angles. The normalized angular distribution at a given ω is described by the conditional probability density *p*_*θ,ψ*_(*θ, ψ*| ω). As shown in Fig. 1c, assuming uniform illumination intensity across the condenser aperture plane yields the following theoretical PDF

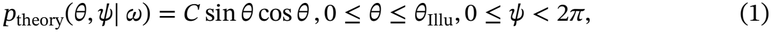

where *C* is the normalization factor. The derivation can be found in Supplementary Section 1L. However, measurements from the experimental microscope reveal a pronounced dip at large polar angles. To incorporate both the theoretical distribution and the experimentally observed behavior, we adopt the following modified PDF in our simulations:

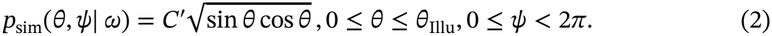

It assigns greater relative weight to contributions from incident light at smaller polar angles. Supplementary Section 5 presents the experimentally measured temporal-frequency spectrum *p*_ω_(ω) and angular distribution *p*_*θ,ψ*_(*θ, ψ*| ω) of the illumination. These measurements inform the idealized distributions used in SPECTRAM and provide the rationale for adopting the angular distribution in Eq. 2.

For each incident plane wave, the light–specimen interaction and the resulting scattering and interference are governed by Maxwell’s equations. The FDTD method solves these equations in the spatial and temporal domains to obtain the full-vector electromagnetic field. In SPECTRAM, the FDTD simulations are performed using the open-source software package Meep [48]. Figure 1d shows an *x* − *z* cross-section of the FDTD computational domain, which consists of source, specimen, and collection regions representing the illumination medium, specimen environment, and objective immersion medium, respectively. Their refractive indices can be independently specified to model configurations involving air, aqueous media, or immersion oil. The computational domain is terminated by perfectly matched layers (PMLs), which absorb outgoing waves and minimize boundary reflections [49]. A planar continuous-wave (CW) source generates monochromatic plane-wave illumination with a prescribed propagation direction and polarization. The simulation proceeds in two stages. During the first stage, the electromagnetic field evolves until transient components decay and a steady periodic response is established, after which the fields exhibit time-harmonic behavior characterized by complex amplitudes [50]. During the second stage, the complex amplitudes on the closed near-field enclosure are calculated and averaged over a fixed time window to reduce numerical errors. The resulting fields are converted into equivalent electric and magnetic surface currents and used in the near-to-far-field transformation to obtain the angular spectrum of light propagating toward the objective [51].

The objective–tube-lens system transforms the angular spectrum into the camera-side spatial-frequency-domain electromagnetic field, accounting for the finite objective aperture, apodization, polarization rotation, and magnification. For each realization with fixed (ω, *θ, ψ*), these fields are then converted into spatial domain via Fourier transform, and the realization-specific image *I*(*x, y*|ω, *θ, ψ*) is calculated from the normal component of the time-averaged Poynting vector, representing the time-averaged energy flux to the camera [52]. As illustrated in Fig. 1e, we analytically establish that it corresponds to the microscope image when the objective is focused at the central axial position *z* = 0 in the FDTD domain. Axial scanning in *z*-stack is generated by applying free-space propagation to each focal position *z*_*f*_ before the Fourier transform. Because distinct source realizations are mutually incoherent in brightfield microscopy, the final *z*-stack is obtained by averaging their image intensities according to the probability distributions:

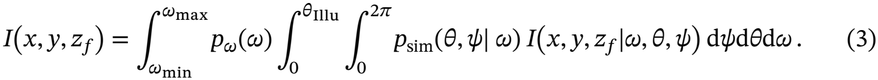

A more detailed theoretical model of brightfield microscopy is presented in Supplementary Section 1.

As illustrated in Fig. 1f, DIC microscopy extends the brightfield optical pathway by introducing a polarizer and a Wollaston prism near the condenser aperture plane [53]. For each incident plane wave, the Wollaston prism generates two orthogonally polarized plane waves with a small angular separation along the shearing direction *γ*. The condenser converts this angular separation into a lateral shear ℓ at the specimen plane, producing two parallel beams that sample positions separated by ℓ. A Nomarski prism and an analyzer near the objective pupil plane subsequently recombine the beams and project their polarizations onto a common direction, enabling interference. A relative bias phase is introduced between the beams, such that their interference converts the differential optical path across the shear distance into intensity contrast, producing the characteristic relief-like DIC appearance. A more detailed model of DIC microscopy is presented in Supplementary Section 2.

Phase-contrast microscopy enhances specimen contrast by modulating the unscattered field relative to the scattered field. As illustrated in Fig. 1g, a condenser annulus restricts the illumination angular spectrum to an annular support. At the conjugate objective pupil plane, a phase ring selectively attenuates and phase-shifts the unscattered field within the corresponding annular region [1]. The attenuation suppresses the background, while interference between the phase-shifted unscattered field and the scattered field converts specimen-induced phase variations into intensity contrast. Compared with the extended source used in brightfield microscopy, the restricted annular source produces greater spatial coherence at the specimen plane, as described by the van Cittert–Zernike relation [54]. Thus, in our system, treating all illumination angles as mutually incoherent was insufficient to reproduce the experimental images. We therefore introduce an effective inter-angle coherence model that combines images obtained in the fully coherent and mutually incoherent limits, with the weighting factor representing the effective degree of mutual coherence among the plane-wave components. A more comprehensive theoretical description of phase-contrast microscopy is provided in Supplementary Section 3.

Figure 2 summarizes the SPECTRAM workflow. Given the specimen geometry, material distributions, source spectrum, and illumination and objective NAs, SPECTRAM samples *N* monochromatic plane-wave realizations from uniform proposal distributions over wavelength and propagation angle. Each realization is simulated once using FDTD and transformed through the objective–tube-lens model to generate a complete *z* -stack. The resulting images are combined using importance weights determined by *p*_sim_(*θ, ψ*) in Eq. 3. This strategy allows different illumination distributions to be evaluated without repeating the FDTD simulations. For DIC microscopy, paired ordinary and extraordinary fields are simulated and coherently recombined, whereas phase-contrast microscopy incorporates annular illumination, pupil-plane amplitude and phase modulation, and the effective inter-angle coherence model.

**Fig 2.**
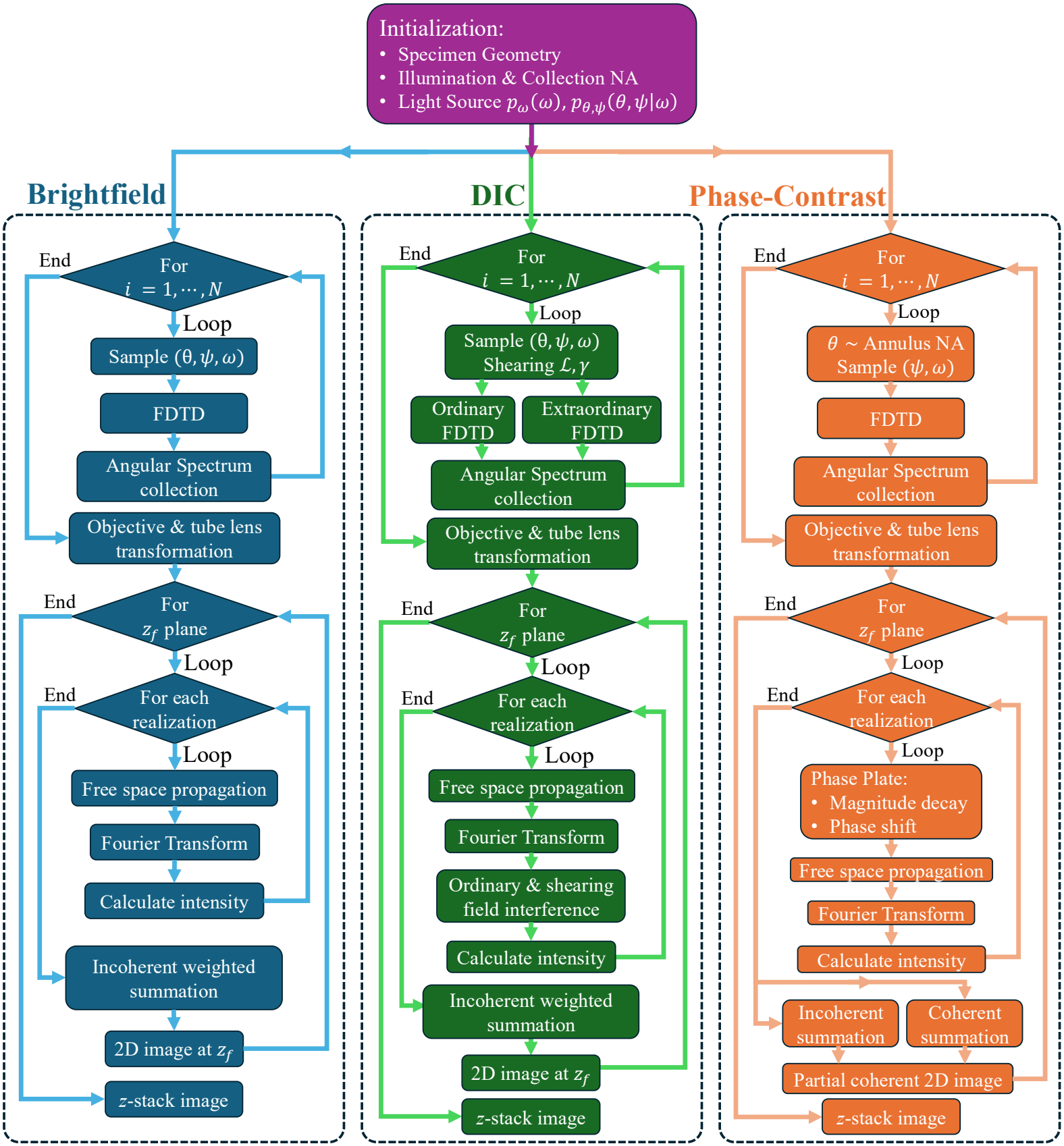
SPECTRAM workflows for brightfield, DIC, and phase-contrast *z*-stack simulation. For each spectral–angular plane-wave realization, the FDTD-derived electromagnetic field is transformed through the objective–tube-lens system and propagated to the target focal positions. Brightfield images are formed by importance-weighted intensity averaging; DIC additionally recombines the ordinary and extraordinary fields, whereas phase contrast incorporates annular illumination, phase-plate modulation, and effective inter-angle coherence.

## 3. Results

After establishing SPECTRAM as described above, we first validate its accuracy in 3D space by comparing its results with those obtained from existing theoretical and experimental models. For theoretical validation, Mie scattering theory provides an analytical solution for the field scattered by a homogeneous sphere illuminated by a polarized plane wave [55]. Note that this scattered field is evaluated on the sample side, before being collected and transformed by the objective and tube lens. In Supplementary Section 4, we demonstrate that the scattered field calculated by SPECTRAM under this configuration agrees closely with the Mie-theory solution, validating the accuracy of its light-sample interaction simulation.

For experimental validation, we acquired 3D *z*-stack images of 1 µm diameter polystyrene beads, whose refractive index is 1.6 at a wavelength 0.516 µm, using the Zeiss Axio Observer 7 microscope at different illumination NAs in brightfield, DIC, and phase-contrast modes. The polystyrene beads were placed in a specimen chamber formed by a glass slide and a coverslip, with deionized water serving as the sample medium (*n*_medium_ = 1.33). SPECTRAM subsequently simulated 3D *z*-stack images of the beads under the corresponding configurations, and the simulated results were compared with the experimental measurements to further validate its modeling of the complete microscope system, as shown in Section 3.1, 3.2, and 3.3. Finally, in Section 3.4, we used SPECTRAM to simulate *z*-stack images of a thick cellular sample within a brain organoid. This example demonstrates that SPECTRAM is applicable not only to geometrically regular objects but can also be extended to samples with arbitrary structures.

### 3.1 Simulation of Brightfield Microscopy

We first compared the brightfield images simulated by SPECTRAM with experimentally acquired 3D *z*-stacks of 1µm-diameter polystyrene beads. During the experimental acquisition, the illumination NA was precisely controlled between 0.09 and 0.55. A 63 × oil-immersion objective with an NA of 1.4 was used, together with immersion oil having a refractive index of 1.515. *z* -stack was acquired by stacking 2-dimensional camera images acquired with the objective at different axial focal positions. The same configuration parameters were incorporated into SPECTRAM to simulate the corresponding *z* -stack images. Figure 3 compares the experimental and simulated brightfield images over illumination NAs ranging from 0.09 to 0.55 using *xy*- and *xz*-plane cross-sections, together with normalized intensity profiles along the *x*- and *z*-directions.

**Fig 3.**
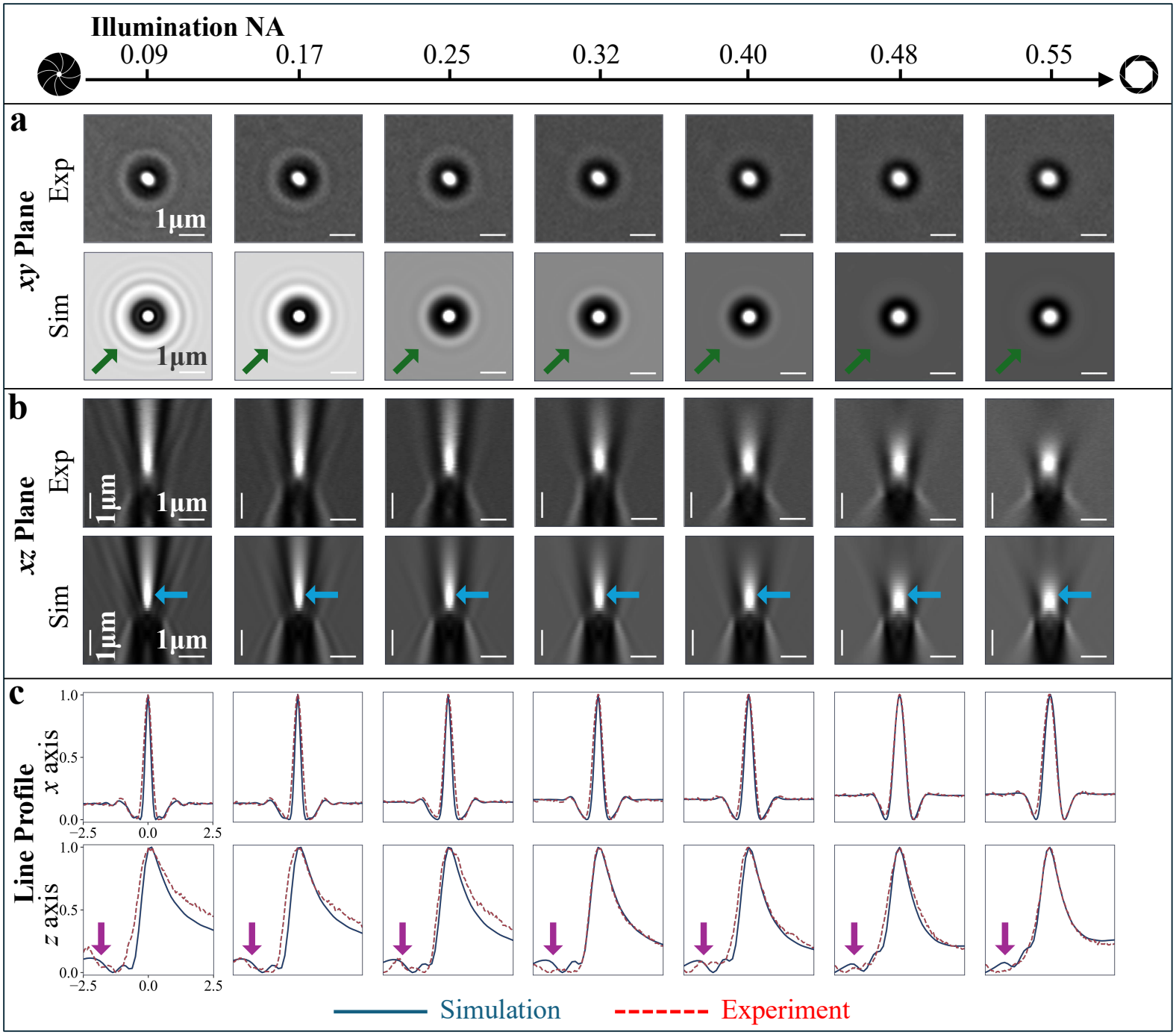
Comparison of experimental and simulated brightfield images of 1µm polystyrene beads at different illumination NAs. **a**, comparison of *xy*-plane cross-sectional images. **b**, comparison of *xz*-plane cross-sectional images. **c**, normalized intensity profiles along the *x*- and *z*-directions.

In the *xy*-plane images, both the experimental and simulated results exhibit a central bright spot surrounded by a dark ring and progressively decaying outer fringes (indicated by the green arrows). As the illumination NA increases, these outer fringes become less pronounced, evolving from multiple concentric rings at 0.09 to an almost completely uniform background at 0.55. This evolution is consistently reproduced in both the experimental and simulated images. The *xz*-plane images further reveal the characteristic asymmetric axial diffraction pattern. At low illumination NAs, the central bright region produced by the bead is narrow and elongated along the axial direction, accompanied by pronounced fringes (indicated by the blue arrows). As the illumination NA increases, its axial extent decreases while it becomes laterally broader around the focal region, and the surrounding fringes gradually disappear. SPECTRAM consistently reproduces this evolution observed in the experimental images.

The normalized line profiles along the *x*- and *z*-directions further demonstrate the close agreement between the experimental and simulated results. Along the *x*-direction, the two profiles overlap closely at both the central intensity peak and the adjacent dark minima. Along the *z*-direction, although some discrepancies are observed in the negative *z* region (indicated by the purple arrows), both profiles reproduce the same overall trend of an initial increase followed by a gradual decrease. The discrepancies in the negative *z* region may arise because the glass slide present in the experimental specimen was not explicitly modeled in the simulation.

Both the cross-sectional images and the normalized line profiles reveal a trade-off associated with increasing illumination NA: the lateral image contrast decreases, whereas the axial sectioning capability improves [56,57]. This behavior is consistently reproduced by SPECTRAM and can be understood from the increased angular diversity of the illumination. At higher illumination NAs, more obliquely incident plane-wave components contribute to image formation. Near the focal plane, their direction-dependent diffraction patterns spread the bright signal over a broader lateral region, thereby reducing the peak-to-background contrast. Away from the focal plane, these patterns undergo larger direction-dependent lateral displacements and no longer remain spatially aligned. Their incoherent averaging therefore suppresses the extended axial fringes and out-of-focus contrast, resulting in a signal that is more strongly localized around the focal plane and hence improved axial sectioning.

In addition, the results of SPECTRAM exhibit an initial defocus of approximately +0.6µm along the axial direction, indicating that the strongest image contrast occurs in a plane above the bead rather than at the bead itself. More fundamentally, a brightfield microscope does not directly image the physical sample; instead, it records the diffraction pattern produced by the sample and the interference between scattering and incident light.

### 3.2 Simulation of DIC Microscopy

Starting from the same optical configuration used for bright-field imaging, experimental DIC images of the 1µm diameter polystyrene beads were acquired by introducing a polarizer, a Wollaston prism, a Nomarski prism, and an analyzer into the optical path, as shown in Fig. 1f. For the 63 ×, 1.4-NA objective used here, a Zeiss DIC III prism was required. Calibration showed that the two orthogonally polarized beams were sheared along a direction oriented at 45^°^ relative to the horizontal axis, with a lateral shear distance of approximately 0.175 µm in the sample plane.

The same parameters were incorporated into SPECTRAM to simulate the corresponding *z*-stack images at different illumination NAs. Figure 4 compares the experimental and simulated results using cross-sectional images in different planes. The selected planes include the *xy*-plane, the vertical plane spanned by the shear direction *y* = *x* (shown in orange) and the *z*-axis, and the vertical plane spanned by the direction *y* = −*x* (shown in green), which is perpendicular to the shear direction, and the *z*-axis.

**Fig 4.**
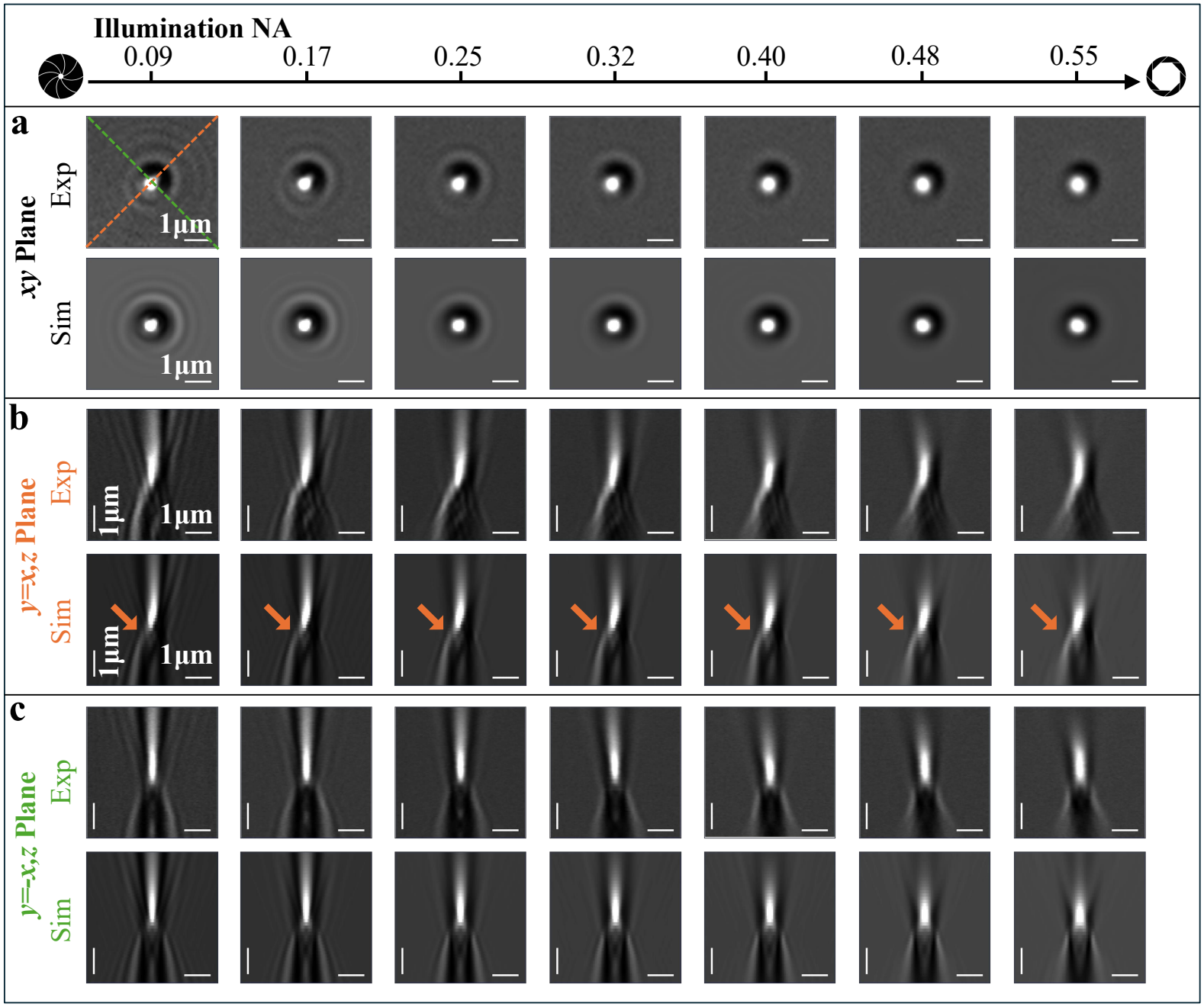
Comparison of experimental and simulated DIC images of 1µm polystyrene beads at different illumination NAs. **a**, *xy*-plane cross-sectional images. **b**, (*y* = *x, z*)-plane cross-sectional images. **c**, (*y* = −*x, z*)-plane cross-sectional images.

In the *xy* -plane, the DIC images no longer exhibit the rotationally symmetric pattern observed in brightfield microscopy. Instead, they display a characteristic bright-dark contrast pair along the shear direction, with one side appearing bright and the opposite side appearing dark. This asymmetry primarily arises because the two orthogonally polarized and laterally sheared beams produced by the Wollaston-Nomarski prism system are projected onto the same polarization direction by the analyzer, allowing them to interfere. The two beams are laterally displaced along the shear direction. Their relative phase bias, determined through SPECTRAM to be approximately −0.5π, converts the differential phase between the two optical paths into the characteristic asymmetric bright-dark intensity pattern. Similar to the brightfield results, the surrounding fringes become progressively weaker and the central bright region broadens as the illumination NA increases.

In the vertical (*y* = *x, z*) plane, this asymmetry becomes even more pronounced along the axial direction. In the negative *z* region, the bright feature bends toward the left (shown at the orange arrow), whereas in the positive *z* region, it remains approximately aligned with the axial direction. In SPECTRAM, the leftward bending can be understood as a consequence of the phase and amplitude perturbations introduced by the bead, which redistribute the transmitted and scattered fields around the bead and shift the region of constructive interference away from the optical axis. Under the −0.5π phase bias, constructive interference occurs predominantly on the left side, producing the observed asymmetric curvature. In the vertical (*y* = −*x, z*) plane, which is perpendicular to the shear direction, both the experimental and simulated images recover an approximately symmetric pattern similar to that observed in brightfield microscopy. Moreover, as in brightfield microscopy, both the (*y* = *x, z*) and (*y* = −*x, z*) planes exhibit the trade-off between reduced image contrast and improved axial sectioning as the illumination NA increases.

In Supplementary Section 2E, we further validate the accuracy of SPECTRAM simulations by comparing simulated and experimentally measured line profiles.

### 3.3 Simulation of Phase-Contrast Microscopy

Compared with bright-field microscopy, phase-contrast microscopy requires an annulus in the condenser aperture plane to produce annular illumination, together with a specialized objective containing a phase plate. To acquire the experimental phase-contrast *z*-stacks, we used a Zeiss Phase 2 condenser annulus and a 40 ×, 0.75-NA dry objective. The illumination NA range produced by the Phase 2 annulus was measured to be 0.25–0.30. In SPECTRAM, the phase plate of the objective was modeled as applying an amplitude attenuation factor of 0.5 and a phase shift of −0.3π to the corresponding annular region in the spatial-frequency domain, thereby suppressing the unscattered incident light and enhancing image contrast. Compared with the brightfield and DIC results, the experimental phase-contrast *z*-stacks retained visible contrast over a substantially larger axial range, approximately 100 µm.

To first validate the modeling of the condenser annulus, we acquired brightfield images using a conventional objective under annular illumination produced by different condenser annuli. These experimental images showed close agreement with the corresponding SPECTRAM simulations. The comparison also explains why annular illumination produces image contrast over a larger axial range, as discussed in detail in Supplementary Section 3A.

Using the same parameters, SPECTRAM simulated the phase-contrast images under the Zeiss Phase 2 configuration. Figure 5 illustrates the simulation procedure and compares the simulated results with the corresponding experimental images. As described in the Methods, the spatial-frequency support of the incident illumination in phase-contrast microscopy is confined to a narrow annular region. Consequently, the plane-wave components exhibit a higher degree of mutual spatial coherence. SPECTRAM models this inter-angle partial coherence by separately calculating the images under the fully incoherent and fully coherent limits and then combining them through a weighted sum [58]. The fully incoherent and fully coherent components contribute 97% and 3%, respectively, to the final image intensity.

**Fig 5.**
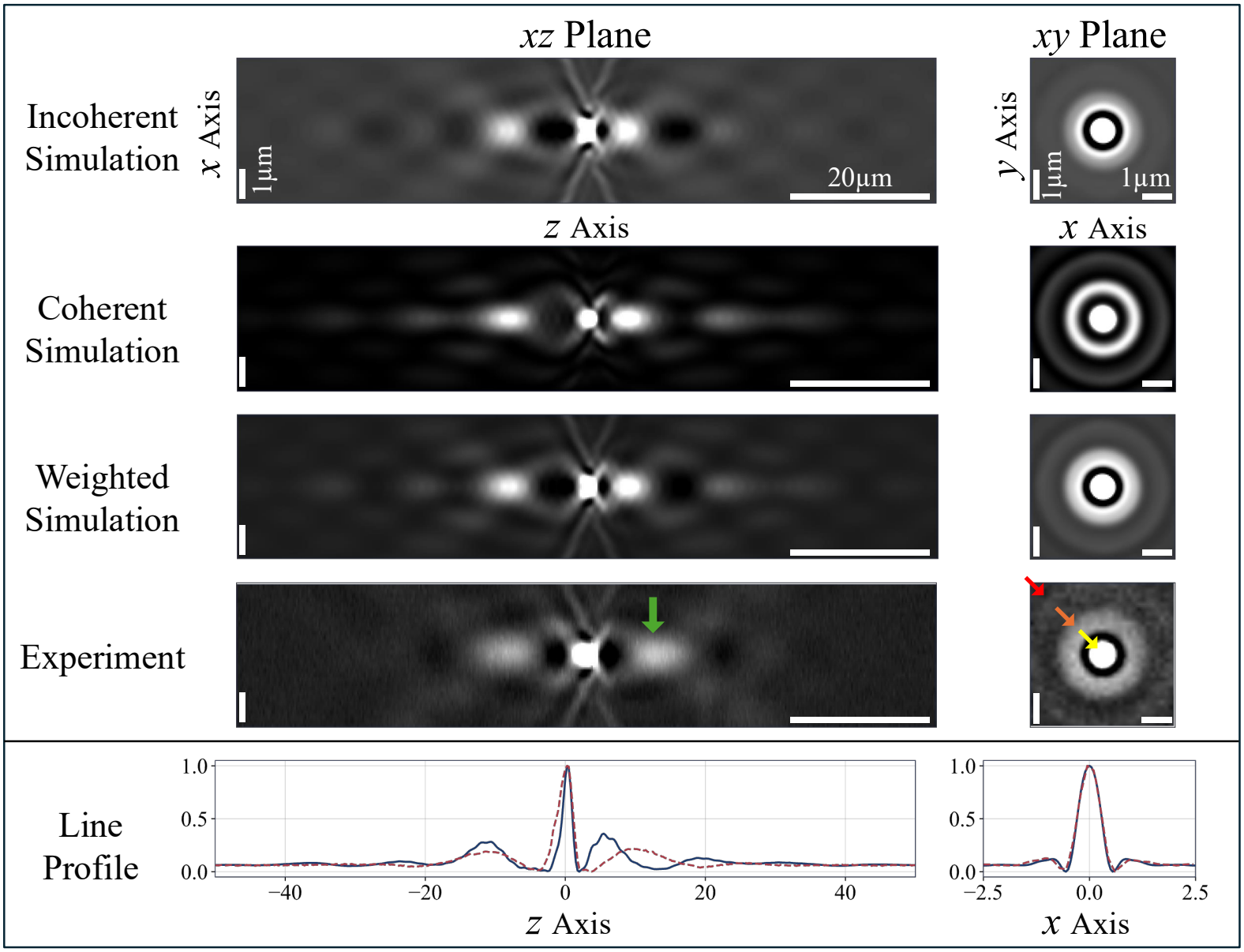
Comparison of experimental and simulated phase-contrast images of 1µm polystyrene beads. The left and right columns show the *xz*- and *xy*-plane images, respectively. From top to bottom, the rows present the fully incoherent simulation, fully coherent simulation, weighted simulation, experimental measurement, and normalized line-profile comparison.

As shown in Fig. 5, incoherent averaging produces a stronger background and more spatially extended bright-fringe features. In contrast, coherent summation produces stronger interference, resulting in more pronounced concentric fringes in the *xy*-plane and a weaker background. After the two components are combined using the prescribed weights, the resulting images show good agreement with the experimental measurements. In the *xy* -plane, the simulated halo effect closely matches the experimental observation, with the central disk surrounded by two outer bright rings (indicated by the yellow, orange and red arrows) [59]. Along the axial direction, the alternating distribution of bright and dark regions is also consistently reproduced. This agreement is further supported by the line profiles along the *x*- and *z*-directions. The primary discrepancy is that the second bright region (indicated by the green arrow) on the positive *z* side appears closer to the focal plane in the simulation than in the experiment. A potential explanation for this discrepancy is that SPECTRAM does not explicitly model the glass slide. In the experiment, the second bright region lies within the glass slide, where refraction at the medium–glass interface may shift its appearance to a greater axial depth than predicted for the homogeneous medium assumed in the simulation.

### 3.4 3D imaging of thick cells

Using 1µm diameter polystyrene beads as a standard sample, we validated SPECTRAM’s ability to simulate brightfield, DIC, and phase-contrast imaging by comparison with experimental measurements. SPECTRAM is not restricted to samples with regular geometries and can be applied to arbitrary sample morphologies. Here, we further demonstrate its capability by simulating the *z*-stack images of an iPSC-derived stem cell.

The primary challenge in simulating the cell is that its true refractive-index distribution is unknown. As shown in Fig. 6, we first used gradient light interference microscopy (GLIM) with a 40× oil-immersion objective to acquire optical-path-induced phase maps throughout the 3D volume for reconstructing this distribution [60]. Subsequently, under the approximation that the measured phase difference depends linearly on the refractive-index contrast, we reconstructed the refractive-index distribution of the cell [61]. The detailed reconstruction algorithm is provided in Supplement 6. The reconstructed cell volume had dimensions of 25µm × 25µm × 20µm and exhibited a complex 3D refractive-index distribution. The reconstructed cellular structure was incorporated into SPECTRAM to simulate its *z*-stack images under brightfield, DIC, and phase-contrast modalities, which were subsequently compared with the corresponding experimental measurements. Each image stack covered an axial range of 50 µm, corresponding to approximately 100 optical wavelengths. The brightfield configuration used an illumination NA of 0.09 and a 40 ×, 1.4-NA oil-immersion objective. The DIC configuration used the same 40x, 1.4 NA objective together with the Zeiss DIC II mode, with a shear angle of 45^°^ and a shear distance of 0.375 µm. The phase-contrast configuration used the Zeiss Phase 2 mode (illumination NA range 0.25-0.30) and a 40 ×, 0.75-NA dry objective.

**Fig 6.**
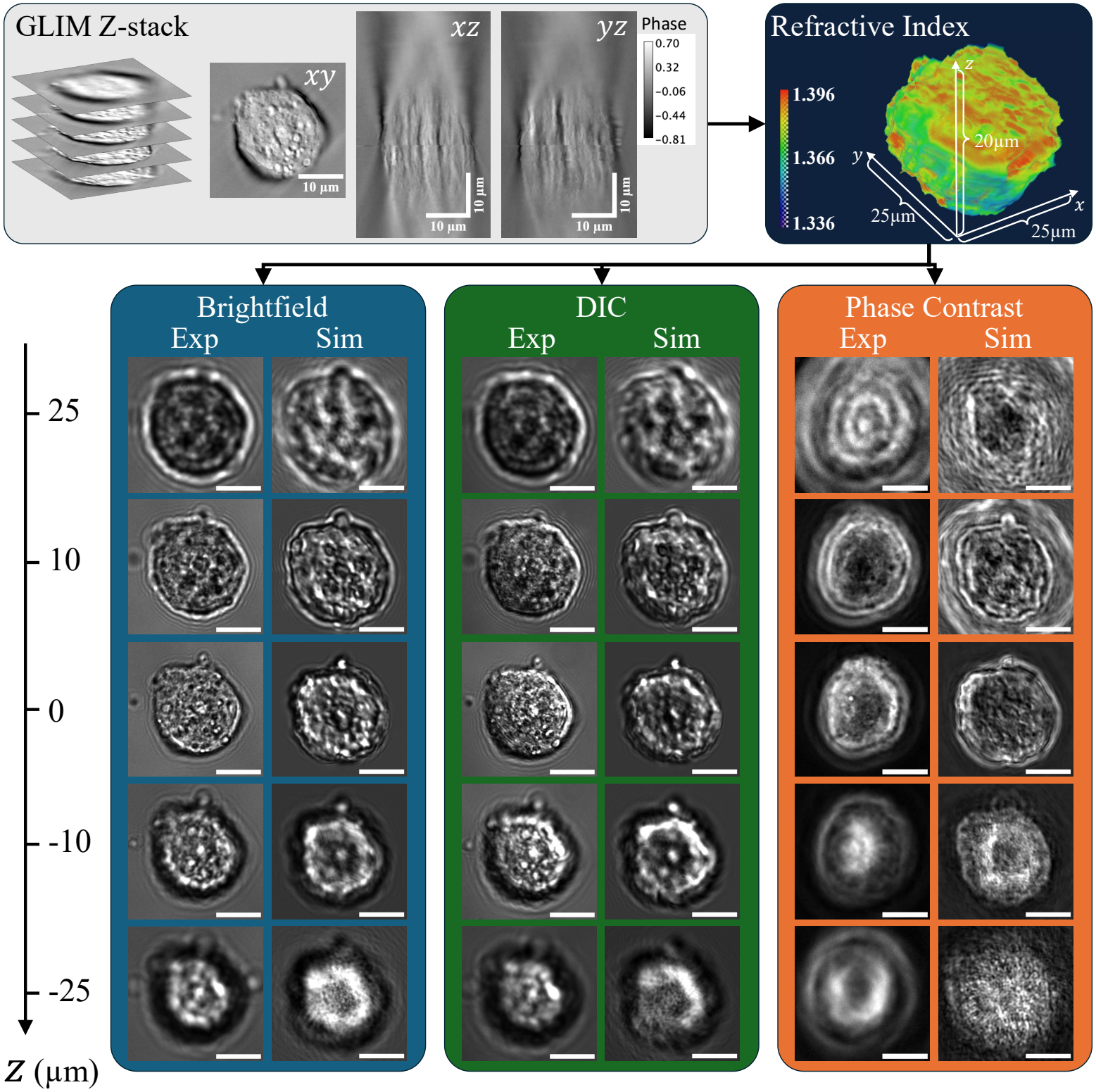
Workflow for simulating a thick cellular structure. A *z*-stack of the cell was first acquired using GLIM, from which its three-dimensional refractive-index distribution was reconstructed computationally. The reconstructed structure was then incorporated into SPECTRAM to simulate 50µm long *z* -stacks under brightfield, DIC, and phase-contrast imaging. Comparison with the corresponding experimental measurements shows close agreement across all three modalities.

Figure 6 compares the experimental and simulated two-dimensional images at different focal positions spanning an axial range from -25 to 25µm, where *z* = 0 corresponds to the focal position exhibiting the highest image contrast. For the brightfield images at the *z* = 0 plane, SPECTRAM successfully reproduces the experimentally observed outer contour of the cell and the alternating bright and dark features within its internal structure. As the focal plane moves in the positive *z* direction, the apparent cellular contour expands outward, and a pronounced bright ring emerges along the boundary. In contrast, as the focal plane moves toward negative *z*, the apparent contour contracts, a dark ring forms around the boundary, and the internal structures become increasingly blurred. This variation with focal position is consistently reproduced by SPECTRAM. Although the detailed cellular morphology differs somewhat from the experimental observations, this discrepancy primarily arises from uncertainties in reconstructing the cell’s refractive-index distribution. Nevertheless, SPECTRAM correctly captures the overall 3D evolution of the image pattern produced by the cellular structure.

In the DIC mode, the most prominent change is that the image intensity is no longer distributed symmetrically across the cell. Instead, a directional bright–dark contrast appears along the 45^°^ shear direction, with lower intensity toward the lower-left side and higher intensity toward the upper-right side. This directional asymmetry is particularly pronounced along the cellular boundary at all focal positions. This behavior primarily arises from the spatially nonuniform interference pattern generated by the two laterally sheared beams in the DIC system. SPECTRAM reproduces this characteristic pattern observed in the experimental images, demonstrating its ability to accurately simulate the *z*-stack features of a cellular sample under DIC imaging.

In phase-contrast imaging, the background is strongly suppressed, enhancing the cellular contrast and giving the internal structures a characteristic translucent appearance. These features are consistently observed in both the experimental and simulated images at the *z* = 0 focal plane. As the focal plane is shifted in either the positive- or negative-*z* direction, both the experimental and simulated images exhibit successive ring-shaped contours that expand outward or contract inward. This behavior mainly arises because the annular illumination causes the diffraction pattern generated by the sample to spread radially in a ring-like manner as it propagates away from the focal plane. However, the experimental and simulated phase-contrast images exhibit substantial differences in their detailed cellular morphology. This discrepancy primarily arises from the need to use different objectives for the two measurements. The GLIM data were acquired using a high-NA oil-immersion objective to improve the accuracy of refractive-index reconstruction. In contrast, phase-contrast imaging required a different objective, necessitating specimen repositioning. The resulting specimen handling induced morphological deformation of the cell and lack of ground truth for the refractive-index distribution of the cell, contributing to the observed discrepancy.

Taken together, these results demonstrate that SPECTRAM can accurately reproduce the 3D image features of complex biological samples across bright-field, DIC, and phase-contrast imaging modalities.

## 4. Discussions

In this work, we developed SPECTRAM, a comprehensive physics-based framework for simulating 3D image formation in phase-sensitive microscopy. SPECTRAM integrates vectorial electromagnetic modeling, polarization, and tunable spatial and temporal coherence into a unified framework that describes the complete image-formation process. Specifically, broadband, partially coherent illumination is decomposed into monochromatic plane-wave components, with their angular and wavelength distributions characterized by probability density functions. The interactions between the illumination field and the sample, as well as the resulting diffraction fields, are calculated by solving the full-vector Maxwell equations. The model further incorporates the signal-collection optics, including the objective and tube lens, and the physical mechanisms underlying image formation and image-stack acquisition. By accounting for these processes, SPECTRAM provides a unified approach to simulating brightfield, DIC, and phase-contrast microscopy, while allowing the optical and illumination parameters to be adapted to different microscopy configurations.

The accuracy of SPECTRAM was validated at multiple levels. First, the calculated scattering field produced by a spherical bead agreed closely with Mie scattering theory, validating the electromagnetic field calculation. Second, comparisons with experimental measurements using specimens containing 1-µm-diameter polystyrene beads demonstrated that SPECTRAM reproduces modality-specific, illumination-NA-dependent, and defocus-dependent image features. Importantly, this agreement extends throughout the axial dimension, enabling the reproduction of experimentally observed image evolution across 3D image stacks rather than only at selected focal planes. Such validation of continuous axial image formation is particularly important because image characteristics in phase-sensitive microscopy can change substantially with focal position, and accurate prediction of these changes is essential for understanding image contrast and phase formation in 3D specimens. Furthermore, applying SPECTRAM to a complex cellular structure produced simulated image stacks that closely matched experimental observations. This demonstrates that the framework is not restricted to idealized samples or regular geometries but can reproduce image formation in morphologically complex biological specimens.

Beyond validating the simulation framework, our results reveal an important distinction between the roles of angular and wavelength spectra in microscopy image formation. The angular spectrum describes the diversity of incident wave vectors, corresponding to different illumination directions, whereas the wavelength spectrum describes the distribution of optical wavelengths. Our simulations demonstrate that angular diversity can have a substantially greater influence on reproducing the experimentally observed image characteristics than wavelength diversity. Specifically, simulations using a single wavelength with an appropriately distributed angular spectrum closely reproduced the experimental images, whereas incorporating wavelength diversity without the corresponding angular diversity did not achieve comparable agreement. This finding highlights the importance of accurately characterizing the angular distribution of illumination when modeling partially coherent microscopy systems. It also suggests that accounting for spectral bandwidth alone is insufficient to capture the full effects of partial coherence on image formation. More broadly, these results emphasize that the spatial and temporal coherence properties of illumination contribute differently to image formation and both should be considered when developing accurate physical models of microscopy systems.

The comprehensive physical modeling capability of SPECTRAM provides opportunities beyond reproducing experimental images. By modifying the illumination distributions, polarization states, optical configurations, and image-acquisition parameters, the framework can be adapted to investigate a broad range of phase-sensitive microscopy modalities and configurations. It can serve as a computational tool for systematically investigating how individual optical parameters and their interactions affect image contrast, phase evolution with focal position, axial image characteristics, and phase sensitivity. For example, simulations can help identify the physical origins of contrast variations and imaging artifacts, determine how illumination NA and coherence properties influence the formation and evolution of phase contrast, and evaluate the conditions under which different imaging modalities provide reliable structural information. For thick and highly scattering specimens, the framework may also help investigate the factors governing imaging depth and the loss of image contrast with increasing depth. These capabilities can provide theoretical guidance for identifying the operating regimes and limitations of existing imaging methods, optimizing microscopy configurations for specific samples, and developing new imaging modalities.

Despite these capabilities, the current implementation of SPECTRAM focuses on transmitted-light wide-field imaging systems, including brightfield, DIC, and phase-contrast microscopy. Extending the framework to other imaging configurations, particularly epi-illumination and laser-scanning systems, will further expand its applicability. Such extensions will require incorporating their corresponding illumination and signal-detection mechanisms, including scanning processes and optical sectioning. In particular, modeling configurations designed to improve axial sectioning in thick specimens will provide opportunities to investigate the interplay between light propagation, scattering, and image formation in more complex imaging systems. Overall, SPECTRAM establishes a physics-based foundation for comprehensive 3D microscopy simulation, enabling quantitative investigation of image formation in complex samples and providing a versatile computational platform for understanding, evaluating, and advancing phase-sensitive microscopy.

## Acknowledgements

We acknowledge the support of the National Institute of Biomedical Imaging and Bioengineering (grant nos. R00EB034164 and K99EB034164 to Xi Chen).

## Author Contributions

X.C. proposed the idea and conceived the project. J.Y. and X.C. developed the computational algorithms and theory. J.Y. conducted the imaging and analyzed the data. X.C. and J.Y. wrote the manuscript. X.C. supervised the project.

## Data Availability

Due to size considerations, the data that support the findings of this study are available from the corresponding author upon reasonable request.

## Corresponding author

Correspondence to X.C.

## References

1. F. Zernike, “Phase contrast, a new method for the microscopic observation of transparent objects,” Physica 9, 686–698 (1942).

2. G. Nomarski, “Microinterféromètre différentiel à ondes polarisées,” J. Phys. Radium 16, 9S–13S (1955).

3. X. Chen, M. E. Kandel, and G. Popescu, “Spatial light interference microscopy: principle and applications to biomedicine,” Adv. Opt. Photon., AOP 13, 353–425 (2021).

4. X. Chen, J. Li, and O. Korotkova, “Light scintillation in soft biological tissues,” Waves in Random and Complex Media 30, 481–489 (2020).

5. J. Li, X. Chen, S. McDuffie, M. A. M. Najjar, S. M. H. Rafsanjani, and O. Korotkova, “Mitigation of atmospheric turbulence with random light carrying OAM,” Optics Communications 446, 178–185 (2019).

6. X. Chen and O. Korotkova, “Optical beam propagation in soft anisotropic biological tissues,” OSA Continuum, OSAC 1, 1055–1067 (2018).

7. X. Chen and O. Korotkova, “Scattering of light from hollow and semi-hollow 3D scatterers with ellipsoidal, cylindrical and cartesian symmetries,” КомпЬЮтеРная оптика 40, 635–641 (2016).

8. X. Chen and O. Korotkova, “Probability density functions of instantaneous Stokes parameters on weak scattering,” Optics Communications 400, 1–8 (2017).

9. M. E. Kandel, M. Fanous, C. Best-Popescu, and G. Popescu, “Real-time halo correction in phase contrast imaging,” Biomed. Opt. Express, BOE 9, 623–635 (2018).

10. T. H. Nguyen, M. Kandel, H. M. Shakir, C. Best-Popescu, J. Arikkath, M. N. Do, and G. Popescu, “Halo-free Phase Contrast Microscopy,” Sci Rep 7, 44034 (2017).

11. M. Chen, L. Tian, and L. Waller, “3D differential phase contrast microscopy,” Biomed. Opt. Express, BOE 7, 3940–3950 (2016).

12. B. Gutiérrez-Medina, “Making sense of blobs, whorls, and shades: methods for label-free, inverse imaging in bright-field optical microscopy,” Biophys Rev 17, 335–345 (2025).

13. X. Chen, M. E. Kandel, S. He, C. Hu, Y. J. Lee, K. Sullivan, G. Tracy, H. J. Chung, H. J. Kong, M. Anastasio, and G. Popescu, “Artificial confocal microscopy for deep label-free imaging,” Nat. Photon. 17, 250–258 (2023).

14. X. Chen, M. E. Kandel, S. Zhao, R. T. Zirkel, K.-Y. Huang, H. J. Kong, C. B. Schaffer, and C. Xu, “Scattering-enabled epi-quantitative phase imaging reveals subcellular detail in organoids and deep mouse brains,” (2026).

15. J. Yao, J. Lyu, N. Shi, and X. Chen, “Intelligent SEEQPI enables reconstruction of inhibitory neuronal nuclei and dry-mass mapping deep in the living mouse brain,” 2026.10.25.761694 (2026).

16. A. Badon, A. C. Boccara, G. Lerosey, M. Fink, and A. Aubry, “Multiple scattering limit in optical microscopy,” Opt. Express, OE 25, 28914–28934 (2017).

17. N. Goswami, N. Winston, W. Choi, N. Z. E. Lai, R. B. Arcanjo, X. Chen, N. Sobh, R. A. Nowak, M. A. Anastasio, and G. Popescu, “EVATOM: an optical, label-free, machine learning assisted embryo health assessment tool,” Commun Biol 7, 268 (2024).

18. C. W. McCutchen, “Generalized Aperture and the Three-Dimensional Diffraction Image,” J. Opt. Soc. Am., JOSA 54, 240–244 (1964).

19. N. Streibl, “Three-dimensional imaging by a microscope,” J. Opt. Soc. Am. A, JOSAA 2, 121–127 (1985).

20. X. Chen and O. Korotkova, “Complex degree of coherence modeling with famous planar curves,” Opt. Lett., OL 43, 6049–6052 (2018).

21. O. Korotkova, X. Chen, and T. Setälä, “Electromagnetic Schell-model beams with arbitrary complex correlation states,” Opt. Lett., OL 44, 4945–4948 (2019).

22. X. Chen and O. Korotkova, “Phase structuring of 2D complex coherence states,” Opt. Lett., OL 44, 2470–2473 (2019).

23. O. Korotkova and X. Chen, “Phase structuring of the complex degree of coherence,” Opt. Lett., OL 43, 4727–4730 (2018).

24. E. Wolf, “Three-dimensional structure determination of semi-transparent objects from holographic data,” Optics Communications 1, 153–156 (1969).

25. X. Chen, M. E. Kandel, C. Hu, Y. J. Lee, and G. Popescu, “Wolf phase tomography (WPT) of transparent structures using partially coherent illumination,” Light Sci Appl 9, 142 (2020).

26. Y. Guo, Z. Zhuang, T. Chen, and M. Hu, “Image reconstruction algorithms for optical diffraction tomography microscopy: A review,” Optics and Lasers in Engineering 194, 109163 (2025).

27. P. R. T. Munro and P. Török, “Vectorial, high-numerical-aperture study of phase-contrast microscopes,” J. Opt. Soc. Am. A, JOSAA 21, 1714–1723 (2004).

28. H. Sierra, C. A. DiMarzio, and D. H. Brooks, “Modeling phase microscopy of transparent three-dimensional objects: a product-of-convolutions approach,” J. Opt. Soc. Am. A, JOSAA 26, 1268–1276 (2009).

29. H.-Y. Liu, D. Liu, H. Mansour, P. T. Boufounos, L. Waller, and U. S. Kamilov, “SEAGLE: Sparsity-Driven Image Reconstruction Under Multiple Scattering,” IEEE Transactions on Computational Imaging 4, 73–86 (2018).

30. L. Tian and L. Waller, “3D intensity and phase imaging from light field measurements in an LED array microscope,” Optica, OPTICA 2, 104–111 (2015).

31. M. Chen, D. Ren, H.-Y. Liu, S. Chowdhury, and L. Waller, “Multi-layer Born multiple-scattering model for 3D phase microscopy,” Optica, OPTICA 7, 394–403 (2020).

32. T. Pham, E. Soubies, A. Ayoub, J. Lim, D. Psaltis, and M. Unser, “Three-Dimensional Optical Diffraction Tomography With Lippmann-Schwinger Model,” IEEE Transactions on Computational Imaging 6, 727–738 (2020).

33. L. Vinckenbosch, C. Lacaux, S. Tindel, M. Thomassin, and T. Obara, “Monte Carlo methods for light propagation in biological tissues,” Mathematical Biosciences 269, 48–60 (2015).

34. F. Cai and S. He, “Electric field Monte Carlo simulation of focused stimulated emission depletion beam, radially and azimuthally polarized beams for in vivo deep bioimaging,” JBO 19, 011022 (2014).

35. A. Taflove, S. C. Hagness, and M. Piket-May, “9 -Computational Electromagnetics: The Finite-Difference Time-Domain Method,” in The Electrical Engineering Handbook, W.-K. Chen, ed. (Academic Press, 2005), pp. 629–670.

36. F. L. Teixeira, C. Sarris, Y. Zhang, D.-Y. Na, J.-P. Berenger, Y. Su, M. Okoniewski, W. C. Chew, V. Backman, and J. J. Simpson, “Finite-difference time-domain methods,” Nat Rev Methods Primers 3, 75 (2023).

37. Q. H. Liu, “The PSTD algorithm: A time-domain method requiring only two cells per wavelength,” Microwave and Optical Technology Letters 15, 158–165 (1997).

38. X. Wei, A. J. Wachters, and H. P. Urbach, “Finite-element model for three-dimensional optical scattering problems,” J. Opt. Soc. Am. A, JOSAA 24, 866–881 (2007).

39. İ. R. Çapoğlu, C. A. White, J. D. Rogers, H. Subramanian, A. Taflove, and V. Backman, “Numerical simulation of partially coherent broadband optical imaging using the finite-difference time-domain method,” Opt. Lett., OL 36, 1596–1598 (2011).

40. İ. R. Çapoğlu, J. D. Rogers, A. Taflove, and V. Backman, “The Microscope in a Computer: Image Synthesis from Three-Dimensional Full-Vector Solutions of Maxwell’s Equations at the Nanometer Scale,” in Progress in Optics (Elsevier, 2012), Vol. 57, pp. 1–91.

41. D. Zhang, I. Capoglu, Y. Li, L. Cherkezyan, J. Chandler, G. Spicer, H. Subramanian, A. Taflove, and V. Backman, “Finite-difference time-domain-based optical microscopy simulation of dispersive media facilitates the development of optical imaging techniques,” JBO 21, 065004 (2016).

42. S. Tanev, J. Pond, P. Paddon, and V. V. Tuchin, “A Finite-Difference Time-Domain Model of Optical Phase Contrast Microscope Imaging,” in Optical Waveguide Sensing and Imaging, W. J. Bock, I. Gannot, and S. Tanev, eds. (Springer Netherlands, 2008), pp. 243–257.

43. P. Choo, A. J. Hryn, K. S. Culver, D. Bhowmik, J. Hu, and T. W. Odom, “Wavelength-Dependent Differential Interference Contrast Inversion of Anisotropic Gold Nanoparticles,” J. Phys. Chem. C 122, 27024–27031 (2018).

44. Y. Suzuki, S. Watanabe, M. Odaira, T. Okamura, and Y. Kawata, “Numerical simulations of partially coherent illumination for multi-scattering phase objects via a beam propagation method,” Appl. Opt. 58, 954 (2019).

45. J. L. Hollmann, A. K. Dunn, and C. A. DiMarzio, “Computational microscopy in embryo imaging,” Opt. Lett. 29, 2267 (2004).

46. S. B. Mehta and R. Oldenbourg, “Image simulation for biological microscopy: microlith,” Biomed. Opt. Express, BOE 5, 1822–1838 (2014).

47. J. W. Goodman, Introduction to Fourier Optics (American Institute of Physics, 1969).

48. A. F. Oskooi, D. Roundy, M. Ibanescu, P. Bermel, J. D. Joannopoulos, and S. G. Johnson, “Meep: A flexible free-software package for electromagnetic simulations by the FDTD method,” Computer Physics Communications 181, 687–702 (2010).

49. J.-P. Berenger, “A perfectly matched layer for the absorption of electromagnetic waves,” Journal of Computational Physics 114, 185–200 (1994).

50. A. Taflove, “Application of the Finite-Difference Time-Domain Method to Sinusoidal Steady-State Electromagnetic-Penetration Problems,” IEEE Transactions on Electromagnetic Compatibility EMC-22, 191–202 (1980).

51. R. C. Bollimuntha, M. F. Hadi, M. J. Piket-May, and A. Z. Elsherbeni, “Near-to-Far Field Transformation in FDTD: A Comparative Study of Different Interpolation Approaches,” Applied Computational Electromagnetics Society Journal (ACES) 496–504 (2021).

52. M. Born and E. Wolf, Principles of Optics: Electromagnetic Theory of Propagation, Interference and Diffraction of Light, 7th ed. (Cambridge University Press, 1999).

53. S. B. Mehta and C. J. R. Sheppard, “Partially coherent image formation in differential interference contrast (DIC) microscope,” Opt. Express, OE 16, 19462–19479 (2008).

54. C. W. McCutchen, “Generalized Source and the van Cittert–Zernike Theorem: A Study of the Spatial Coherence Required for Interferometry,” J. Opt. Soc. Am., JOSA 56, 727–733 (1966).

55. H. Du, “Mie-scattering calculation,” Appl. Opt., AO 43, 1951–1956 (2004).

56. X. Ma, P. Qi, Z. Wu, Y. Yang, C. Li, and J. Zhong, “Image Contrast Enhancement by Using LED Annular Oblique Illumination in Bright-Field Microscopy,” Photonics 10, 404 (2023).

57. J. Huang, Y. Bao, and T. K. Gaylord, “Three-dimensional phase optical transfer function in axially symmetric microscopic quantitative phase imaging,” J. Opt. Soc. Am. A, JOSAA 37, 1857–1872 (2020).

58. X. Chen, J. Li, S. M. H. Rafsanjani, and O. Korotkova, “Synthesis of Im-Bessel correlated beams via coherent modes,” Opt. Lett., OL 43, 3590–3593 (2018).

59. M. E. Kandel, M. Fanous, C. Best-Popescu, and G. Popescu, “Real-time halo correction in phase contrast imaging,” Biomed. Opt. Express, BOE 9, 623–635 (2018).

60. T. H. Nguyen, M. E. Kandel, M. Rubessa, M. B. Wheeler, and G. Popescu, “Gradient light interference microscopy for 3D imaging of unlabeled specimens,” Nat Commun 8, 210 (2017).

61. M. Rinehart, Y. Zhu, and A. Wax, “Quantitative phase spectroscopy,” Biomed. Opt. Express, BOE 3, 958–965 (2012)

