## Supplementary Material for "SPECTRAM: three-dimensional full-vector simulation of coherence-tunable microscopy images"


**JIAQI YAO[1] AND XI CHEN[1,*]**

[1] *Department of Biomedical Engineering, Northwestern University, Evanston, Illinois, 60208, USA*


This supplemental material is organized as follows. Sections 1-3 present the theoretical formulation and numerical implementation of SPECTRAM for brightfield, differential interference contrast, and phase-contrast microscopy, respectively. It includes electromagnetic-field calculations, optical transformations, coherence models, and z-stack generation. Section 4 validates SPECTRAM by comparing its simulated scattered fields with Mie theory in both the spatial-frequency and spatial domains. Finally, Section 5 presents the experimentally measured temporal-frequency spectrum and angular distribution of the illumination and describes the corresponding distributions adopted in SPECTRAM.

#### 1. THEORY OF BRIGHTFIELD MICROSCOPY

##### A. Light as electromagnetic waves

As electromagnetic wave, light can be described by the following complex-valued vector functions of three-dimensional space $\mathbf{r} = (x, y, z)$ and time $t$:

$$\mathbf{E}(\mathbf{r}, t) = [E_x, E_y, E_z](\mathbf{r}, t) : \mathbb{R}^4 \to \mathbb{C}^3, \tag{S1}$$

$$\mathbf{H}(\mathbf{r}, t) = [H_x, H_y, H_z](\mathbf{r}, t) : \mathbb{R}^4 \to \mathbb{C}^3. \tag{S2}$$

Take the electrical fields as an example, the three components $E_x$, $E_y$, and $E_z$ represent the polarization of light in the $x$, $y$, and $z$ directions, respectively. Hence, electromagnetic fields are also vector fields. The complex-valued fields retain both the amplitude and phase information of light.

In the temporal domain, the electrical fields are governed by Maxwell's equations. In our model, we first make the following assumptions:

- Except for the prescribed source distributions used to generate the illumination, the surrounding medium and the sample contain no free charges or conduction currents.
- The surrounding medium and sample are linear, isotropic, dispersion-free and lossless.

Under these assumptions, the Maxwell's equations can be expressed as

$$\nabla \cdot [\epsilon(\mathbf{r})\mathbf{E}(\mathbf{r}, t)] = \rho_s(\mathbf{r}, t), \tag{S3a}$$

$$\nabla \cdot [\mu(\mathbf{r})\mathbf{H}(\mathbf{r}, t)] = 0, \tag{S3b}$$

$$\nabla \times \mathbf{E}(\mathbf{r}, t) = -\mu(\mathbf{r})\frac{\partial \mathbf{H}(\mathbf{r}, t)}{\partial t}, \tag{S3c}$$

$$\nabla \times \mathbf{H}(\mathbf{r}, t) = \epsilon(\mathbf{r})\frac{\partial \mathbf{E}(\mathbf{r}, t)}{\partial t} + \mathbf{J}_s(\mathbf{r}, t). \tag{S3d}$$

Here $\epsilon(\mathbf{r})$ and $\mu(\mathbf{r})$ are the permittivity and permeability of the medium-sample system, respectively. In free space, they are denoted as $\epsilon_0$ and $\mu_0$. The speed of light in free space is $c = 1/\sqrt{\epsilon_0 \mu_0}$. $\rho_s(\mathbf{r}, t)$ and $\mathbf{J}_s(\mathbf{r}, t)$ denote the prescribed free-charge and electric-current densities that represent the incident light source, respectively. They vanish outside the source region.

The prescribed source densities are not independent but satisfy the charge-continuity equation

$$\frac{\partial \rho_s(\mathbf{r}, t)}{\partial t} + \nabla \cdot \mathbf{J}_s(\mathbf{r}, t) = 0. \tag{S4}$$

#### B. Poynting vector and energy flux

Though complex-valued variables provide a convenient way to describe the amplitude and phase of light, they are not directly measurable. The instantaneous physical electromagnetic fields are the real parts of the complex-valued fields:

$$\mathbf{e}(\mathbf{r},t) = \mathrm{Re}[\mathbf{E}(\mathbf{r},t)], \quad \mathbf{h}(\mathbf{r},t) = \mathrm{Re}[\mathbf{H}(\mathbf{r},t)]. \tag{S5}$$

.

The instantaneous energy flux of light is described by the Poynting vector, which is defined as

$$\mathbf{s}(\mathbf{r},t) = \mathbf{e}(\mathbf{r},t) \times \mathbf{h}(\mathbf{r},t). \tag{S6}$$

At the camera plane, the final image $I(\mathbf{r})$ is formed by the time-averaged energy flux, which is given by [1]

$$I(\mathbf{r}) = \frac{1}{T}\int_0^T [\mathbf{s}(\mathbf{r},t)\cdot\hat{\mathbf{n}}]\mathrm{d}t = [\frac{1}{T}\int_0^T \mathbf{s}(\mathbf{r},t)\mathrm{d}t]\cdot\hat{\mathbf{n}}, \tag{S7}$$

where $\hat{\mathbf{n}}$ is the unit vector normal to the camera plane. The model places the camera perpendicular to the axial direction, i.e. $\hat{\mathbf{n}} = [0,0,1]$. $T$ is the exposure time of the camera, which takes milliseconds. In our model, we assume that $T$ is much longer than the period of light oscillation, which is on the order of femtoseconds. Hence, we can take $T \to \infty$ in our model.

#### C. Wave decomposition under temporal Fourier transform

Transmitted microscopies are usually illuminated by broadband light sources, such as LEDs and halogen lamps. They have a wide bandwidth and can be modeled as a superposition of monochromatic light sources with different temporal frequencies $\omega \in \mathbb{R}$. Under the long exposure time of camera, we can first define the finite time Fourier transform of the electromagnetic fields as

$$\mathbf{E}_T(\mathbf{r},\omega) = \int_0^T \mathbf{E}(\mathbf{r},t)e^{i\omega t}\mathrm{d}t, \tag{S8a}$$

$$\mathbf{H}_T(\mathbf{r},\omega) = \int_0^T \mathbf{H}(\mathbf{r},t)e^{i\omega t}\mathrm{d}t, \quad \omega \in \mathbb{R}. \tag{S8b}$$

They can be seen as the accumlated electromagnetic fields within exposure time $T$. Under this definition, the broadband superposition can be expressed by the inverse temporal Fourier transform of the electromagnetic fields:

$$\mathbf{E}(\mathbf{r},t) = \frac{1}{2\pi}\int_{-\infty}^{\infty} \mathbf{E}_T(\mathbf{r},\omega)e^{-i\omega t}\mathrm{d}\omega, \tag{S9a}$$

$$\mathbf{H}(\mathbf{r},t) = \frac{1}{2\pi}\int_{-\infty}^{\infty} \mathbf{H}_T(\mathbf{r},\omega)e^{-i\omega t}\mathrm{d}\omega. \tag{S9b}$$

Take the electrical fields as an example, the transform consists of the following two terms:

- $\mathbf{E}_T(\mathbf{r},\omega) : \mathbb{R}^4 \to \mathbb{C}^3$ is the finite time complex magnitude of the electrical fields at frequency $\omega$. It has both amplitude and phase information. Its overall amplitude represents the spectral power density of the light source at frequency $\omega$, while the local amplitude and phase represent the spatial distribution.
- $e^{-i\omega t}$ is the temporal oscillation of the electrical fields.

We then analyze the effect of the complex amplitude at different wavelengths on the final image. The real-valued Poynting vector can be expressed as

$$\begin{aligned}\mathbf{s}(\mathbf{r},t) &= \mathbf{e}(\mathbf{r},t) \times \mathbf{h}(\mathbf{r},t)\\ &= \mathrm{Re}[\mathbf{E}(\mathbf{r},t)] \times \mathrm{Re}[\mathbf{H}(\mathbf{r},t)].\end{aligned} \tag{S10}$$

**Identity** For any complex-valued vectors $\mathbf{a},\ \mathbf{b} \in \mathbb{C}^3$, we have the following identity:

$$\mathrm{Re}[\mathbf{a}] \times \mathrm{Re}[\mathbf{b}] \equiv \frac{1}{2}\mathrm{Re}[\mathbf{a}\times\mathbf{b}^*] + \frac{1}{2}\mathrm{Re}[\mathbf{a}\times\mathbf{b}]. \tag{S11}$$

Apply this identity to Eq. S10, we have

$$\mathbf{s}(\mathbf{r},t) = \underbrace{\frac{1}{2}\mathrm{Re}[\mathbf{E}(\mathbf{r},t)\times\mathbf{H}^*(\mathbf{r},t)]}_{\mathbf{T_1}(\mathbf{r},t)} + \underbrace{\frac{1}{2}\mathrm{Re}[\mathbf{E}(\mathbf{r},t)\times\mathbf{H}(\mathbf{r},t)]}_{\mathbf{T_2}(\mathbf{r},t)}, \tag{S12}$$

and the image can be expressed as

$$I(\mathbf{r}) = [\frac{1}{T}\int_0^T \mathbf{T_1}(\mathbf{r},t)\mathrm{d}t]\cdot\hat{\mathbf{n}} + [\frac{1}{T}\int_0^T \mathbf{T_2}(\mathbf{r},t)\mathrm{d}t]\cdot\hat{\mathbf{n}}. \tag{S13}$$

Assuming the function satisfies the condition of absolute integrability, with the Cross-Parseval and Fubini Theorem, the first term can be simplified as

$$\begin{aligned}\frac{1}{T}\int_0^T \mathbf{T_1}(\mathbf{r},t)\mathrm{d}t &= \frac{1}{T}\int_0^T \frac{1}{2}\mathrm{Re}\{\mathbf{E}(\mathbf{r},t)\times\mathbf{H}^*(\mathbf{r},t)\}\mathrm{d}t\\ &= \int_0^T \mathrm{Re}\{\frac{1}{4\pi T}[\int_{-\infty}^{\infty}\mathbf{E}_T(\mathbf{r},\omega)e^{-i\omega t}\mathrm{d}\omega]\times\mathbf{H}^*(\mathbf{r},t)\}\mathrm{d}t\\ &= \int_{-\infty}^{\infty}\mathrm{Re}\{\frac{1}{4\pi T}\mathbf{E}_T(\mathbf{r},\omega)\times[\int_0^T \mathbf{H}(\mathbf{r},t)e^{i\omega t}\mathrm{d}t]^*\}\mathrm{d}\omega\\ &= \int_{-\infty}^{\infty}\mathrm{Re}\{\frac{1}{4\pi T}\mathbf{E}_T(\mathbf{r},\omega)\times\mathbf{H}_T^*(\mathbf{r},\omega)\}\mathrm{d}\omega. \end{aligned} \tag{S14}$$

We then define the following terms:

$$P_0 = \int_{-\infty}^{\infty}\frac{1}{T}\mathcal{M}_s\left[\mathbf{E}_T(\mathbf{r},\omega)\times\mathbf{H}_T^*(\mathbf{r},\omega)\right]\mathrm{d}\omega, \tag{S15a}$$

$$p_\omega(\omega) = \frac{1}{P_0 T}\mathcal{M}_s\left[\mathbf{E}_T(\mathbf{r},\omega)\times\mathbf{H}_T^*(\mathbf{r},\omega)\right], \tag{S15b}$$

$$\hat{\mathbf{E}}(\mathbf{r},\omega) = \frac{e^{-i\phi(\omega)}}{\sqrt{2\pi P_0 p_\omega(\omega)T}}\mathbf{E}_T(\mathbf{r},\omega), \quad \hat{\mathbf{H}}(\mathbf{r},\omega) = \frac{e^{-i\phi(\omega)}}{\sqrt{2\pi P_0 p_\omega(\omega)T}}\mathbf{H}_T(\mathbf{r},\omega), \tag{S15c}$$

$$\mathbf{S}(\mathbf{r},\omega) = \frac{1}{2}\mathrm{Re}\{\hat{\mathbf{E}}(\mathbf{r},\omega)\times\hat{\mathbf{H}}^*(\mathbf{r},\omega)\}. \tag{S15d}$$

Their physical meanings are

- $\mathcal{M}_s$ denotes the source-plane power-flux functional, which integrates the normal component of the complex Poynting flux over the illumination reference plane. $P_0$ is the total time-averaged incident power.
- $p_\omega(\omega)$ is the corresponding normalized power spectral density, satisfying $\int_{-\infty}^{\infty} p_\omega(\omega)\mathrm{d}\omega = 1$. Thus, $P_0 p_\omega(\omega)\,\mathrm{d}\omega$ represents the incident power contained within the infinitesimal frequency interval $[\omega, \omega + \mathrm{d}\omega]$.
- The fields $\widehat{\mathbf{E}}(\mathbf{r},\omega)$ and $\widehat{\mathbf{H}}(\mathbf{r},\omega)$ are the normalized complex amplitude at frequency $\omega$. $e^{-i\phi(\omega)}$ means the random phase that accounts for the absence of fixed phase relationships among different frequencies has been removed. The denominator normalises the amplitude in the complex plane to a unit input power.
- $\mathbf{S}(\mathbf{r},\omega)$ defines the complex-valued Poynting vector under complex fields at frequency $\omega$.

Hence, Eq. S14 can be written as

$$\frac{1}{T}\int_0^T \mathbf{T_1}(\mathbf{r},t)\mathrm{d}t = P_0\int_{-\infty}^{\infty} p_\omega(\omega)\mathbf{S}(\mathbf{r},t)\mathrm{d}\omega. \tag{S16}$$

With the same principle, it can be shown that contains only sum-frequency oscillations under light vibration timescale, which cannot be captured by the camera and the temporal average vanishes:

$$\frac{1}{T}\int_0^T \mathbf{T_2}(\mathbf{r},t)\mathrm{d}t = \mathbf{0}. \tag{S17}$$

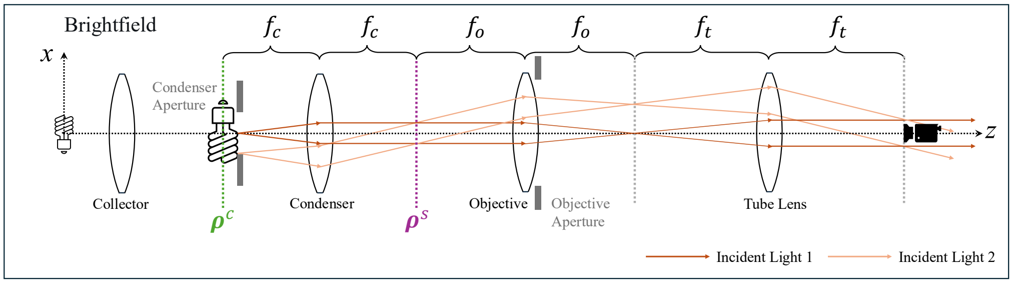


**Fig. S1. Optical configuration and image formation in brightfield microscopy.** The collector and condenser establish Köhler illumination, mapping different source positions to plane waves that illuminate the specimen from different directions. The objective collects the emerging electromagnetic field, and the tube lens forms the image at the camera plane. The relevant focal lengths and transverse coordinate planes are indicated.

In total, at the camera plane, the shown image can be expressed as

$$I(\mathbf{r}) = \left[P_0 \int_{-\infty}^{\infty} p_\omega(\omega)\,(\mathbf{S}(\mathbf{r},\omega)\cdot\hat{\mathbf{n}})\,\mathrm{d}\omega\right]. \tag{S18}$$

That is, the overall image is a linear summation of Poynting vector contribution under all frequency of given illumination light source. This is a result arising from the temporal incoherence property. In SPECTRAM, the cause of the temporal incoherence property is attributed to the interference of light of different frequencies being averaged out during prolonged exposure imaging. In fact, temporal incoherence may also arise from uncertainties in the initial relative phase of light of different frequencies emitted by the light source itself, as well as from the finite length of the wave packet. However, in this context, we are primarily concerned with the effects of prolonged exposure.

#### D. Monochromatic incident wave decomposition under spatial Fourier transform

The previous section concludes that, for a microscopy with broadband light source, the image shown on the camera is given by the linear summation of Poyning vector at each temporal frequency $\omega$ weighted by its power spectral density. Hence, in this section, we focus on the electromagnetic field in monochromatic waves, i.e. single temporal frequency. Eq. S9 and Eq. S15c states that its temporal field can be written as

$$\mathbf{E}(\mathbf{r},t) = \hat{\mathbf{E}}(\mathbf{r},\omega)e^{-i\omega t}, \mathbf{H}(\mathbf{r},t) = \hat{\mathbf{H}}(\mathbf{r},\omega)e^{-i\omega t}. \tag{S19}$$

So the goal is to find the normalized complex magnitude $\hat{\mathbf{E}}, \hat{\mathbf{H}}$ and its corresponding Poynting vector.

**Definition** Before formally introducing the model, we define the spatial Fourier transform here to facilitate the discussion that follows. Consider a monochromatic scalar field $U(\mathbf{r}|\omega)$, defined over the three-dimensional coordinate $\mathbf{r} = (x,y,z)$, propagating in a homogeneous medium with refractive index $n$. its three-dimenional spatial Fourier transform is defined as

$$\widetilde{U}(\mathbf{k}|\omega) = \iiint_{-\infty}^{\infty} U(\mathbf{r}|\omega)e^{-i\mathbf{k}\cdot\mathbf{r}}\mathrm{d}\mathbf{r}, \tag{S20}$$

and the corresponding inverse three-dimenional spatial Fourier transform

$$U(\mathbf{r}|\omega) = \frac{1}{(2\pi)^3}\iiint_{-\infty}^{\infty} \widetilde{U}(\mathbf{k}|\omega)e^{i\mathbf{k}\cdot\mathbf{r}}\mathrm{d}\mathbf{k}. \tag{S21}$$

$\mathbf{k}$ is the three-dimensional spatial frequency vector, or wavevector that satisfies

$$\mathbf{k} = [k_x, k_y, k_z], |\mathbf{k}| = k = \frac{n\omega}{c} = \frac{2\pi n}{\lambda_0}, \tag{S22}$$

where $k$ is the wavenumber.

Without considering evanescent waves, the direction of the wavevector is commonly parameterized using spherical coordinates to explicitly describe the propagation angle. Taking the $z$-axis as the optical axis, we define the polar angle $\theta \in [0, \pi)$ as the angle between the propagation direction and the positive $z$-axis, and the azimuthal angle $\psi \in [0, 2\pi)$ as the angle between the projection of the propagation direction onto the $x-y$-plane and the positive $x$-axis. The wavevector can then be expressed as

$$\mathbf{k} = [k_x, k_y, k_z] = k \begin{bmatrix} \sin\theta\cos\psi \\ \sin\theta\sin\psi \\ \cos\theta \end{bmatrix}^T. \tag{S23}$$

On a 2-dimenional $x-y$ plane $\boldsymbol{\rho} = (x, y)$ at position $z = z^0$, one can also define the two-dimensional spatial Fourier transform

$$\widetilde{U}(\mathbf{k}_\perp|z^0, \omega) = \iint_{-\infty}^{\infty} U(\boldsymbol{\rho}|z^0, \omega) e^{-i\mathbf{k}_\perp \cdot \boldsymbol{\rho}} \mathrm{d}\boldsymbol{\rho}, \tag{S24}$$

where $\mathbf{k}_\perp = [k_x, k_y]$ is the lateral wavevector. The corresponding inverse transform is

$$U(\boldsymbol{\rho}|z^0, \omega) = \frac{1}{(2\pi)^2} \iint_{-\infty}^{\infty} \widetilde{U}(\mathbf{k}_\perp|z^0, \omega) e^{i\mathbf{k}_\perp \cdot \boldsymbol{\rho}} \mathrm{d}\mathbf{k}_\perp. \tag{S25}$$

Figure S1 shows the model of brightfield microscopy in the $x-z$ cross-section. For incident illumination light, as discussed in the main text, every point one the condenser plane $\boldsymbol{\rho}^c$ acts as a light source and generate a plane wave to illuminate the sample. We first consider the case where there is no sample on the stage plane $\boldsymbol{\rho}^s$. Condenser plane $\boldsymbol{\rho}^c = (x^c, y^c)$ and stage plane $\boldsymbol{\rho}^s = (x^s, y^s)$ lies in the two-dimensional $x-y$ coordinates, and their $z$ axis position are $z^c$ and $z^s$, respectively. They are situated at the front and rear focal planes, respectively, of a condenser with a focal length of $f_c$.

We first model the condenser effect under scalar diffraction theory, where the scalar field $U$ can be any components in $\hat{\mathbf{E}}$ and $\hat{\mathbf{H}}$, i.e. $U \in \{E_x, E_y, E_z, H_x, H_y, H_z\}$. Suppose the sclar field on condenser plane is $U^c(\boldsymbol{\rho}^c|z^c, \omega)$, and the incident sclar field on stage plane is $U^s_{\text{inc}}(\boldsymbol{\rho}^s|z^s, \omega)$, the scalar diffraction theory tells [1]

$$U^s_{\text{inc}}(\boldsymbol{\rho}^s|z^s, \omega) = C(\omega) A \iint_{-\infty}^{\infty} U^c(\boldsymbol{\rho}^c|z^c, \omega) e^{-i\frac{k_0}{f_c}\boldsymbol{\rho}^c \cdot \boldsymbol{\rho}^s} \mathrm{d}\boldsymbol{\rho}^c, \quad A = \frac{k_0 e^{2ik_0 f_c}}{i2\pi f_c}, \tag{S26}$$

where $k_0 = \omega/c$ is the wavenumber in the vacuum. Once the incident light has passed through the condenser and entered the source region, we can also define $k = n_{\text{source}}\omega/c$ as the wavenumber in the surrounding medium of illumination source side. $C(\omega) \in \mathbb{R}$ is a normalization factor for fields with different temporal frequency $\omega$. $A$ is a constant. $U^c(\boldsymbol{\rho}^c, z^c)$ describes the magnitude and phase distribution of light source on the aperture plane. For a LED source, we consider the phase at each position to be a random variable taking values between 0 and $2\pi$. We then define

$$\mathbf{k}^c_\perp = -\frac{k_0}{f_c}\boldsymbol{\rho}^c, \quad U^{c'}(\mathbf{k}^c_\perp|z^c, \omega) = U^c(-\frac{\mathbf{k}^c_\perp f_c}{k_0}|z^c, \omega) = U^c(\boldsymbol{\rho}^c|z^c, \omega). \tag{S27}$$

Hence,

$$U^s_{\text{inc}}(\boldsymbol{\rho}^s|z^s, \omega) = C(\omega)(\frac{f_c}{k_0})^2 A \iint_{-\infty}^{\infty} U^{c'}(\mathbf{k}^c_\perp|z^c, \omega) e^{i\mathbf{k}^c_\perp \cdot \boldsymbol{\rho}^s} \mathrm{d}\mathbf{k}^c_\perp. \tag{S28}$$

This shows the 2-dimensional Fourier transform between the condenser aperture and stage plane. More specifically, the incident field consists of multiple plane waves with different $x-y$ spatial frequency $\mathbf{k}^c_\perp$, each of which is determined by a single point on the condenser aperture.

The condenser aperture plane $\boldsymbol{\rho}^c$ can be modeled as a finite circular area with radius $|\boldsymbol{\rho}^c|_{\text{max}}$, and so is the $x-y$ spatial frequency support $\mathbf{k}^c_\perp$. Using the definition of wavevector in spherical coordinate in Eq. S23 and equation Eq. S27, the maximum polar angle $\theta^{\text{Illu}}_{\text{max}}$ of illumination light propagating direction is found as

$$|\mathbf{k}^c_\perp|_{\text{max}} = \frac{\omega}{c} n_{\text{source}} \sin\theta^{\text{Illu}}_{\text{max}} = \frac{\omega}{cf_c}|\boldsymbol{\rho}^c|_{\text{max}} \Rightarrow \theta^{\text{Illu}}_{\text{max}} = \arcsin(\frac{|\boldsymbol{\rho}^c|_{\text{max}}}{n_{\text{source}} f_c}). \tag{S29}$$

Based on this polar angle, the microscopy illumination numerical aperture $\mathrm{NA}_{\mathrm{Illu}}$ is defined as

$$\mathrm{NA}_{\mathrm{Illu}} = n_{\mathrm{source}} \sin \theta_{\mathrm{max}}^{\mathrm{Illu}}. \tag{S30}$$

It measures the spatial frequency range of the illumination light under single-frequency light illumination. To increase this range, one can either increase the refractive index of the medium $n_{\mathrm{source}}$ or increase the maximum polar angle $\theta_{\mathrm{max}}^{\mathrm{Illu}}$.

The integration in frequency domain in Eq. S28 can also be written in spherical coordinate, which can be regarded as the scalar Debye diffraction integral [2]. It takes into account both the Jacobian resulting from the coordinate transformation and the reciprocal aplanatic apodization factor required by power conservation between the planar pupil and the Gaussian reference sphere. It can be expressed as

$$\begin{aligned} U_{\mathrm{inc}}^{s}(\boldsymbol{\rho}^s|z^s,\omega) &= C(\omega) n_{\mathrm{source}}^2 f_c^2 A \int_0^{\theta_{\mathrm{max}}^{\mathrm{Illu}}} \int_0^{2\pi} U^{c'}(\mathbf{k}_\perp^c|z^c,\omega) \sin\theta\cos\theta \frac{1}{\sqrt{\cos\theta}} \exp[i\mathbf{k}_\perp^c(\theta,\psi)\cdot\boldsymbol{\rho}^s] \mathrm{d}\psi \mathrm{d}\theta \\ &= C(\omega) A' \int_0^{\theta_{\mathrm{max}}^{\mathrm{Illu}}} \int_0^{2\pi} U^{c''}(\theta,\psi|\omega) \exp[i\mathbf{k}_\perp^c(\theta,\psi)\cdot\boldsymbol{\rho}^s] \mathrm{d}\psi \mathrm{d}\theta, \end{aligned} \tag{S31}$$

where $A' = A n_{\mathrm{source}}^2 f_c^2$ is a constant,

$$U^{c''}(\theta,\psi|\omega) = U^{c'}(\mathbf{k}_\perp^c(\theta,\psi)|z^c,\omega)\sin\theta\sqrt{\cos\theta}, \;\; \mathbf{k}_\perp^c(\theta,\psi) = k \begin{bmatrix} \sin\theta\cos\psi \\ \sin\theta\sin\psi \end{bmatrix}^T, \;\; k = \frac{n_{\mathrm{source}}\omega}{c}. \tag{S32}$$

This equation describes the sclar field at one certain $x-y$ plane. To generalize it to the 3-dimenional space, we need to first introduce free space propagation.

**Definition** Still, we consider a monochromatic scalar field $U(\mathbf{r}|\omega)$ in the 3-dimenional space $\mathbf{r} = (x, y, z)$ in the homogeneous medium with refractive index $n$. Also, there are no volumetric sources within the region. Then, Maxwell's equation, which controls the electromagnetic fields, can be deduced to the scalar Helmholtz equation

$$(\nabla^2 + k^2) U(\mathbf{r}|\omega) = 0, \tag{S33}$$

where $k = n\omega/c$.
The free-space propagation is formulated as follows: given a field $U(\boldsymbol{\rho}_1|z_1,\omega)$ in one $x-y$ plane at $z_1$, how can one obtain the field on the whole 3-dimensional domain. Here we don't consider the evanescent waves. The angular specturm method is an accurate method for solving this problem, whose solution satisfies the Helmholtz equation. It works in the spatial frequency domain, and the solution in $\mathbf{r} = (\boldsymbol{\rho}, z) = (x, y, z)$ is

$$U(\mathbf{r}|\omega) = \frac{1}{(2\pi)^2} \iint_{-\infty}^{\infty} \widetilde{U}(\mathbf{k}_\perp|z_1,\omega) e^{ik_z(z-z_1)} e^{i\mathbf{k}_\perp\cdot\boldsymbol{\rho}} \mathrm{d}\mathbf{k}_\perp, \tag{S34}$$

where $\widetilde{U}(\mathbf{k}_\perp|z_1,\omega)$ is the 2-d spatial Fouier transform of $U(\boldsymbol{\rho}_1|z_1,\omega)$ defined as Eq. S24. $k_z = \sqrt{k^2 - (k_x^2 + k_y^2)}$.

Since the incident field $U_{\mathrm{inc}}^s$ is propagating in the uniform medium with $n_{\mathrm{source}}$, we can generalize it to the 3-dimentional

$$U_{\mathrm{inc}}^s(\mathbf{r}^s|\omega) = C(\omega) A' \int_0^{\theta_{\mathrm{max}}^{\mathrm{Illu}}} \int_0^{2\pi} U^{c'''}(\theta,\psi|\omega) \exp[i\mathbf{k}^c(\theta,\psi)\cdot\ \mathbf{r}^s] \mathrm{d}\psi \mathrm{d}\theta, \tag{S35}$$

where $U^{c'''}(\theta,\psi|\omega) = U^{c''}(\theta,\psi|\omega) e^{-ik_z^c z^s}$ and $k_z^c(\theta,\psi) = k\cos\theta$.

This scalar field can then be generalized to vector electromagnetic fields

$$\hat{\mathbf{E}}_{\mathrm{inc}}^s(\mathbf{r}^s|\omega) = C(\omega) A' \int_0^{\theta_{\mathrm{max}}^{\mathrm{Illu}}} \int_0^{2\pi} U^{c'''}(\theta,\psi|\omega) \hat{\mathbf{E}}_{\mathrm{inc}}^s(\mathbf{r}^s|\theta,\psi,\omega) \mathrm{d}\psi \mathrm{d}\theta, \tag{S36a}$$

$$\hat{\mathbf{H}}_{\mathrm{inc}}^s(\mathbf{r}^s|\omega) = C(\omega) \frac{A'}{Z_{\mathrm{source}}} \int_0^{\theta_{\mathrm{max}}^{\mathrm{Illu}}} \int_0^{2\pi} U^{c'''}(\theta,\psi|\omega) \hat{\mathbf{H}}_{\mathrm{inc}}^s(\mathbf{r}^s|\theta,\psi,\omega) \mathrm{d}\psi \mathrm{d}\theta, \tag{S36b}$$

where $Z_{\text{source}} = \sqrt{\mu_{\text{source}}/\epsilon_{\text{source}}}$ is the root squre ratio between permeability and permittivity in the medium, and

$$\hat{\mathbf{E}}^s_{\text{inc}}(\mathbf{r}^s|\theta,\psi,\omega) = \vec{\mathbf{e}}(\theta,\psi)\exp[i\mathbf{k}^c(\theta,\psi)\cdot\ \mathbf{r}^s], \tag{S37a}$$

$$\hat{\mathbf{H}}^s_{\text{inc}}(\mathbf{r}^s|\theta,\psi,\omega) = \vec{\mathbf{h}}(\theta,\psi)\exp[i\mathbf{k}^c(\theta,\psi)\cdot\ \mathbf{r}^s], \tag{S37b}$$

are normalized vector fields. $\vec{\mathbf{e}}(\theta,\psi)$ and $\vec{\mathbf{h}}(\theta,\psi)$ are unit vectors that represent the polarization direction. Since its temporal oscillating version S19 satisfies the Maxwell's equation, it can be yielded that

$$\vec{\mathbf{h}}(\theta,\psi) = \frac{\mathbf{k}^c(\theta,\psi)}{k}\times\vec{\mathbf{e}}(\theta,\psi), \quad \vec{\mathbf{e}}(\theta,\psi)\cdot\vec{\mathbf{h}}(\theta,\psi) = 0, \tag{S38a}$$

$$\vec{\mathbf{e}}(\theta,\psi)\cdot\mathbf{k}^c(\theta,\psi) = 0, \quad \vec{\mathbf{h}}(\theta,\psi)\cdot\mathbf{k}^c(\theta,\psi) = 0. \tag{S38b}$$

As discussed above, the initial phase at each point on the condenser aperture plane can be considered as a random variable, and so is $U^{c'''}(\theta,\psi|\omega)$. Hence, it can be written as

$$U^{c'''}(\theta,\psi|\omega) = |U^{c'''}(\theta,\psi|\omega)|e^{i\varphi(\theta,\psi|\omega)}, \tag{S39}$$

where $\varphi(\theta,\psi|\omega)$ is a random variable that is uniformly distributed in $[0,2\pi)$. The resulting Poynting vector needs to be written as a exptectation under the joint probability density distribution of $\varphi$ at different $[\theta,\psi]$. Based on the properties of this random variable, the calculation of the Poynting vector for the incident light can be divided into two extreme cases.

##### *D.1. Case 1: Absolute coherence of incident plane waves*

Although the phase term at different positions is a random variable, it is also possible that the relative phase remains constant across all positions. In this case, the plane waves generated at different positions on the condenser aperture plane retain absolute coherence. Without loss of generality, it can be assumed that $\varphi(\theta,\psi|\omega) = 0, \forall[\theta,\psi]$. In this absolute coherent case, the total Poynting vector of incident light can be written as

$$\begin{aligned}\mathbf{S}^s_{\text{inc}}(\mathbf{r}|\omega) &= \mathbb{E}_\varphi\left[\frac{1}{2}\text{Re}\{\hat{\mathbf{E}}^s_{\text{inc}}(\mathbf{r}^s|\omega)\times\hat{\mathbf{H}}^{s*}_{\text{inc}}(\mathbf{r}^s|\omega)\}\right]\\ &= \frac{1}{2}\text{Re}\{\left[C(\omega)A'\int_0^{\theta^{\text{Illu}}_{\max}}\int_0^{2\pi}|U^{c'''}(\theta,\psi|\omega)|\hat{\mathbf{E}}^s_{\text{inc}}(\mathbf{r}^s|\theta,\psi,\omega)\text{d}\psi\text{d}\theta\right]\\ &\quad\times\left[C(\omega)\frac{A'}{Z_{\text{source}}}\int_0^{\theta^{\text{Illu}}_{\max}}\int_0^{2\pi}|U^{c'''}(\theta,\psi|\omega)|\hat{\mathbf{H}}^s_{\text{inc}}(\mathbf{r}^s|\theta,\psi,\omega)\text{d}\psi\text{d}\theta\right]^*\}.\end{aligned} \tag{S40}$$

##### *D.2. Case 2: Absolute incoherence of incident plane waves*

In the second case, we consider that the relative phase at different positions on the condenser aperture plane exhibits greater randomness, which is more agreed with the real situation. For every point $[\theta,\psi]$, there exists a infinity small neighbourhood $\Delta\alpha(\theta,\psi) = (\Delta\theta\Delta\psi)(\theta,\psi)\to 0$ such that $\forall\theta'\in[\theta-\Delta\theta/2,\theta+\Delta\theta/2,\forall\psi'\in[\psi-\Delta\psi/2,\psi+\Delta\psi/2]$, $\varphi(\theta',\psi'|\omega) = \varphi(\theta,\psi|\omega)$. That is, the phase in this small neighbourhood keeps the same. However, for two positions outside this small neighbourhood, their phases are two independent random variables uniformly distributed over $[0,2\pi)$, and so is their relative phase difference. Hence, we have the following expectation

$$\mathbb{E}_\varphi\left[e^{i[\varphi(\theta,\psi|\omega)-\varphi(\theta',\psi'|\omega)]}\right] = \begin{cases}1 & \forall\theta'\in[\theta-\frac{\Delta\theta}{2},\theta+\frac{\Delta\theta}{2}],\psi'\in[\psi-\frac{\Delta\psi}{2},\psi+\frac{\Delta\psi}{2}]\\ 0 & \text{else}\end{cases}. \tag{S41}$$

Hence, the Poynting vector under this expectation is

$$\begin{aligned}&\mathbf{S}^s_{\text{inc}}(\mathbf{r}|\omega) = \mathbb{E}_\varphi\left[\frac{1}{2}\text{Re}\{\hat{\mathbf{E}}^s_{\text{inc}}(\mathbf{r}^s|\omega)\times\hat{\mathbf{H}}^{s*}_{\text{inc}}(\mathbf{r}^s|\omega)\}\right]\\ &= \frac{C^2(\omega)|A'|^2}{2Z_{\text{source}}}\int_0^{\theta^{\text{Illu}}_{\max}}\int_0^{2\pi}|U^{c'''}(\theta,\psi|\omega)|^2\Delta\alpha(\theta,\psi)\frac{1}{2}\text{Re}\{\hat{\mathbf{E}}^s_{\text{inc}}(\mathbf{r}^s|\theta,\psi,\omega)\times\hat{\mathbf{H}}^{s*}_{\text{inc}}(\mathbf{r}^s|\theta,\psi,\omega)\}\text{d}\psi\text{d}\theta.\end{aligned} \tag{S42}$$

The neighbourhood size $\Delta\alpha(\theta,\psi)$ varies as a function of $(\theta,\psi)$. However, on the condenser aperture plane $[k^c_x,k^c_y]$, the coherence neighbourhood size can be assumed to be a constant, called coherence area $\Delta\Omega$. According to the mapping between $[k_x^c, k_y^c]$ and $[\theta, \psi]$, this coherence neighbourhood size in the $[\theta, \psi]$ can be found by the Jacobian as $\Delta\alpha(\theta, \psi) = \Delta\Omega/(f_c^2 \sin\theta\cos\theta)$. Hence,

$$\mathbf{S}_{\text{inc}}^s(\mathbf{r}|\omega) = \int_0^{\theta_{\max}^{\text{Illu}}} \int_0^{2\pi} \frac{C^2(\omega)|A'|^2\Delta\Omega}{2f_c^2 Z_{\text{source}}} \frac{|U^{c'''}(\theta,\psi|\omega)|^2}{\sin\theta\cos\theta} \frac{1}{2}\text{Re}\{\hat{\mathbf{E}}_{\text{inc}}^s(\mathbf{r}^s|\theta,\psi,\omega) \times \hat{\mathbf{H}}_{\text{inc}}^{s*}(\mathbf{r}^s|\theta,\psi,\omega)\} \text{d}\psi\text{d}\theta. \tag{S43}$$

Let the normalization factor become

$$C(\omega) = \sqrt{\frac{2f_c^2 Z_{\text{source}}}{|A'|^2\Delta\Omega} \left[\int_0^{\theta_{\max}^{\text{Illu}}} \int_0^{2\pi} \frac{|U^{c'''}(\theta,\psi|\omega)|^2}{\sin\theta\cos\theta} \text{d}\psi\text{d}\theta\right]^{-1}}, \tag{S44}$$

and define

$$p_{\theta,\psi}(\theta,\psi|\omega) = \frac{C^2(\omega)|A'|^2\Delta\Omega}{2f_c^2 Z_{\text{source}}} \frac{|U^{c'''}(\theta,\psi|\omega)|^2}{\sin\theta\cos\theta} = \frac{\frac{|U^{c'''}(\theta,\psi|\omega)|^2}{\sin\theta\cos\theta}}{\int_0^{\theta_{\max}^{\text{Illu}}} \int_0^{2\pi} \frac{|U^{c'''}(\theta,\psi|\omega)|^2}{\sin\theta\cos\theta} \text{d}\psi\text{d}\theta}. \tag{S45}$$

It describes the relative contribution of the plane wave propagating in the direction $[\theta, \psi]$ to the illumination field at frequency $\omega$. Equivalently, in probabilistic terms, if the incident light field is approximated using multiple plane waves of unit magnitude, this represents the probability density function for the propagation angle of each plane wave. It should be noted that this differs from the incident light's power density distribution of $[\theta, \psi]$, which requires further weighting of the power generated by different plane waves.

The choice of $C(\omega)$ is to ensure $\int_0^{\theta_{\max}^{\text{Illu}}} \int_0^{2\pi} p_{\theta,\psi}(\theta,\psi|\omega)\text{d}\psi\text{d}\theta = 1$, $\forall\omega$. Hence, $\mathbf{S}_{\text{inc}}^s(\mathbf{r}|\omega)$ can be regarded as normalized Poynting vector at temporal frequency $\omega$, and its generated power on the camera does not vary between different $\omega$.

#### E. Incident illumination light as superposition of plane waves

If the incident plane are absolutely incoherent to each other, take this in Eq. S18, the overall illumination light intensity is

$$I_{\text{inc}}^s(\mathbf{r}) = P_0 \int_{-\infty}^{\infty} p_\omega(\omega) \int_0^{\theta_{\max}^{\text{Illu}}} \int_0^{2\pi} p_{\theta,\psi}(\theta,\psi|\omega) \left[\frac{1}{2}\text{Re}\{\hat{\mathbf{E}}_{\text{inc}}^s(\mathbf{r}^s|\theta,\psi,\omega) \times \hat{\mathbf{H}}_{\text{inc}}^{s*}(\mathbf{r}^s|\theta,\psi,\omega)\} \cdot \hat{\mathbf{n}}\right] \text{d}\psi\text{d}\theta\text{d}\omega. \tag{S46}$$

#### F. Simulating diffraction with FDTD methods

The above describes the model of incident illumination light at the microscope's specimen stage. When a sample is introduced, it perturbs the incident field through scattering and, where applicable, absorption. The interference between the incident and scattered fields produces a complex diffraction pattern that is difficult to describe using a simple analytical model. Nevertheless, the electromagnetic fields still obey Maxwell's equations, allowing the diffraction process to be simulated by solving these equations with numerical methods.

The finite-difference time-domain (FDTD) method is a representative numerical approach for this purpose. It solves Maxwell's curl equations in the temporal domain, which are Eq. S3c and Eq. S3d:

$$\nabla \times \mathbf{E}(\mathbf{r},t) = -\mu(\mathbf{r})\frac{\partial \mathbf{H}(\mathbf{r},t)}{\partial t}, \tag{S47a}$$

$$\nabla \times \mathbf{H}(\mathbf{r},t) = \epsilon(\mathbf{r})\frac{\partial \mathbf{E}(\mathbf{r},t)}{\partial t} + \mathbf{J}_s(\mathbf{r},t). \tag{S47b}$$

with initial condition $\mathbf{E}(\mathbf{r},0) = \mathbf{H}(\mathbf{r},0) = \mathbf{0}$. Boundary condition is selected according to the physical configuration. Perfectly matched layers (PMLs) implement absorbing boundary condition, in which outgoing waves reaching the boundaries are absorbed with minimal artificial reflection. Periodic or Bloch-periodic boundary condition is used to model periodically repeated structures.

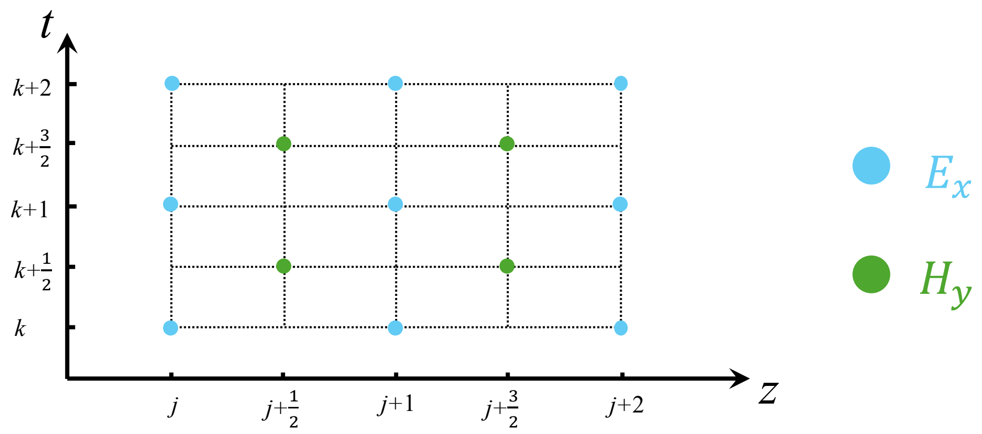


**Fig. S2. Spatial and temporal discretization of electromagnetic fields in the FDTD method.** The electric-field component $E_x$ and magnetic-field component $H_y$ are staggered by half a grid spacing along both the spatial and temporal axes. This arrangement enables alternating field updates through centered finite-difference approximations to Maxwell's curl equations.

In SPECTRAM, unless the permeability $\mu(\mathbf{r})$ and permittivity $\epsilon(\mathbf{r})$ of the simulation target are explicitly specified, all materials are assumed to be nonmagnetic, such that $\mu(\mathbf{r}) = \mu_0$. The permittivity may vary spatially and determines the refractive index distribution by $n(\mathbf{r}) = \sqrt{\epsilon(\mathbf{r})}$.

Since the incident illumination in Eq. S46 has been decomposed into plane-wave components with different temporal frequencies $\omega$ and propagation directions $[\theta, \psi]$, we can first simulate the diffraction for each plane wave separately and then add the results together. Thus, the prescribed current density $\mathbf{J}_s(\mathbf{r}, t|\theta, \psi, \omega)$ in each FDTD simulation should be configured to generate a monochromatic plane wave of unit magnitude. A suitable choice is a continuous-wave (CW) source, which can be expressed by the following current density

$$\mathbf{J}_s(\mathbf{r}, t|\theta, \psi, \omega) = f_s(\mathbf{r}) g(t) \vec{\mathbf{e}}(\theta, \psi) \exp\{i[\mathbf{k}^c(\theta, \psi) \cdot \mathbf{r} - \omega t]\}, \tag{S48}$$

where $f_s(\mathbf{r})$ specifies the spatial region of light source and $g(t)$ is a smooth turn-on function used to suppress transient numerical artifacts. $\vec{\mathbf{e}}(\theta, \psi)$ and $\mathbf{k}^c(\theta, \psi)$ are the polarization direction of electrical field and propagation direction of plane wave under parameter $[\theta, \psi, \omega]$, respectively.

FDTD discretizes both spatial and temporal domain and replacing the derivatives with centered finite-difference approximations, thereby converting the differential equations into explicit algebraic equations. Starting from the initial conditions, the fields are iteratively updated to obtain the solution at each time step. Meanwhile, the resulting solution has been shown to preserve Maxwell's divergence equations Eq. S3a and Eq. S3b:

$$\nabla \cdot [\epsilon(\mathbf{r}) \mathbf{E}(\mathbf{r}, t)] = \rho_s(\mathbf{r}, t), \quad \nabla \cdot [\mu(\mathbf{r}) \mathbf{H}(\mathbf{r}, t)] = 0. \tag{S49}$$

Therefore, it satisfies the complete set of Maxwell's equations, up to numerical discretization errors.

The spatial discretization used in FDTD is known as the Yee grid, in which the electric- and magnetic-field components are spatially staggered and evaluated at interleaved time steps [3]. This arrangement naturally conforms to the curl structure of Maxwell's equations and enables the electric and magnetic fields to be updated alternately using a local leapfrog scheme.

The following simplified example illustrates the basic principle of FDTD. Consider an electromagnetic wave that varies only along the 1-dimenional spatial coordinate $z$ and time $t$. Assuming that the electric and magnetic fields are polarized along the $x$ and $y$-directions, respectively, they can be expressed as $E_x(z, t)$ and $E_y(z, t)$. Then, the first curl equation Eq. S47a can be simplified as

$$\frac{\partial H_y(z, t)}{\partial t} = \frac{1}{\mu(z)} \frac{\partial E_x(z, t)}{\partial z}. \tag{S50}$$

On the Yee grid shown in Fig S2, define

$$E_x^{j,k} = E_x\left(j\Delta z, k\Delta t\right), \quad H_y^{j+\frac{1}{2}, k+\frac{1}{2}} = H_y\left((j+\frac{1}{2})\Delta z, (k+\frac{1}{2})\Delta t\right), \quad \mu^{j+\frac{1}{2}} = \mu\left((j+\frac{1}{2})\Delta z\right), \tag{S51}$$

where $\Delta z$ and $\Delta t$ are the step size of grid on the $z-$ and $t-$ axis, respectively. The derivatives are estimated with centered finite-difference, under which Eq. S50 becomes

$$\frac{H_y^{j+\frac{1}{2},k+\frac{1}{2}} - H_y^{j+\frac{1}{2},k-\frac{1}{2}}}{\Delta t} = \frac{1}{\mu^{j+\frac{1}{2}}} \frac{E_x^{j+1,k} - E_x^{j,k}}{\Delta z}. \tag{S52}$$

The remaining task is then to evaluate the resulting algebraic equation iteratively. The same procedure can be applied to the second curl equation and extended to the full-vector Maxwell equations in three spatial dimensions.

For calculating diffraction at a single temporal frequency, the FDFD method is theoretically more computationally efficient. It solves Maxwell's equations in the frequency domain and eliminates the temporal dimension $t$, thereby theoretically reducing the computational cost. However, solving the FDFD equations usually requires an iterative procedure, which effectively introduces an implicit iteration process analogous to time stepping. Moreover, our experiments show that FDFD is often numerically unstable in our simulations, making its solution difficult to converge. Meanwhile, we demonstrate that our simulation does not need FDTD to be evaluated over an extremely long period of time. Therefore, although FDFD is theoretically more suitable for this problem, we ultimately use FDTD to perform the calculations.

#### G. Diffraction simulation configuration

As described above, SPECTRAM independently simulates the diffraction and corresponding generated image under each monochromatic plane-wave illumination, and then obtains the final simulated image by combining the results of all individual simulations using the weights defined in Eq. S46.

Each simulation is performed in three spatial dimensions and the time domain. Figure S3 shows an $x-z$ plane cross-section of the spatial simulation configuration at $y=0$. The computational domain is surrounded by a perfectly matched layer (PML) with a thickness of one vacuum wavelength $\lambda_0 = \frac{2\pi c}{\omega}$, which absorbs waves propagating toward the boundaries while minimizing artificial reflections [4]. The PML thereby implements an open-boundary condition for the finite computational domain.

The computational domain enclosed by the PML is divided into three main regions—the source, sample, and collection regions—whose refractive indices are set to $n_{\text{source}}$, $n_{\text{sample}}$, and $n_{\text{collection}}$, respectively. The source region contains a current sheet spanning the entire $x$-$y$ plane (red line in Fig. S3), which acts as an external light source and generates a plane wave propagating along the $+z$ direction. The resulting field is a continuous wave with a single angular frequency $\omega$ and a single propagation direction defined by the wavevector $\mathbf{k}^c(\theta, \psi)$, described by Eq. S48. The refractive index of the source region, $n_{\text{source}}$, is set to 1.0 by default, consistent with the common microscopy configuration in which illumination is incident from air.

The sample region contains the sample to be imaged, which perturbs the incident field and generates a diffraction pattern. The collection region is where the diffracted field is collected and recorded. The sample region represents the physical model of the specimen to be imaged. Its background refractive index $n_{\text{sample}}$ corresponds to that of the medium in which the specimen is embedded. In the experiments described in the main text, the bead sample was sandwiched between a glass slide and a coverslip and immersed in deionized water, thereby $n_{\text{sample}} = 1.33$. In the subsequent simulations of cells within the brain organoid, the sample was immersed in cell-culture medium in a Petri dish. Therefore, the refractive index of the background medium was set to $n_{\text{sample}} = 1.336$. Within the surrounding medium, the specimen morphology is modeled through its spatial refractive-index distribution. As an intrinsic optical property of the specimen, the refractive index reflects its underlying biochemical organization. The bead is modeled as a homogeneous sphere with a known refractive index of 1.6. For cells within the brain organoid, images were acquired using gradient light interference microscopy (GLIM), and their refractive-index distribution was reconstructed computationally. As we assume all the samples we imaged are nonmagnetic, the refractive index is modeled by the permittivity distribution as $\epsilon(\mathbf{r}) = n^2(\mathbf{r})$, with $\mu(\mathbf{r}) = \mu_0$.

The source and sample regions are embedded within the collection region, which occupies most of the downstream $+z$ side of the computational domain. The collection region represents the medium through which the microscope objective collects light emerging from the specimen. Low-NA objectives typically operate with air immersion, for which $n_{\text{collection}} = 1.0$. In contrast, high-NA objectives commonly use a high-refractive-index immersion medium, which shortens

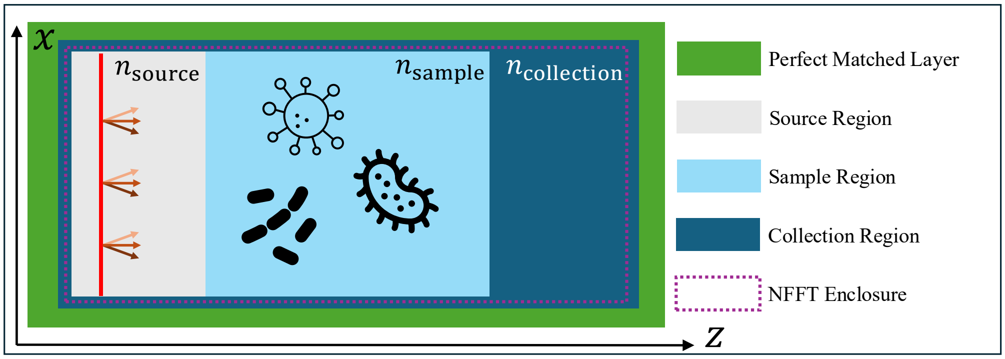


**Fig. S3. Computational domain and field-collection geometry used in SPECTRAM.** The $xz$ cross-section shows the source, sample, and collection regions, whose background refractive indices are specified independently. Perfectly matched layers surround the computational domain, while a closed near-field surface records the electromagnetic fields required for the near-to-far-field transformation.

the wavelength in the medium and increases the achievable NA, thereby extending the range of transverse spatial frequencies collected by the objective. The high NA objective used in our experiments was oil-immersion, and thus $n_{\text{collection}} = 1.515$. The collection region is located primarily on the $+z$ side because, in transmitted-light microscopy, the incident light propagates mainly along the optical $z$-axis and scattering from the specimen is predominantly forward-directed. Therefore, the $+z$ side constitutes the primary light-collection region. Within the collection region, a closed near-field surface is defined for the near-to-far-field transformation, which is shown as the purple line in Fig. S3. The equivalent electric and magnetic surface currents on this enclosure represent the fields generated by the internal CW source and scattered by the specimen, and are used to calculate the angular spectrum of the diffracted light, as detailed in subsection H.

In its time-domain simulation workflow, SPECTRAM employs two frames, each consisting of two stages. The first frame simulates the incident-light field in the absence of a specimen, with the sample region containing only the background medium. The first stage of this frame, referred to as the warm-up stage, allows transient fields to decay and the electromagnetic field within the simulation domain to reach a steady state. Under continuous-wave excitation, the steady-state field oscillates harmonically with a time-independent complex amplitude, as described in Eq. S19:

$$\mathbf{E}_{\text{inc}}(\mathbf{r},t) = \hat{\mathbf{E}}_{\text{inc}}(\mathbf{r}|\omega,\theta,\psi)e^{-i\omega t}, \quad \mathbf{H}_{\text{inc}}(\mathbf{r},t) = \hat{\mathbf{H}}_{\text{inc}}(\mathbf{r}|\omega,\theta,\psi)e^{-i\omega t}. \tag{S53}$$

The second stage lasts for ten optical periods. An optical period is defined as the time required for the electromagnetic field to complete one oscillation and is related to the temporal angular frequency. Ten optical periods is defined as $T(\omega) = 10 \cdot \frac{2\pi}{\omega}$. During this interval, the electromagnetic fields on the near-field enclosure used for the near-to-far-field transformation are accumulated and averaged at the target frequency $\omega$ using the discrete Fourier transform (DFT), thereby obtaining the complex-amplitude distribution of the near field [5]. Given Eq. S53, we found

$$\hat{\mathbf{E}}^{\text{near}}_{\text{inc}}(\mathbf{r}|\omega,\theta,\psi) = \frac{1}{T(\omega)}\int_0^{T(\omega)} \mathbf{E}_{\text{inc}}(\mathbf{r},t)e^{i\omega t}\mathrm{d}t = \hat{\mathbf{E}}_{\text{inc}}(\mathbf{r}|\omega,\theta,\psi), \tag{S54a}$$

$$\hat{\mathbf{H}}^{\text{near}}_{\text{inc}}(\mathbf{r}|\omega,\theta,\psi) = \frac{1}{T(\omega)}\int_0^{T(\omega)} \mathbf{H}_{\text{inc}}(\mathbf{r},t)e^{i\omega t}\mathrm{d}t = \hat{\mathbf{H}}_{\text{inc}}(\mathbf{r}|\omega,\theta,\psi). \tag{S54b}$$

This indicates that the near-field complex amplitude obtained by DFT averaging is equivalent to the steady-state complex amplitude produced by the CW source. Averaging over ten optical periods is primarily used to reduce numerical errors in FDTD and thereby improve the accuracy of the near-field results.

In Frame 2, we use the same simulation configuration but introduce the sample into the computational domain. In this case, the incident field is scattered by the sample, and the scattered field interferes with the incident field. The resulting electromagnetic field, referred to as the total

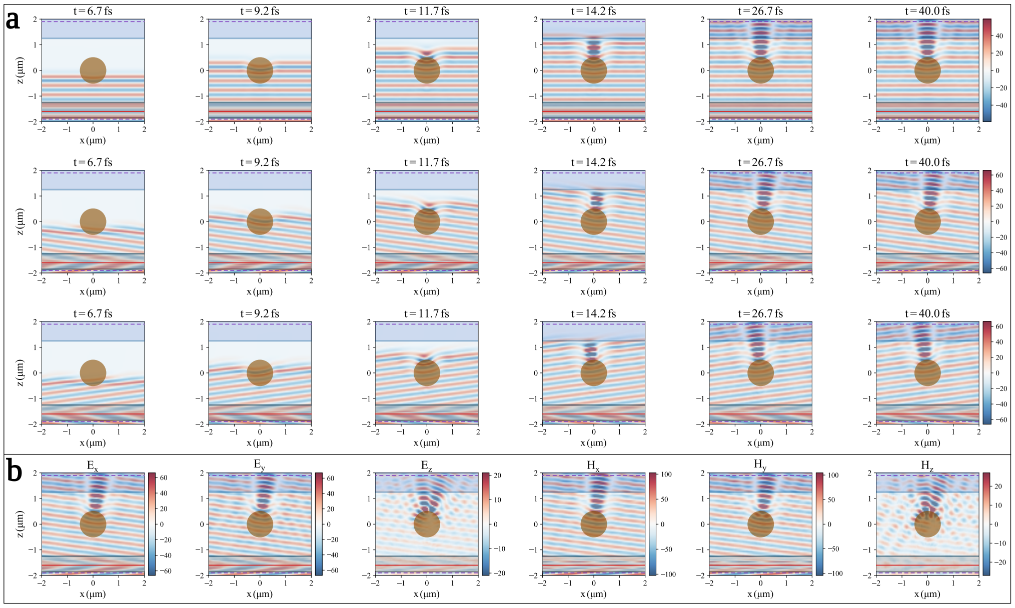


**Fig. S4. Transient evolution and steady-state electromagnetic fields around a polystyrene bead.** (a) Temporal evolution of the real part of $E_x$ for a 1-μm-diameter bead illuminated by plane waves with different propagation directions. The initial warm-up stage establishes the incident and scattered fields, followed by steady periodic oscillations after approximately 26.7 fs. (b) All six electromagnetic-field components in the steady-state regime for $\theta = 10°$ and $\psi = 0°$.

field, can be numerically decomposed into the incident and scattered fields.

$$\mathbf{E}_{\text{tot}}(\mathbf{r}, t) = \hat{\mathbf{E}}_{\text{tot}}(\mathbf{r}|\omega, \theta, \psi)e^{-i\omega t}, \quad \mathbf{H}_{\text{tot}}(\mathbf{r}, t) = \hat{\mathbf{H}}_{\text{tot}}(\mathbf{r}|\omega, \theta, \psi)e^{-i\omega t}, \tag{S55a}$$

$$\hat{\mathbf{E}}_{\text{tot}}(\mathbf{r}|\omega, \theta, \psi) = \hat{\mathbf{E}}_{\text{inc}}(\mathbf{r}|\omega, \theta, \psi) + \hat{\mathbf{E}}_{\text{sca}}(\mathbf{r}|\omega, \theta, \psi), \quad \hat{\mathbf{H}}_{\text{tot}}(\mathbf{r}|\omega, \theta, \psi) = \hat{\mathbf{H}}_{\text{inc}}(\mathbf{r}|\omega, \theta, \psi) + \hat{\mathbf{H}}_{\text{sca}}(\mathbf{r}|\omega, \theta, \psi). \tag{S55b}$$

Similarly, stage 1 of Frame 2 allows the electromagnetic field to reach a steady state. Figure S4a shows the temporal evolution of the real part of the total-field component $E_x$ for a 1-μm-diameter polystyrene bead illuminated by plane waves with different propagation angles $(\theta, \psi)$. During the first 26.7 fs, the incident wave propagates into the computational domain and interacts with the bead, establishing the scattered field. This transient interval corresponds to the warm-up process in stage 1. After 26.7 fs, the field exhibits steady periodic oscillations, corresponding to the steady-state regime in stage 2. Figure S4b shows all vector components of the total electromagnetic field after steady state is reached for illumination at $\theta = 10°$ and $\psi = 0°$.

During stage 2, the fields on the near-field enclosure are accumulated and averaged at the target frequency using the DFT over ten optical periods to obtain their complex amplitudes. The same principle yields that

$$\hat{\mathbf{E}}^{\text{near}}_{\text{tot}}(\mathbf{r}|\omega, \theta, \psi) = \hat{\mathbf{E}}_{\text{tot}}(\mathbf{r}|\omega, \theta, \psi), \quad \hat{\mathbf{H}}^{\text{near}}_{\text{tot}}(\mathbf{r}|\omega, \theta, \psi) = \hat{\mathbf{H}}_{\text{tot}}(\mathbf{r}|\omega, \theta, \psi). \tag{S56}$$

By combining the results from the two frames, we can further obtain the complex amplitude of the scattered field on the near-field enclosure by subtracting the incident field from the total field.

$$\hat{\mathbf{E}}^{\text{near}}_{\text{sca}}(\mathbf{r}|\omega, \theta, \psi) = \hat{\mathbf{E}}^{\text{near}}_{\text{tot}}(\mathbf{r}|\omega, \theta, \psi) - \hat{\mathbf{E}}^{\text{near}}_{\text{inc}}(\mathbf{r}|\omega, \theta, \psi), \tag{S57a}$$

$$\hat{\mathbf{H}}^{\text{near}}_{\text{sca}}(\mathbf{r}|\omega, \theta, \psi) = \hat{\mathbf{H}}^{\text{near}}_{\text{tot}}(\mathbf{r}|\omega, \theta, \psi) - \hat{\mathbf{H}}^{\text{near}}_{\text{inc}}(\mathbf{r}|\omega, \theta, \psi). \tag{S57b}$$

#### H. Angular spectrum collection via Near to Far-field transform

On the enclosure $\mathbf{r}^{\perp} \in S$, we define the equivalent electric and magnetic surface currents as

$$\mathbf{J}^{e}(\mathbf{r}^{\perp}|\omega, \theta, \psi) = \hat{\mathbf{n}}(\mathbf{r}^{\perp}) \times \hat{\mathbf{H}}^{\text{near}}(\mathbf{r}^{\perp}|\omega, \theta, \psi), \tag{S58a}$$

$$\mathbf{M}^{e}(\mathbf{r}^{\perp}|\omega, \theta, \psi) = -\hat{\mathbf{n}}(\mathbf{r}^{\perp}) \times \hat{\mathbf{E}}^{\text{near}}(\mathbf{r}^{\perp}|\omega, \theta, \psi), \tag{S58b}$$

where $\hat{\mathbf{n}}(\mathbf{r}^\perp)$ is the unit normal vector of the surface at $\mathbf{r}^\perp$. This formula is applied separately to the incident and scattered fields to calculate their respective equivalent currents.

Throughout this subsection, all calculations are performed for a fixed $(\omega, \theta, \psi)$. For notational simplicity, we therefore omit the conditioning arguments $|\omega, \theta, \psi$ from the functions, with these quantities understood to remain fixed.

The outward-radiating electromagnetic fields generated by the equivalent electric $\mathbf{J}^e$ and magnetic surface currents $\mathbf{M}^e$ are identical to those produced by the sources and sample scattering enclosed by the surface. These equivalent currents can therefore be used to calculate the complex amplitudes of the fields radiated from the enclosed region into the exterior domain. Because the present calculation is restricted to a single temporal frequency, the frequency-domain Stratton-Chu formulation is well suited for this purpose [6]. Under this formulation, the electromagnetic fields in the region outside the near-field enclosure, referred to as the ***far-field region***, can be calculated as follows:

$$\hat{\mathbf{E}}^{\text{far}}(\mathbf{r}) = \oiint_S \left[ i\omega\mu_c \overline{\overline{\mathbf{G}}}(\mathbf{r}, \mathbf{r}^\perp)\mathbf{J}^e(\mathbf{r}^\perp) - \nabla \times [\overline{\overline{\mathbf{G}}}(\mathbf{r}, \mathbf{r}^\perp)\mathbf{M}^e(\mathbf{r}^\perp)] \right] \mathrm{d}\mathbf{r}^\perp, \tag{S59a}$$

$$\hat{\mathbf{H}}^{\text{far}}(\mathbf{r}) = \oiint_S \left[ i\omega\epsilon_c \overline{\overline{\mathbf{G}}}(\mathbf{r}, \mathbf{r}^\perp)\mathbf{M}^e(\mathbf{r}^\perp) + \nabla \times [\overline{\overline{\mathbf{G}}}(\mathbf{r}, \mathbf{r}^\perp)\mathbf{J}^e(\mathbf{r}^\perp)] \right] \mathrm{d}\mathbf{r}^\perp, \tag{S59b}$$

where $\mu_c$ and $\epsilon_c$ are the relative permeability and permittivity of the collection region medium. The refractive index of the collection region is $n_{\text{collection}} = n_c = \sqrt{\mu_c}$. Under temporal frequency $\omega$, the wavenumber in the collection region is $k_c = \frac{n_c\omega}{c}$. $S$ represents the near-field enclosure surface, $\mathbf{r}^\perp \in S$ is a point on the surface, and $\mathbf{r}$ is a point in the far-field region.

The dyadic Green's function $\overline{\overline{\mathbf{G}}}(\mathbf{r}, \mathbf{r}^\perp)$ describes the propagation of electromagnetic waves from the source point $\mathbf{r}^\perp$ to the observation point $\mathbf{r}$ in the collection region. For three-dimensional vector fields, it also accounts for the coupling between different polarization components and can therefore be represented as a $\mathbb{C}^{3\times 3}$ matrix acting on three-dimensional vector fields $\mathbf{J}^e$ and $\mathbf{M}^e$. Specifically, it can be expressed as

$$\overline{\overline{\mathbf{G}}}(\mathbf{r}, \mathbf{r}^\perp) = \left[ \overline{\overline{\mathbf{I}}} + \frac{1}{k_c^2}\nabla\nabla \right] G(\mathbf{r}, \mathbf{r}^\perp), \tag{S60a}$$

$$G(\mathbf{r}, \mathbf{r}^\perp) = \frac{e^{ik_c|\mathbf{r}-\mathbf{r}^\perp|}}{4\pi|\mathbf{r}-\mathbf{r}^\perp|}. \tag{S60b}$$

where $\overline{\overline{\mathbf{I}}}$ is the $3 \times 3$ identity matrix. $k_c$ is the wavenumber and $G(\mathbf{r}, \mathbf{r}^\perp)$ is the scalar Green's function. $\nabla\nabla$ represents the Hessian matrix of second derivatives.

As shown in Fig. S5a, the far-field diffraction of interest lies on the positive side of the optical axis, namely, in the region beyond the $+z$-facing plane of the near-field enclosure. This region is referred to as the $+z$ ***far region***. In addition, we define an infinite $xy$-plane $\boldsymbol{\rho}^t = (x^t, y^t)$ at the coordinate $z^t$ of the $+z$-facing surface of the near-field enclosure, referred to as the $+\mathbf{z}$ ***top plane***. Unlike the finite computational domain used in the FDTD simulation, this plane extends infinitely in the $x$- and $y$-directions.

According to the surface equivalence theorem and the Silver-Müller radiation condition, the diffraction field in the $+z$ far-field region calculated from the near-field enclosure, in Eq. S59a and S59b, can be shown to be equivalent to that obtained by treating the $+z$ top plane as an equivalent source plane.

$$\hat{\mathbf{E}}^{\text{far}}(\mathbf{r}) = \iint_{-\infty}^{\infty} \left[ i\omega\mu_c \overline{\overline{\mathbf{G}}}(\mathbf{r}, \{\rho^t|z^t\})\mathbf{J}^e(\{\rho^t|z^t\}) - \nabla \times [\overline{\overline{\mathbf{G}}}(\mathbf{r}, \{\rho^t|z^t\})\mathbf{M}^e(\{\rho^t|z^t\})] \right] \mathrm{d}\rho^t, \tag{S61a}$$

$$\hat{\mathbf{H}}^{\text{far}}(\mathbf{r}) = \iint_{-\infty}^{\infty} \left[ i\omega\epsilon_c \overline{\overline{\mathbf{G}}}(\mathbf{r}, \{\rho^t|z^t\})\mathbf{M}^e(\{\rho^t|z^t\}) + \nabla \times [\overline{\overline{\mathbf{G}}}(\mathbf{r}, \{\rho^t|z^t\})\mathbf{J}^e(\{\rho^t|z^t\})] \right] \mathrm{d}\rho^t. \tag{S61b}$$

Note that the FDTD simulation is restricted to a finite spatial domain, and therefore the complete $+z$ top plane cannot be directly obtained from the simulation. Consequently, the near-field enclosure must be used as the equivalent source for the calculation. Based on this construction, as shown in Fig. S5b, we define a spherical observation surface in the $+z$ far-field region, centered at the origin of the simulation domain, with its radius approaching infinity $R \to \infty$. Each point on this surface is parameterized in spherical coordinates $\mathbf{q} = (u, v, w)$ by the polar angle between its

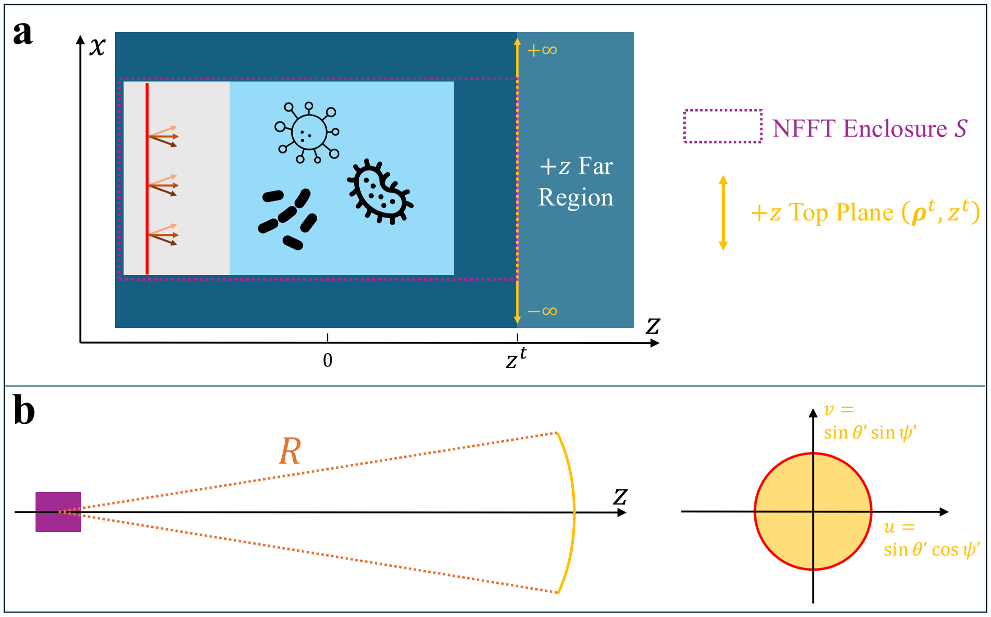


**Fig. S5. Geometrical construction for extracting the angular spectrum through the near-to-far-field transformation.** (a) Definition of the near-field enclosure, the infinite $+z$ top plane, and the forward observation region. (b) Spherical observation geometry used to relate the far-field radiation pattern to the transverse spatial-frequency components of the field on the top plane.

radial direction and the optical axis $\theta'$, together with its azimuthal angle $\psi'$:

$$\mathbf{q} = (u, v, w), \quad u = \sin\theta' \cos\psi', \quad v = \sin\theta' \sin\psi', \quad w = \cos\theta', \tag{S62a}$$

$$\mathbf{r}^{\text{far}} = (x^{\text{far}}, y^{\text{far}}, z^{\text{far}}), \quad x^{\text{far}} = Ru, \quad y^{\text{far}} = Rv, \quad z^{\text{far}} = Rw. \tag{S62b}$$

On this spherical surface, the sclar Green's function can be approximated as

$$|\mathbf{r} - \mathbf{r}^{\perp}| \approx R - \mathbf{q} \cdot \mathbf{r}^{\perp} \Rightarrow G(R\mathbf{q}, \mathbf{r}^{\perp}) \approx \frac{e^{ik_c R}}{4\pi R} e^{-ik_c \mathbf{q} \cdot \mathbf{r}^{\perp}}. \tag{S63}$$

Under this approximation, it can be derived that the far-field diffraction in Eq. S61a and S61b is equivalent to

$$\hat{\mathbf{E}}^{\text{far}}(R\mathbf{q}) = -\frac{ik_c e^{ik_c R}}{2\pi R} w \left[ e^{-ik_z z^t} \widetilde{\mathbf{E}}(\mathbf{k}_{\perp} | z^t) \right], \tag{S64a}$$

$$\hat{\mathbf{H}}^{\text{far}}(R\mathbf{q}) = -\frac{ik_c e^{ik_c R}}{2\pi R} w \left[ e^{-ik_z z^t} \widetilde{\mathbf{H}}(\mathbf{k}_{\perp} | z^t) \right], \tag{S64b}$$

where $k_z = \sqrt{k_c^2 - |\mathbf{k}_{\perp}|^2}$ and

$$\widetilde{\mathbf{E}}(\mathbf{k}_{\perp} | z^t) = \iint_{-\infty}^{\infty} \hat{\mathbf{E}}(\boldsymbol{\rho}^t | z^t) e^{-i\mathbf{k}_{\perp} \cdot \boldsymbol{\rho}^t} \mathrm{d}\boldsymbol{\rho}^t, \tag{S65a}$$

$$\widetilde{\mathbf{H}}(\mathbf{k}_{\perp} | z^t) = \iint_{-\infty}^{\infty} \hat{\mathbf{H}}(\boldsymbol{\rho}^t | z^t) e^{-i\mathbf{k}_{\perp} \cdot \boldsymbol{\rho}^t} \mathrm{d}\boldsymbol{\rho}^t, \tag{S65b}$$

are two dimensional spatial Fourier transforms of the complex amplitudes of the electric and magnetic fields on the $+z$ top plane, respectively. The relationship between the transverse wavevector $\mathbf{k}_{\perp} = (k_x, k_y)$ and spherical coordinates $(u, v)$ is given by $\mathbf{k}_{\perp} = k_c(u, v)$.

Hence, by combining Eq. S59a, Eq. S59b, Eq. S65a and Eq. S65b, the spatial-frequency-domain field associated with the infinite $+z$ top plane can be obtained through the near-to-far-field transformation. Applying this formulation separately to the incident and scattered fields yields their corresponding frequency-domain fields $\widetilde{\mathbf{E}}_{\text{inc}}(\mathbf{k}_{\perp} | z^t)$, $\widetilde{\mathbf{H}}_{\text{inc}}(\mathbf{k}_{\perp} | z^t)$, $\widetilde{\mathbf{E}}_{\text{sca}}(\mathbf{k}_{\perp} | z^t)$, and $\widetilde{\mathbf{H}}_{\text{sca}}(\mathbf{k}_{\perp} | z^t)$.

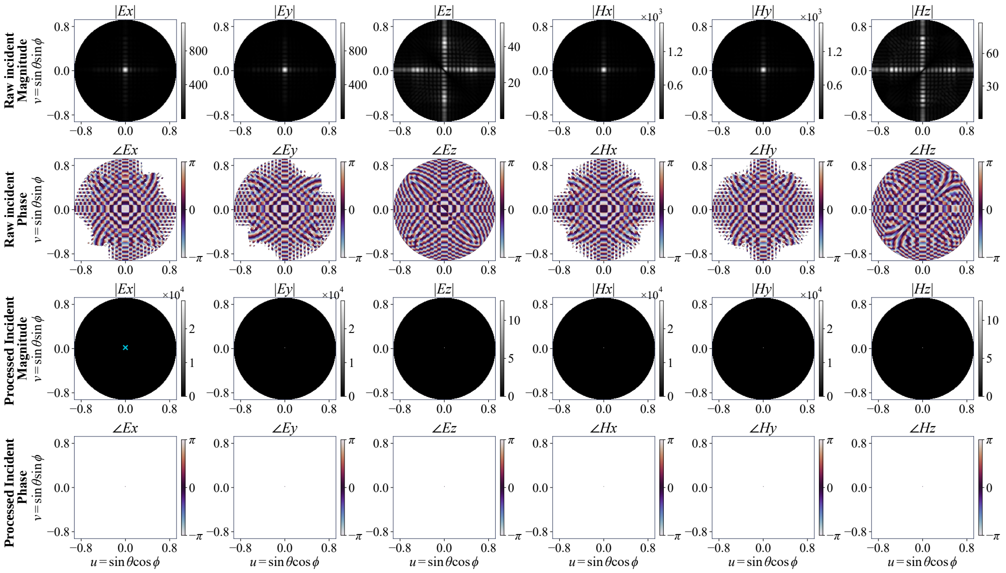

**Fig. S6. Correction of the incident-field spectrum for finite-domain truncation.** The magnitude and phase of all six incident electromagnetic-field components are shown before and after preprocessing. The broadened spectra obtained from the finite computational domain are replaced by discrete plane-wave components at the prescribed spatial-frequency location, preserving the total squared magnitude of each sampled spectrum.

The incident and scattered fields are treated separately because the finite simulation domain introduces truncation effects into the collected incident-field spectrum $\widetilde{\mathbf{E}}_{\text{inc}}(\mathbf{k}_\perp|z^t)$ and $\widetilde{\mathbf{H}}_{\text{inc}}(\mathbf{k}_\perp|z^t)$, requiring preprocessing before subsequent calculations. Figure S6 compares the full-vectoral incident field before and after preprocessing for a normally incident plane wave that is polarized along the $xy$ plane. Ideally, a plane wave is represented by a delta function in the spatial-frequency domain. However, within the finite computational domain bounded by PMLs, the sampled incident field is effectively multiplied by a two-dimensional rectangular window in the $xy$-plane. Consequently, its spatial-frequency spectrum exhibits a product of sinc functions—the Fourier transform of the rectangular window—rather than an ideal delta function. However, the known illumination angles $(\theta, \psi)$ uniquely determine the location of the ideal incident-field delta function in the spatial-frequency domain. We therefore sum the squared magnitudes of the unprocessed incident-field spectrum and use this quantity to determine the amplitude of a single nonzero component at the corresponding spatial-frequency location. This procedure preserves the total squared magnitude of the sampled spectrum while restoring the ideal plane-wave representation, as shown in the bottom two rows of Fig. S6. Figure S7 shows the spatial-frequency-domain representation of the full-vector scattered field. Unlike the incident plane wave, the scattered field originates from the specimen and propagates outward. For the simulation configuration considered here, its collected spectrum can be used directly without the preprocessing applied to the incident field.

### I. Objective-tube lens transformation

Finally, we model the transformation of the sample-side electromagnetic fields to the camera plane by the objective-tube-lens system, together with the objective-scanning mechanism used to generate $z$-stack images.

As shown in Fig. S8, we first assume that the objective focal plane coincides with the $z = 0$ plane of the sample-side simulation domain. Because the objective is oil-immersed, the distance from this focal plane to the objective is $f_o/n_{\text{collection}}$, where $f_o$ denotes the vacuum-referenced physical focal length of the objective. In contrast, the optical paths from the objective to the tube lens and from the tube lens to the camera lie in air; their propagation distances are therefore determined by the corresponding physical focal lengths in vacuum.

The objective-tube-lens transformation can be divided into two stages. First, the electromagnetic field at the sample plane is propagated through the objective to its back focal plane $(\boldsymbol{\rho}^b|z^b) =$

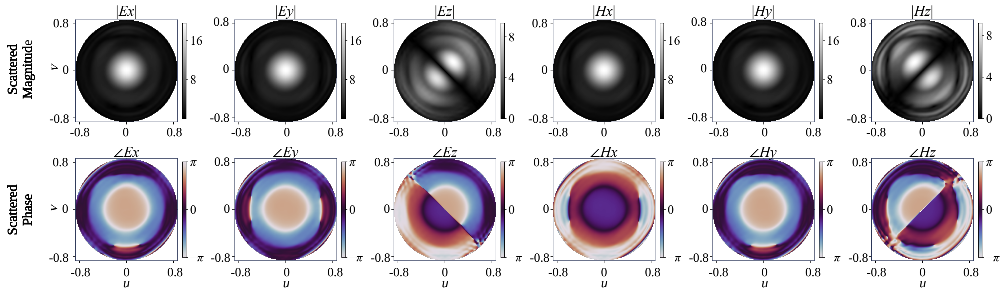


**Fig. S7. Spatial-frequency representation of the full-vector scattered field.** The magnitude and phase of the six scattered electromagnetic-field components are displayed over the transverse spatial-frequency plane $(k_x, k_y)$. These spectra describe the angular distribution of the scattered field and provide the input to the subsequent objective–tube-lens transformation.

$(x^b, y^b|z^b)$. Using the Richards-Wolf integral, the electric field, whether incident field or scattered field, at the objective back focal plane can be expressed as [7]

$$\hat{\mathbf{E}}(\boldsymbol{\rho}^b|z^b) = C_o P_o'(\boldsymbol{\rho}^b) \left[ \sqrt{\gamma_o(\mathbf{k}_\perp)}\, \overline{\overline{\mathbf{R}}}_o(\mathbf{k}_\perp) e^{-ik_z z^t} \widetilde{\mathbf{E}}(\mathbf{k}_\perp|z^t) \right]_{\substack{\mathbf{k}_\perp=(k_x,k_y) \\ k_x=k_0x^b/f_o \\ k_y=k_0y^b/f_o}} . \tag{S66}$$

Here, $k_0 = \frac{\omega}{c}$ is the wavenumber in vacuum. $C_o = -\frac{ik_c\sqrt{n_{\text{collection}}}}{2\pi f_o}$ is a constant. The pupil function $P_o(\boldsymbol{\rho}^b)$ accounts for the finite extent of the objective back aperture, which is typically circular. For an aperture of radius $|\boldsymbol{\rho}|_{\text{max}}$, it can be expressed as

$$P_o'(\boldsymbol{\rho}^b) = \begin{cases} 1, & |\boldsymbol{\rho}^b|^2 \leq |\boldsymbol{\rho}|^2_{\text{max}} \\ 0, & \text{else} \end{cases} . \tag{S67}$$

We wish to transform it to the spatial-frequency domain. Define

$$P_o(\mathbf{k}_\perp) = \begin{cases} 1, & \left(\frac{f_o}{k_0}\right)^2 |\mathbf{k}_\perp|^2 \leq |\boldsymbol{\rho}|^2_{\text{max}} \\ 0, & \text{else} \end{cases} . \tag{S68}$$

Since $k_x = k_0x^b/f_o$, $k_y = k_0y^b/f_o$, we find

$$\hat{\mathbf{E}}(\boldsymbol{\rho}^b|z^b) = C_o \left[ P_o(\mathbf{k}_\perp) \sqrt{\gamma_o(\mathbf{k}_\perp)}\, \overline{\overline{\mathbf{R}}}_o(\mathbf{k}_\perp) e^{-ik_z z^t} \widetilde{\mathbf{E}}(\mathbf{k}_\perp|z^t) \right]_{\substack{\mathbf{k}_\perp=(k_x,k_y) \\ k_x=k_0x^b/f_o \\ k_y=k_0y^b/f_o}} . \tag{S69}$$

Hence, the limit of spatial-frequency magnitude is $|\mathbf{k}_\perp| \leq \frac{k_0|\boldsymbol{\rho}|_{\text{max}}}{f_o}$. If the spatial frequency is expressed in terms of the polar angle $\theta'$ and azimuthal angle $\psi'$, then the limit of polar angle is $k_0 n_{\text{collection}} \sin\theta' \leq \frac{k_0|\boldsymbol{\rho}|_{\text{max}}}{f_o}$. Now define

$$\theta'_{\text{max}} = \arcsin\left(\frac{|\boldsymbol{\rho}|_{\text{max}}}{n_{\text{collection}} f_o}\right), \quad \text{NA}_{\text{obj}} = n_{\text{collection}} \sin\theta'_{\text{max}}. \tag{S70}$$

These define, respectively, the maximum polar angle of the wave components admitted by the objective and the numerical aperture of the objective $\text{NA}_{\text{obj}}$. Hence, the finite aperture function $P_o(\mathbf{k}_\perp)$ can be expressed as

$$P_o(\mathbf{k}_\perp) = \begin{cases} 1, & \frac{|\mathbf{k}_\perp|}{k_0} \leq \text{NA}_{\text{obj}} \\ 0, & \text{else} \end{cases} . \tag{S71}$$

$\gamma_o(\mathbf{k}_\perp) = \frac{\sqrt{k_c^2 - |\mathbf{k}_\perp|^2}}{k_c}$ denotes the apodization factor of the high-NA objective, accounting for the spatial-frequency-dependent variation in field amplitude, as light passes through the objective.

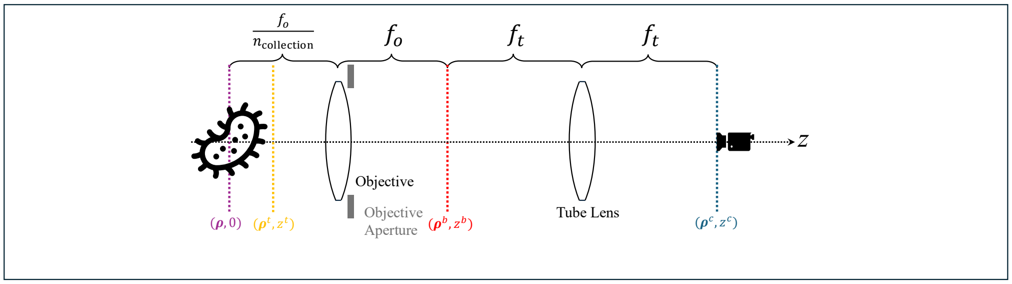


**Fig. S8. Optical geometry of the objective–tube-lens transformation.** The sample-side collection plane, objective back focal plane, and camera plane are identified together with the corresponding propagation distances. This geometry defines the mapping of the collected electromagnetic fields to the camera, including aperture restriction, apodization, polarization transformation, and magnification.

$\overline{\overline{\mathbf{R}}}_o \in \mathbb{R}^{3\times 3}$ is a real orthogonal rotation matrix acting on the vector field, accounting for the change in polarization orientation as light passes through the objective. This vectorial polarization effect becomes increasingly pronounced at high NAs. Here, we do not consider the polarization rotation introduced by the objective in isolation, but rather the net polarization transformation through the combined objective-tube-lens system. We therefore defer the explicit form of $\overline{\overline{\mathbf{R}}}_o$ until both optical elements are considered.

Similarly, we can derive the equation from the back focal plane of the objective to the camera plane. With another Richards-Wolf integral, the electric field, whether incident field or scattered field, at the camera plane in the spatial frequency domain domain $(\mathbf{k}_\perp^c|z^c) = (k_x^c, k_y^c|z^c)$ can be expressed as

$$\widetilde{\mathbf{E}}(\mathbf{k}_\perp^c|z^c) = C_t \left[ P_t'(\boldsymbol{\rho}^b) \frac{1}{\sqrt{\gamma_t'(\boldsymbol{\rho}^b)}} \overline{\overline{\mathbf{R}}}_t(\boldsymbol{\rho}^b) \hat{\mathbf{E}}(\boldsymbol{\rho}^b|z^b) \right]_{\substack{\boldsymbol{\rho}^b=(x^b,y^b) \\ x^b=-f_t k_x^c/k_0 \\ y^b=-f_t k_y^c/k_0}} . \tag{S72}$$

Here $C_t = -\frac{2\pi i f_t}{k_0}$ is a constant. $P_t'(\boldsymbol{\rho}^b)$ accounts for the finite aperture and has the same expression as $P_o'(\boldsymbol{\rho}^b)$:

$$P_t'(\boldsymbol{\rho}^b) = \begin{cases} 1, & |\boldsymbol{\rho}^b|^2 \le |\boldsymbol{\rho}|^2_{\max} \\ 0, & \text{else} \end{cases} . \tag{S73}$$

$\gamma_t'(\boldsymbol{\rho}^b)$ is the apodization factor of the tube lens, and is defined as $\gamma_t'(\boldsymbol{\rho}^b) = \frac{\sqrt{f_t^2 - |\boldsymbol{\rho}^b|^2}}{f_t}$. $\overline{\overline{\mathbf{R}}}_t \in \mathbb{R}^{3\times 3}$ is another real orthogonal rotation matrix that represent the polarization rotation introduced by the tube lens.

Taken together, we build the complete objective-tube-lens transformation model. It applies to the magnetic field as well. The camera plane electromagnetic field in the spatial-frequency domain can be expressed by the field at the sample plane as

$$\widetilde{\mathbf{E}}(\frac{\mathbf{k}_\perp}{M}|z^c) = M \left[ \frac{(n_{\text{collection}} k_0)^2 - |\mathbf{k}_\perp|^2}{k_0^2 - \frac{1}{M^2}|\mathbf{k}_\perp|^2} \right]^{\frac{1}{4}} P(\mathbf{k}_\perp) e^{-ik_z z^t} \left( \overline{\overline{\mathbf{R}}}(\mathbf{k}_\perp) \cdot \widetilde{\mathbf{E}}(\mathbf{k}_\perp|z^t) \right), \tag{S74a}$$

$$\widetilde{\mathbf{H}}(\frac{\mathbf{k}_\perp}{M}|z^c) = M \left[ \frac{(n_{\text{collection}} k_0)^2 - |\mathbf{k}_\perp|^2}{k_0^2 - \frac{1}{M^2}|\mathbf{k}_\perp|^2} \right]^{\frac{1}{4}} P(\mathbf{k}_\perp) e^{-ik_z z^t} \left( \overline{\overline{\mathbf{R}}}(\mathbf{k}_\perp) \cdot \widetilde{\mathbf{H}}(\mathbf{k}_\perp|z^t) \right), \tag{S74b}$$

where $k_z = \sqrt{(n_{\text{collection}} k_0)^2 - |\mathbf{k}_\perp|^2}$. $M = -\frac{f_t}{f_o}$ is the magnification of the objective-tube-lens system. The negative sign indicates that the image is inverted, and the ratio of the tube-lens focal length to the objective focal length determines the magnification. $P(\mathbf{k}_\perp) = P_o(\mathbf{k}_\perp) P_t'(\frac{f_o}{k_0}\mathbf{k}_\perp)$

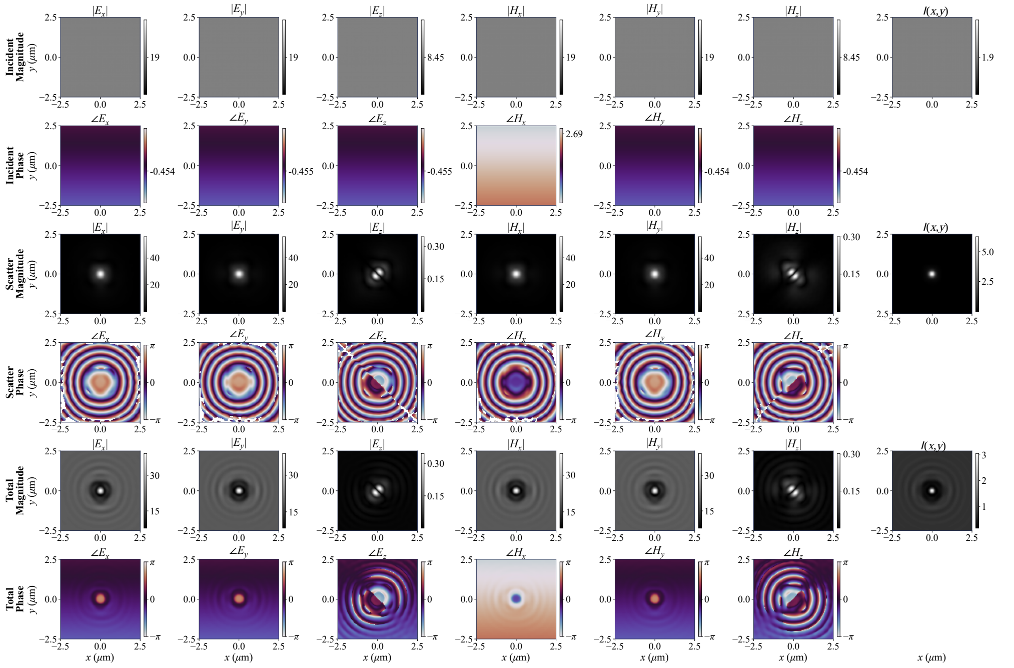


**Fig. S9. Camera-plane field decomposition and image formation under oblique plane-wave illumination.** The magnitude and phase of all six electromagnetic-field components are shown separately for the incident, scattered, and total fields. The rightmost column presents the image intensity calculated from each field using the normal component of the time-averaged Poynting vector.

represents the overlapping support defined by the finite apertures of the objective and tube lens, and is found to be

$$P(\mathbf{k}_\perp) = \begin{cases} 1, & \frac{|\mathbf{k}_\perp|}{k_0} \leq \mathrm{NA}_{\mathrm{obj}} \\ 0, & \text{else} \end{cases} . \tag{S75}$$

The polarization rotation matrix $\overline{\overline{\mathbf{R}}}(\mathbf{k}_\perp) = \overline{\overline{\mathbf{R}}}_o(\mathbf{k}_\perp)\overline{\overline{\mathbf{R}}}_t(\frac{f_o}{k_0}\mathbf{k}_\perp) \in \mathbb{R}^{3\times 3}$ accounts for the combined polarization transformation induced by the objective and tube lens and can be expressed as

$$\overline{\overline{\mathbf{R}}}(\mathbf{k}_\perp) = \vec{\boldsymbol{\theta}}'(\mathbf{k}_\perp)\vec{\boldsymbol{\theta}}^T(\mathbf{k}_\perp) + \vec{\boldsymbol{\psi}}'(\mathbf{k}_\perp)\vec{\boldsymbol{\psi}}^T(\mathbf{k}_\perp), \tag{S76}$$

where

$$\vec{\boldsymbol{\theta}}(\mathbf{k}_\perp) = \begin{bmatrix} \frac{k_x}{|\mathbf{k}_\perp|}\sqrt{1 - \frac{|\mathbf{k}_\perp|^2}{(n_{\mathrm{collection}}k_0)^2}} \\ \frac{k_y}{|\mathbf{k}_\perp|}\sqrt{1 - \frac{|\mathbf{k}_\perp|^2}{(n_{\mathrm{collection}}k_0)^2}} \\ -\frac{|\mathbf{k}_\perp|}{n_{\mathrm{collection}}k_0} \end{bmatrix}, \quad \vec{\boldsymbol{\psi}}(\mathbf{k}_\perp) = \begin{bmatrix} -\frac{k_y}{|\mathbf{k}_\perp|} \\ \frac{k_x}{|\mathbf{k}_\perp|} \\ 0 \end{bmatrix}, \tag{S77a}$$

$$\vec{\boldsymbol{\theta}}'(\mathbf{k}_\perp) = \begin{bmatrix} \frac{k_x}{|\mathbf{k}_\perp|}\sqrt{1 - \frac{|\mathbf{k}_\perp|^2}{(Mk_0)^2}} \\ \frac{k_y}{|\mathbf{k}_\perp|}\sqrt{1 - \frac{|\mathbf{k}_\perp|^2}{(Mk_0)^2}} \\ -\frac{|\mathbf{k}_\perp|}{Mk_0} \end{bmatrix}, \quad \vec{\boldsymbol{\psi}}'(\mathbf{k}_\perp) = \begin{bmatrix} -\frac{k_y}{|\mathbf{k}_\perp|} \\ \frac{k_x}{|\mathbf{k}_\perp|} \\ 0 \end{bmatrix}. \tag{S77b}$$

Hence, when the objective is focused at $z = 0$ sample plane, the microscopy image $I(M\boldsymbol{\rho}, 0)$

can be calculated with

$$\hat{\mathbf{E}}(M\boldsymbol{\rho},0) = \frac{1}{(2\pi M)^2}\iint_{-\infty}^{\infty}\widetilde{\mathbf{E}}(\frac{\mathbf{k}_\perp}{M}|z^c)e^{i\mathbf{k}_\perp\cdot\boldsymbol{\rho}}\mathrm{d}\mathbf{k}_\perp, \tag{S78a}$$

$$\hat{\mathbf{H}}(M\boldsymbol{\rho},0) = \frac{1}{(2\pi M)^2}\iint_{-\infty}^{\infty}\widetilde{\mathbf{E}}(\frac{\mathbf{k}_\perp}{M}|z^c)e^{i\mathbf{k}_\perp\cdot\boldsymbol{\rho}}\mathrm{d}\mathbf{k}_\perp \tag{S78b}$$

$$\mathbf{S}(M\boldsymbol{\rho},0) = \frac{1}{2}\mathrm{Re}\{\hat{\mathbf{E}}(M\boldsymbol{\rho},0)\times\hat{\mathbf{H}}^*(M\boldsymbol{\rho},0)\},\quad I(M\boldsymbol{\rho},0) = \mathbf{S}(M\boldsymbol{\rho},0)\cdot\hat{\mathbf{n}}. \tag{S78c}$$

This applies for both the incident and scattered fields $\hat{\mathbf{E}}_{\text{inc}}$, $\hat{\mathbf{E}}_{\text{scat}}$ and $\hat{\mathbf{H}}_{\text{inc}}$, $\hat{\mathbf{H}}_{\text{scat}}$. Finally, the total field and the corresponding image can be obtained by superposition:

$$\hat{\mathbf{E}}_{\text{tot}}(M\boldsymbol{\rho},0) = \hat{\mathbf{E}}_{\text{inc}}(M\boldsymbol{\rho},0)+\hat{\mathbf{E}}_{\text{sca}}(M\boldsymbol{\rho},0),\quad \hat{\mathbf{H}}_{\text{tot}}(M\boldsymbol{\rho},0) = \hat{\mathbf{H}}_{\text{inc}}(M\boldsymbol{\rho},0)+\hat{\mathbf{H}}_{\text{sca}}(M\boldsymbol{\rho},0), \tag{S79a}$$

$$\mathbf{S}_{\text{tot}}(M\boldsymbol{\rho},0) = \frac{1}{2}\mathrm{Re}\{\hat{\mathbf{E}}_{\text{tot}}(M\boldsymbol{\rho},0)\times\hat{\mathbf{H}}^*_{\text{tot}}(M\boldsymbol{\rho},0)\},\quad I_{\text{tot}}(M\boldsymbol{\rho},0) = \mathbf{S}_{\text{tot}}(M\boldsymbol{\rho},0)\cdot\hat{\mathbf{n}}. \tag{S79b}$$

This represents the complete field and image formation. Hereafter, $\hat{\mathbf{E}} \equiv \hat{\mathbf{E}}_{\text{tot}}$ and $\hat{\mathbf{H}} \equiv \hat{\mathbf{H}}_{\text{tot}}$ denote the total electric and magnetic fields, respectively, and $I \equiv I_{\text{tot}}$ denotes the corresponding image intensity, unless otherwise specified.

Figure S9 shows the magnitude and phase of all vector components of the incident, scattered, and total fields under illumination by an obliquely incident plane wave with its transverse propagation direction along the $y$-axis. The rightmost column shows the corresponding images calculated from each field.

#### J. z-stack imaging simulation

Eq. S78c describes the formation of the two-dimensional camera image when the objective focal plane coincides with the $z = 0$ plane. As shown in Fig. S10, for a given sample, we define $z_{\text{max}}$ as the maximum axial coordinate occupied by the specimen. The region $z > z^{\text{max}}$ is therefore a homogeneous half-space with refractive index $n_{\text{collection}}$. Therefore, the electromagnetic fields on the $z = z_{\text{max}}$ and $z = z^t$ planes can be transformed into one another through free-space propagation. Hence, the Richards-Wolf integral Eq. S66 can be rewritten as

$$\begin{aligned}\hat{\mathbf{E}}(\boldsymbol{\rho}^b|z^b) &= C_o P_o'(\boldsymbol{\rho}^b)\left[\sqrt{\gamma_o(\mathbf{k}_\perp)}\,\overline{\overline{\mathbf{R}}}_o(\mathbf{k}_\perp)e^{-ik_z z^{\text{max}}}\left(\widetilde{\mathbf{E}}(\mathbf{k}_\perp|z^t)e^{ik_z(z^{\text{max}}-z^t)}\right)\right]_{\substack{\mathbf{k}_\perp=(k_x,k_y)\\ k_x=k_0x^b/f_o\\ k_y=k_0y^b/f_o}}\cdot \\ &= C_o P_o'(\boldsymbol{\rho}^b)\left[\sqrt{\gamma_o(\mathbf{k}_\perp)}\,\overline{\overline{\mathbf{R}}}_o(\mathbf{k}_\perp)e^{-ik_z z^{\text{max}}}\widetilde{\mathbf{E}}(\mathbf{k}_\perp|z^{\text{max}})\right]_{\substack{\mathbf{k}_\perp=(k_x,k_y)\\ k_x=k_0x^b/f_o\\ k_y=k_0y^b/f_o}}\cdot\end{aligned} \tag{S80}$$

Subsequently, Eq. S74a and Eq. S74b can be rewritten as

$$\widetilde{\mathbf{E}}(\frac{\mathbf{k}_\perp}{M}|z^c) = M\left[\frac{(n_{\text{collection}}k_0)^2-|\mathbf{k}_\perp|^2}{k_0^2-\frac{1}{M^2}|\mathbf{k}_\perp|^2}\right]^{\frac{1}{4}} P(\mathbf{k}_\perp)e^{-ik_z z^{\text{max}}}\left(\overline{\overline{\mathbf{R}}}(\mathbf{k}_\perp)\cdot\widetilde{\mathbf{E}}(\mathbf{k}_\perp|z^{\text{max}})\right), \tag{S81a}$$

$$\widetilde{\mathbf{H}}(\frac{\mathbf{k}_\perp}{M}|z^c) = M\left[\frac{(n_{\text{collection}}k_0)^2-|\mathbf{k}_\perp|^2}{k_0^2-\frac{1}{M^2}|\mathbf{k}_\perp|^2}\right]^{\frac{1}{4}} P(\mathbf{k}_\perp)e^{-ik_z z^{\text{max}}}\left(\overline{\overline{\mathbf{R}}}(\mathbf{k}_\perp)\cdot\widetilde{\mathbf{H}}(\mathbf{k}_\perp|z^{\text{max}})\right), \tag{S81b}$$

where $\widetilde{\mathbf{E}}(\mathbf{k}_\perp|z^{\text{max}})$ and $\widetilde{\mathbf{H}}(\mathbf{k}_\perp|z^{\text{max}})$ are the electric and magnetic fields at the $z = z^{\text{max}}$ plane, respectively.

We now consider the formation of a three-dimensional $z$-stack. When imaging a three-dimensional structure with a microscope, the objective is typically focused sequentially on different $xy$ planes, with an image acquired at each focal position. These images are then combined to form a $z$-stack. The objective focal position is commonly controlled by translating the objective along the optical axis. Figure S6 illustrates the cases in which the objective focal plane $z_f$ is shifted in the negative and positive $z$ directions. Under ideal imaging conditions, the objective should not contact or mechanically displace the specimen. Therefore, we require $z_f > z^{\text{max}} - \frac{f_o}{n_{\text{collection}}}$.

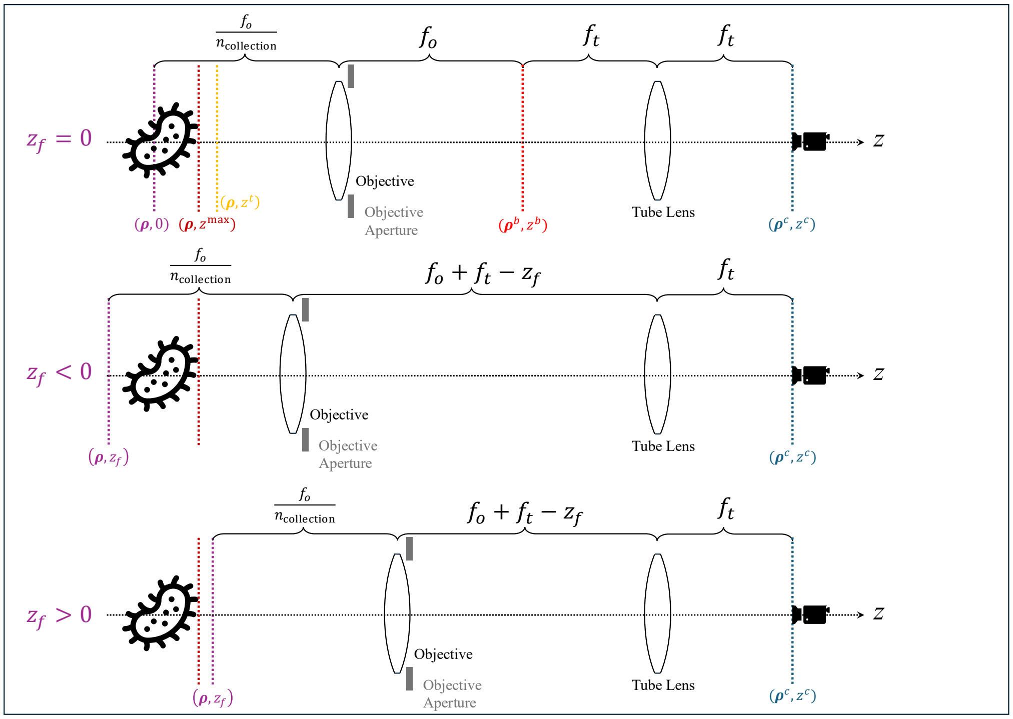


**Fig. S10. Objective-scanning geometry for generating images at different axial focal positions.** The three configurations illustrate objective focal positions at $z_f = 0$, $z_f < 0$, and $z_f > 0$. The changing propagation distances provide the basis for calculating the corresponding camera-plane fields and assembling the images into a z-stack.

When the objective is focused at plane $z = z_f$, the distance between the objective focal plane and the $z = z^{\max}$ plane becomes $z^{\max} - z_f$. Thus the Richards-Wolf integral Eq. S66 becomes

$$\hat{\mathbf{E}}'(\boldsymbol{\rho}^b|z^b) = C_o P_o'(\boldsymbol{\rho}^b) \left[ \sqrt{\gamma_o(\mathbf{k}_\perp)}\, \overline{\overline{\mathbf{R}}}_o(\mathbf{k}_\perp) e^{-ik_z(z^{\max}-z_f)} \widetilde{\mathbf{E}}(\mathbf{k}_\perp|z^{\max}) \right]_{\substack{\mathbf{k}_\perp=(k_x,k_y) \\ k_x=k_0 x^b/f_o \\ k_y=k_0 y^b/f_o}}, \tag{S82}$$

and the electromagnetic field at the camera plane is

$$\begin{aligned} \widetilde{\mathbf{E}}'(\frac{\mathbf{k}_\perp}{M}|z^c) &= M \left[ \frac{(n_{\text{collection}}k_0)^2 - |\mathbf{k}_\perp|^2}{k_0^2 - \frac{1}{M^2}|\mathbf{k}_\perp|^2} \right]^{\frac{1}{4}} P(\mathbf{k}_\perp) e^{-ik_z(z^{\max}-z_f)} \left( \overline{\overline{\mathbf{R}}}(\mathbf{k}_\perp) \cdot \widetilde{\mathbf{E}}(\mathbf{k}_\perp|z^{\max}) \right) \\ &= \widetilde{\mathbf{E}}(\frac{\mathbf{k}_\perp}{M}|z^c) e^{ik_z z_f}, \qquad \text{(S83a)} \\ \widetilde{\mathbf{H}}'(\frac{\mathbf{k}_\perp}{M}|z^c) &= \widetilde{\mathbf{H}}(\frac{\mathbf{k}_\perp}{M}|z^c) e^{ik_z z_f}. \qquad \text{(S83b)} \end{aligned}$$

Therefore, when the objective focal plane is located at $z = z_f$, the 2D image captured by the camera can be expressed as

$$\hat{\mathbf{E}}(M\boldsymbol{\rho}, z_f) = \frac{1}{(2\pi M)^2} \iint_{-\infty}^{\infty} \widetilde{\mathbf{E}}(\frac{\mathbf{k}_\perp}{M}|z^c) e^{ik_z z_f} e^{i\mathbf{k}_\perp\cdot\boldsymbol{\rho}} \mathrm{d}\mathbf{k}_\perp, \tag{S84a}$$

$$\hat{\mathbf{H}}(M\boldsymbol{\rho}, z_f) = \frac{1}{(2\pi M)^2} \iint_{-\infty}^{\infty} \widetilde{\mathbf{H}}(\frac{\mathbf{k}_\perp}{M}|z^c) e^{ik_z z_f} e^{i\mathbf{k}_\perp\cdot\boldsymbol{\rho}} \mathrm{d}\mathbf{k}_\perp \tag{S84b}$$

$$\mathbf{S}(M\boldsymbol{\rho}, z_f) = \frac{1}{2}\mathrm{Re}\{\hat{\mathbf{E}}(M\boldsymbol{\rho}, z_f) \times \hat{\mathbf{H}}^*(M\boldsymbol{\rho}, z_f)\}, \quad I(M\boldsymbol{\rho}, z_f) = \mathbf{S}(M\boldsymbol{\rho}, z_f) \cdot \hat{\mathbf{n}}, \tag{S84c}$$

where $k_z(\mathbf{k}_\perp) = \sqrt{(n_{\text{collection}}k_0)^2 - |\mathbf{k}_\perp|^2}$ and $\hat{\mathbf{n}} = [0, 0, 1]$. A $z$-stack is generated by repeatedly calculating the image at different values of $z_f$ and subsequently stacking the resulting images.

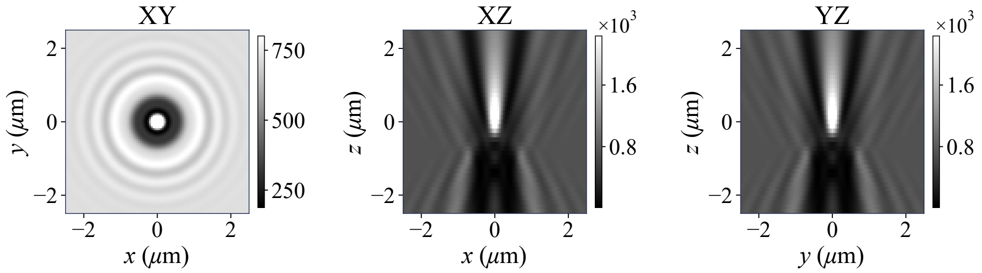


**Fig. S11. Three-dimensional brightfield image structure under single-plane-wave illumination.** The $xy$, $xz$, and $yz$ cross-sections show the simulated z-stack of a 1-µm-diameter polystyrene bead illuminated by a normally incident plane wave. The transverse and axial views illustrate the spatial distribution of image intensity as the objective focal position is varied.

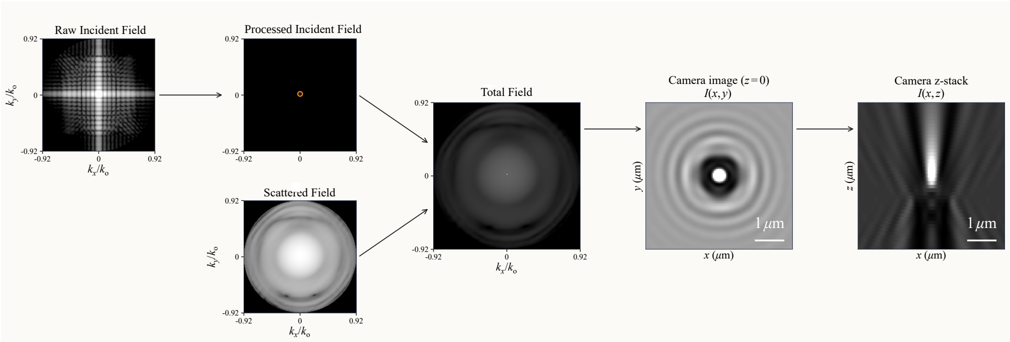


**Fig. S12. Computational workflow for generating a z-stack from a single illumination realization.** The incident-field spectrum is first corrected and combined with the scattered-field spectrum through the objective–tube-lens model. Fourier transformation and Poynting-vector evaluation yield the camera image, while propagation to different focal positions produces the complete z-stack.

Figure S11 shows $xy$, $xz$, and $yz$ cross-sections of the three-dimensional image of a 1-µm-diameter polystyrene bead under normally incident plane-wave illumination.

Figure S12 summarizes the complete workflow for generating a z-stack under single-plane-wave illumination. First, two simulations are performed to obtain the incident and scattered fields. The incident field is then preprocessed to recover an ideal plane-wave representation. Next, the objective–tube-lens transformation is applied separately to both fields, which are summed to obtain the camera-side total field in the spatial-frequency domain. A Fourier transform yields the spatial-domain fields, from which the image in a single $xy$-plane is calculated using the Poynting vector. Repeating this calculation at different focal positions through free-space propagation produces the complete z-stack.

### K. Partial Coherence in Brightfield Microscopy

Eq. S84c describes image formation under illumination by a plane wave with a single temporal frequency and propagation direction $(\omega, \theta, \psi)$. Therefore, its more general form can be expressed as $I(M\boldsymbol{\rho}, z_f) = I(M\boldsymbol{\rho}, z_f|\omega, \theta, \psi)$. As discussed previously, however, in brightfield microscopy using a broadband, spatially extended LED source, the condenser back aperture plane produces plane-wave components with different temporal frequencies and propagation directions. These components jointly illuminate the specimen and contribute to the final image. This imaging condition is described by the partial-coherence model. Analogous to the decomposition of the incident-field intensity in Eq. S46, the intensity generated by the total field at the camera plane can be decomposed over the individual source realizations. Therefore, after accounting for the source spectrum and illumination NA, the final image obtained when the objective focal plane is

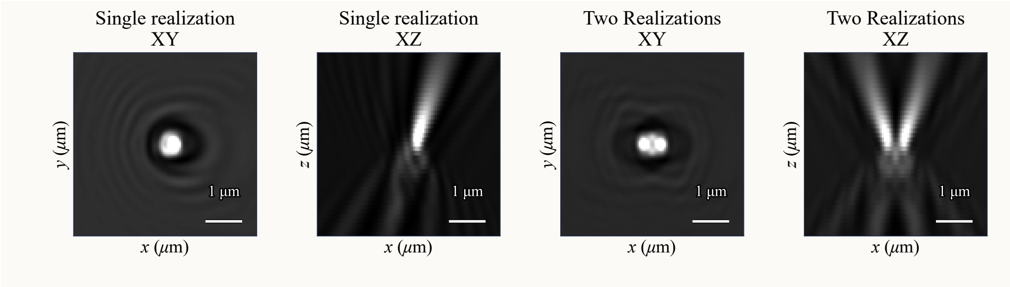


**Fig. S13. Image formation through incoherent combination of different illumination directions.** The left two panels show $xy$ and $xz$ cross-sections for a single plane wave with $\omega = 3.77 \times 10^{15}$ rad, $\mathrm{s}^{-1}$, $\theta = 10°$, and $\psi = 0°$. The right two panels show the intensity sum for two mutually incoherent plane waves with the same frequency and polar angle but azimuthal angles of $0°$ and $180°$. Their separate contributions produce spatially offset bright features in the combined image.

located at $z = z_f$ can be expressed as:

$$I_{\text{tot}}(M\boldsymbol{\rho}, z_f) = P_0 \int_{-\infty}^{\infty} p_\omega(\omega) \int_0^{\theta_{\max}^{\text{Illu}}} \int_0^{2\pi} p_{\theta,\psi}(\theta, \psi|\omega) I(M\boldsymbol{\rho}, z_f|\omega, \theta, \psi) \mathrm{d}\psi \mathrm{d}\theta \mathrm{d}\omega. \tag{S85}$$

Similarly, the $z$-stack is generated by simulating images at different focal-plane positions and stacking the resulting images.

Figure S13 illustrates image formation by summing the intensity contributions from plane waves with different parameters $(\omega, \theta, \psi)$. The two panels on the left show the $xy$ and $xz$ cross-sections of the image formed under illumination by a single plane wave with $\omega = 3.77 \times 10^{15}$ rad $\mathrm{s}^{-1}$ (vacuum wavelength 0.5μm), $\theta = 10°$, and $\psi = 0°$. The two panels on the right show the corresponding cross-sections obtained by incoherently combining two plane-wave contributions with the same wavelength and polar angle but azimuthal angles of $\psi = 0°$ and $180°$. Their distinct propagation directions produce spatially offset bright regions that follow separate axial trajectories. These features coexist without mutual interference, consistent with the assumption of mutually incoherent illumination components in the partially coherent illumination model. The complete microscope image is obtained by summing contributions from a larger ensemble of plane waves spanning different $(\omega, \theta, \psi)$ values, weighted by the source distributions.

### L. Theoretical $p_{\theta,\psi}(\theta, \psi|\omega)$ under uniform condenser aperture model

An ideal angular distribution of the incident illumination $p_{\theta,\psi}(\theta, \psi|\omega)$ can be derived under a prior source brightness distribution on condenser aperture plane. If the model assumes that the light source magnitude is uniform across the whole condenser aperture plane, then we can set $|U^c(\boldsymbol{\rho}^c|z^c, \omega)| = |U_0|, \forall \boldsymbol{\rho}^c$, where $|U_0|$ is a constant. It should be noted that the objective introduces an additional apodization factor during the objective–tube-lens transformation, as shown in Eq. S66. This factor cancels the $1/\sqrt{\cos\theta}$ term in Eq. S31. At the tube lens, the propagation angles are sufficiently small for the paraxial approximation to apply, allowing its associated apodization factor to be approximated as unity. Therefore, upon reaching the camera plane,

$$\begin{aligned} p_{\theta,\psi}(\theta, \psi|\omega) &\propto \frac{|U^{c'''}(\theta, \psi|\omega)\sqrt{\cos\theta}|^2}{\sin\theta\cos\theta} \\ &= \frac{|U_0|^2 \sin^2\theta \cos^2\theta}{\sin\theta\cos\theta} \\ &\propto \sin\theta\cos\theta \end{aligned} \tag{S86}$$

So the normalized distribution is

$$p_{\theta,\psi}(\theta, \psi|\omega) = \frac{\sin\theta\cos\theta}{\int_0^{\theta_{\max}^{\text{Illu}}} \int_0^{2\pi} \sin\theta\cos\theta \mathrm{d}\psi \mathrm{d}\theta}. \tag{S87}$$

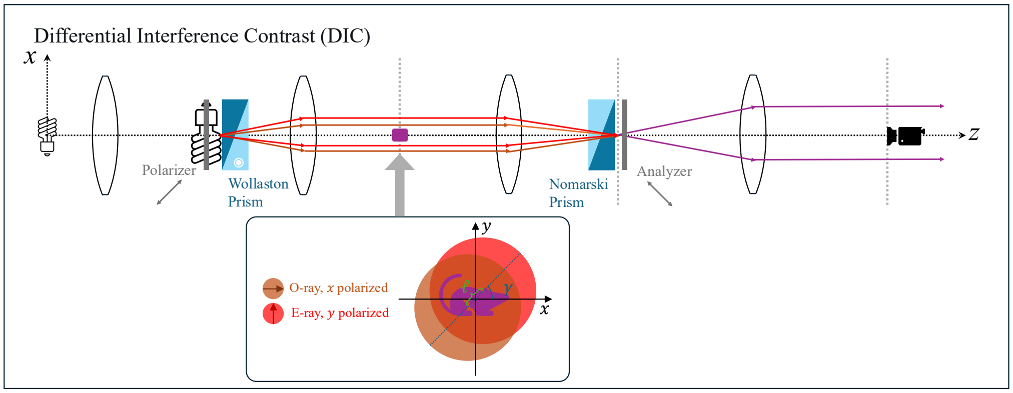


**Fig. S14. Optical configuration, polarization splitting, and lateral shear in DIC microscopy.** The polarizer and Wollaston prism generate two orthogonally polarized illumination channels that probe laterally displaced specimen positions. The Nomarski prism recombines the two channels, and the analyzer projects their fields onto a common polarization direction to enable interference. The inset illustrates the shear geometry in the specimen plane.

Therefore, the angular distribution, or the PDF of plane wave propagation angle, $p_{\theta,\psi}(\theta, \psi|\omega)$ is uniform with respect to $\psi$ and proportional to $\sin\theta\cos\theta$ under ideal uniform condenser aperture model.

### 2. THEORY OF DIC MICROSCOPY

Differential interference contrast (DIC) microscopy extends the brightfield optical pathway by splitting the illumination into two orthogonally polarized components, introducing a lateral shear between their specimen coordinates, and recombining the resulting fields through an analyzer [8], as shown in Fig. S14. The two components originating from the same source realization retain a definite relative phase and interfere at the camera, whereas distinct source realizations are averaged according to the spectral and angular distributions. This distinction between interference within a realization and intensity averaging across realizations is central to partially coherent DIC image formation.

The FDTD calculation, near-to-far-field transformation, and objective–tube-lens transformation derived for brightfield microscopy remain applicable. In the following, we introduce the modifications required for polarization splitting, specimen shear, and field recombination. As before, we first describe image formation for a single realization with parameters $(\omega, \theta, \psi)$ and then explain how contributions from different realizations are combined.

#### A. Polarization Splitting and Lateral Shear

As shown in Fig. S14, after passing through the polarizer, the incident plane wave becomes linearly polarized. Without loss of generality, we assume that the incident light is normally propagating and linearly polarized at 45° to the $x$-axis in the $xy$-plane. This linear polarization can be decomposed into two orthogonal components linearly polarized along the $x$- and $y$-axes, here designated as the ordinary (O) and extraordinary (E) rays, respectively. Upon entering the Wollaston prism, the two components undergo polarization-dependent angular splitting due to the prism's birefringence.

The condenser then converts this angular separation into a lateral displacement at the specimen plane, producing two parallel illumination beams. This lateral shear is characterized by the shearing distance and shearing angle. Let $\gamma$ denote the shear angle measured from the positive $x$ axis in specimen coordinates, and let $\ell$ denote the shearing distance between O and E rays. We define $\boldsymbol{\ell} = \ell[\cos\gamma, \sin\gamma, 0]$. Symmetric shifts are used to keep the midpoint of the two virtual specimen positions at the origin.

In general, for each realization $(\omega, \theta, \psi)$, the incident field in the sample region $\mathbf{r}^s$ can be expressed as

$$\hat{\mathbf{E}}^s_{\rm inc}(\mathbf{r}^s|\theta,\psi,\omega) = \underbrace{\vec{\mathbf{e}}_o(\theta,\psi)\exp[i\mathbf{k}^c(\theta,\psi)\cdot(\mathbf{r}^s+\frac{1}{2}\boldsymbol{\ell})]}_{\hat{\mathbf{E}}^s_{{\rm inc},o}(\mathbf{r}^s|\theta,\psi,\omega)} + \underbrace{\vec{\mathbf{e}}_e(\theta,\psi)\exp[i\mathbf{k}^c(\theta,\psi)\cdot(\mathbf{r}^s-\frac{1}{2}\boldsymbol{\ell})]}_{\hat{\mathbf{E}}^s_{{\rm inc},e}(\mathbf{r}^s|\theta,\psi,\omega)}, \quad \text{(S88a)}$$

$$\hat{\mathbf{H}}^s_{\rm inc}(\mathbf{r}^s|\theta,\psi,\omega) = \underbrace{\vec{\mathbf{h}}_o(\theta,\psi)\exp[i\mathbf{k}^c(\theta,\psi)\cdot(\mathbf{r}^s+\frac{1}{2}\boldsymbol{\ell})]}_{\hat{\mathbf{H}}^s_{{\rm inc},o}(\mathbf{r}^s|\theta,\psi,\omega)} + \underbrace{\vec{\mathbf{h}}_e(\theta,\psi)\exp[i\mathbf{k}^c(\theta,\psi)\cdot(\mathbf{r}^s-\frac{1}{2}\boldsymbol{\ell})]}_{\hat{\mathbf{H}}^s_{{\rm inc},e}(\mathbf{r}^s|\theta,\psi,\omega)}, \quad \text{(S88b)}$$

where subscripts $o$ and $e$ represent O-ray and E-ray, respectively. $\vec{\mathbf{e}}_o, \vec{\mathbf{e}}_e$ and $\vec{\mathbf{h}}_o, \vec{\mathbf{h}}_e$ are unit vectors that represent the polarization direction. Therefore, the polarization vectors of O and E rays are orthogonal to each other and transverse to the incident wavevector. Therefore,

$$\vec{\mathbf{e}}_o\cdot\vec{\mathbf{e}}_e = 0, \quad \vec{\mathbf{e}}_o\cdot\mathbf{k}^c(\theta,\psi) = 0, \quad \vec{\mathbf{e}}_e\cdot\mathbf{k}^c(\theta,\psi) = 0. \quad \text{(S89)}$$

Since the electromagnetic field propegates in the homogeneous medium, the magnetic field also has the orthogonal relationship

$$\vec{\mathbf{h}}_o(\theta,\psi) = \frac{\mathbf{k}^c(\theta,\psi)}{k}\times\vec{\mathbf{e}}_o(\theta,\psi), \quad \vec{\mathbf{h}}_e(\theta,\psi) = \frac{\mathbf{k}^c(\theta,\psi)}{k}\times\vec{\mathbf{e}}_e(\theta,\psi), \quad \text{(S90a)}$$

$$\vec{\mathbf{h}}_o\cdot\vec{\mathbf{h}}_e = 0, \quad \vec{\mathbf{h}}_o\cdot\mathbf{k}^c(\theta,\psi) = 0, \quad \vec{\mathbf{h}}_e\cdot\mathbf{k}^c(\theta,\psi) = 0. \quad \text{(S90b)}$$

As in brightfield microscopy, the incident field interacts with the specimen to generate a scattered field, which interferes with the incident field to form the total field. Taking the electric field as an example, the total field can be decomposed into components associated with the O- and E-ray channels

$$\hat{\mathbf{E}}^s_{\rm tot}(\mathbf{r}^s|\theta,\psi,\omega) = \hat{\mathbf{E}}^s_{{\rm tot},o}(\mathbf{r}^s|\theta,\psi,\omega) + \hat{\mathbf{E}}^s_{{\rm tot},e}(\mathbf{r}^s|\theta,\psi,\omega), \quad \text{(S91a)}$$

$$\hat{\mathbf{E}}^s_{{\rm tot},o}(\mathbf{r}^s|\theta,\psi,\omega) = \hat{\mathbf{E}}^s_{{\rm inc},o}(\mathbf{r}^s|\theta,\psi,\omega) + \hat{\mathbf{E}}^s_{{\rm sca},o}(\mathbf{r}^s|\theta,\psi,\omega), \quad \text{(S91b)}$$

$$\hat{\mathbf{E}}^s_{{\rm tot},e}(\mathbf{r}^s|\theta,\psi,\omega) = \hat{\mathbf{E}}^s_{{\rm inc},e}(\mathbf{r}^s|\theta,\psi,\omega) + \hat{\mathbf{E}}^s_{{\rm sca},e}(\mathbf{r}^s|\theta,\psi,\omega). \quad \text{(S91c)}$$

#### B. Implementing Lateral Shear through Specimen Translation

In SPECTRAM, the lateral shear is implemented by translating the specimen within the FDTD domain for the two polarization calculations. Two separate simulations are performed to calculate the electromagnetic fields associated with the O- and E-ray illumination channels.

After defining the experimental configuration, the refractive-index distribution representing the specimen in the O-ray simulation is specified as $n_o(\mathbf{r}) = n(\mathbf{r}-\frac{1}{2}\boldsymbol{\ell})$. It yields the fields $\hat{\mathbf{E}}^s_{{\rm tot},o}(\mathbf{r}^s|\theta,\psi,\omega)$ and $\hat{\mathbf{H}}^s_{{\rm tot},o}(\mathbf{r}^s|\theta,\psi,\omega)$.

With other configuration unchanged, the E-ray simulation is performed under the refractive-index distribution $n_e(\mathbf{r}) = n(\mathbf{r}+\frac{1}{2}\boldsymbol{\ell})$. It yields the fields $\hat{\mathbf{E}}^s_{{\rm tot},e}(\mathbf{r}^s|\theta,\psi,\omega)$ and $\hat{\mathbf{H}}^s_{{\rm tot},e}(\mathbf{r}^s|\theta,\psi,\omega)$.

#### C. Recombination and Polarization Projection

As in brightfield microscopy, the total field reaches the top plane of the collection region and is mapped to the camera plane through the objective–tube-lens system. This transformation can likewise be divided into two stages. The first maps the field from the top plane $z = z^t$ to the objective back focal plane $z = z^b$. Equation S66 remains applicable and is applied independently to the fields associated with the O- and E-ray simulations.

$$\hat{\mathbf{E}}_o(\boldsymbol{\rho}^b|z^b) = C_o P_o'(\boldsymbol{\rho}^b)\left[\sqrt{\gamma_o(\mathbf{k}_\perp)}\,\overline{\overline{\mathbf{R}}}_o(\mathbf{k}_\perp)e^{-ik_z z^t}\widetilde{\mathbf{E}}_o(\mathbf{k}_\perp|z^t)\right]_{\substack{\mathbf{k}_\perp=(k_x,k_y)\\ k_x=k_0x^b/f_o\\ k_y=k_0y^b/f_o}}, \quad \text{(S92a)}$$

$$\hat{\mathbf{E}}_e(\boldsymbol{\rho}^b|z^b) = C_o P_o'(\boldsymbol{\rho}^b)\left[\sqrt{\gamma_o(\mathbf{k}_\perp)}\,\overline{\overline{\mathbf{R}}}_o(\mathbf{k}_\perp)e^{-ik_z z^t}\widetilde{\mathbf{E}}_e(\mathbf{k}_\perp|z^t)\right]_{\substack{\mathbf{k}_\perp=(k_x,k_y)\\ k_x=k_0x^b/f_o\\ k_y=k_0y^b/f_o}}. \quad \text{(S92b)}$$

As shown in Fig. S14, a Nomarski prism is positioned at the objective back focal plane in the DIC configuration and is optically conjugate to the Wollaston prism at the condenser aperture plane. When the two prisms are properly matched and aligned, the Nomarski prism recombines the spatially separated, orthogonally polarized beams generated by the Wollaston prism, i.e. O and E rays, into a common propagation path, while their polarization states remain orthogonal.

It should be noted, however, that ideal recombination is ensured only for the components of the O- and E-ray fields polarized along their respective incident polarization directions. Scattering can generate cross-polarized components that are not fully matched to the designed recombination channels. To simplify the model, we assume that these components are weak relative to the co-polarized components and do not contribute appreciably to the designed output path after the Nomarski prism. Accordingly, only the projection of each total field onto its incident polarization direction is retained, and the two projected fields are then spatially recombined. Take the electrical field as the example, only the components of $\hat{\mathbf{E}}_o(\boldsymbol{\rho}^b|z^b)$ and $\hat{\mathbf{E}}_e(\boldsymbol{\rho}^b|z^b)$ along the incident polarization directions $\vec{\mathbf{e}}_o$ and $\vec{\mathbf{e}}_e$ are retained, respectively.

In addition, the Nomarski prism spatially recombines the two laterally sheared optical paths into a common propagation path. It will also generate a global phase difference between the two rays $e^{i\Delta\phi}$. Therefore, the recombined field is

$$\hat{\mathbf{E}}_{\text{comb}}(\boldsymbol{\rho}^b|z^b) = \vec{\mathbf{e}}_o\vec{\mathbf{e}}_o^\dagger\hat{\mathbf{E}}_o(\boldsymbol{\rho}^b - \frac{1}{2}\boldsymbol{\ell}_\perp|z^b) + e^{i\Delta\phi}\vec{\mathbf{e}}_e\vec{\mathbf{e}}_e^\dagger\hat{\mathbf{E}}_e(\boldsymbol{\rho}^b + \frac{1}{2}\boldsymbol{\ell}_\perp|z^b), \tag{S93a}$$

$$\hat{\mathbf{H}}_{\text{comb}}(\boldsymbol{\rho}^b|z^b) = \vec{\mathbf{h}}_o\vec{\mathbf{h}}_o^\dagger\hat{\mathbf{H}}_o(\boldsymbol{\rho}^b - \frac{1}{2}\boldsymbol{\ell}_\perp|z^b) + e^{i\Delta\phi}\vec{\mathbf{h}}_e\vec{\mathbf{h}}_e^\dagger\hat{\mathbf{H}}_e(\boldsymbol{\rho}^b + \frac{1}{2}\boldsymbol{\ell}_\perp|z^b), \tag{S93b}$$

where $\boldsymbol{\ell}_\perp = [\boldsymbol{\ell}_x, \boldsymbol{\ell}_y]$.

After the Nomarski prism, the analyzer projects the two orthogonally polarized fields onto a common polarization direction, thereby enabling their interference. The analyzer transmission axis is typically perpendicular to that of the polarizer. For example, if the polarizer is oriented at 45° to the $x$-axis, the analyzer transmits the component polarized at 135° (equivalently, −45°). In general, if the transmission axis of analyzer is $\mathbf{a}$, the projected electrical field is

$$\hat{\mathbf{E}}_{\text{anal}}(\boldsymbol{\rho}^b|z^b) = \mathbf{a}\mathbf{a}^\dagger\hat{\mathbf{E}}_{\text{comb}}(\boldsymbol{\rho}^b|z^b). \tag{S94}$$

The same principles apply to the magnetic field.

The electromagnetic field transmitted by the analyzer is subsequently transformed to the camera plane by the tube lens. This calculation follows the same procedure as in the brightfield model. Therefore, Eqs. S72, S78a-S78c, and S84a-S84c can be used to calculate the camera-plane electromagnetic fields, the fields obtained when the objective is focused at different axial positions $z_f$, and the corresponding intensity z-stack.

Let the camera-plane electromagnetic fields associated with the O ray, when the objective is focused at $z_f$, be denoted by $\hat{\mathbf{E}}_o(M(\boldsymbol{\rho} - \frac{1}{2}\boldsymbol{\ell}_\perp), z_f)$, $\hat{\mathbf{H}}_o(M(\boldsymbol{\rho} - \frac{1}{2}\boldsymbol{\ell}_\perp), z_f)$ and those associated with the E ray be denoted by $e^{i\Delta\phi}\hat{\mathbf{E}}_e(M(\boldsymbol{\rho} + \frac{1}{2}\boldsymbol{\ell}_\perp), z_f)$, $e^{i\Delta\phi}\hat{\mathbf{H}}_e(M(\boldsymbol{\rho} + \frac{1}{2}\boldsymbol{\ell}_\perp), z_f)$. The resulting image at this focal position is then

$$\begin{aligned} I(M\boldsymbol{\rho}, z_f) = &\underbrace{\frac{1}{2}\text{Re}\{\hat{\mathbf{E}}_o \times \hat{\mathbf{H}}_o^*\} \cdot \hat{\mathbf{n}}}_{I_o(M\boldsymbol{\rho}, z_f)} + \underbrace{\frac{1}{2}\text{Re}\{\hat{\mathbf{E}}_e \times \hat{\mathbf{H}}_e^*\} \cdot \hat{\mathbf{n}}}_{I_e(M\boldsymbol{\rho}, z_f)} \\ &+ \underbrace{\frac{1}{2}\text{Re}\{e^{-i\Delta\phi}\hat{\mathbf{E}}_o \times \hat{\mathbf{H}}_e^* + e^{i\Delta\phi}\hat{\mathbf{E}}_e \times \hat{\mathbf{H}}_o^*\} \cdot \hat{\mathbf{n}}}_{\text{Interference}}. \end{aligned} \tag{S95}$$

This expression shows that, for an incident plane wave with a single temporal frequency and propagation direction $(\omega, \theta, \psi)$, the resulting DIC image comprises three contributions: the intensity of the O-ray channel, the intensity of the E-ray channel, and their interference after projection onto a common polarization direction by the analyzer.

Figure S15 shows the three individual contributions to the DIC image under normally incident plane-wave illumination, together with the resulting image. The polarizer transmission axis is oriented at 45° to the $x$-axis in the $xy$-plane. The O- and E-ray eigenpolarization directions of the Wollaston prism are aligned with the $x$- and $y$-axes, respectively, while the analyzer transmission axis is oriented at 135°. The lateral shear is directed at 45° to the $x$-axis in the $xy$-plane, with a shear distance of 0.175 $\mu$m. The relative phase difference between the O- and E-ray channels is $-0.5\pi$. The individual intensity contributions of the O- and E-ray channels exhibit the same symmetric structure as the corresponding brightfield image. In contrast, their interference term is asymmetric, forming a positive–negative dipolar pattern along the shearing direction. This interference term is therefore responsible for the characteristic asymmetric contrast of the final DIC image.

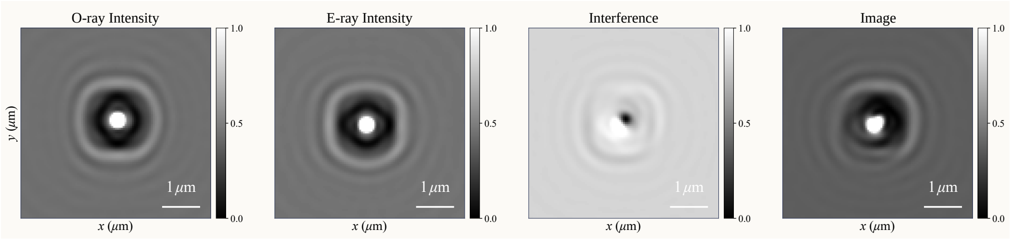


**Fig. S15. Individual intensity and interference contributions to DIC image formation.** The panels show the O-ray intensity, E-ray intensity, interference term, and resulting DIC image under normally incident plane-wave illumination. The shear distance is 0.175 $\mu$m along a direction at 45° to the $x$-axis. The asymmetric interference contribution produces the characteristic bright–dark contrast along the shearing direction.

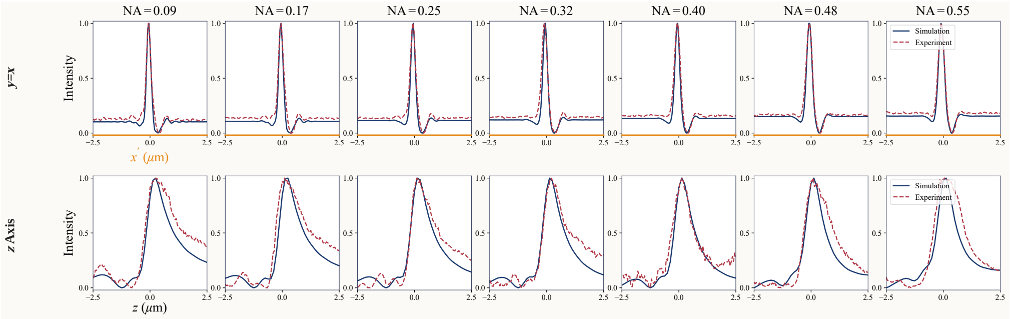


**Fig. S16. Comparison of simulated and experimental DIC line profiles across different illumination NAs.** Columns correspond to illumination NAs ranging from 0.09 to 0.55. The upper row compares lateral profiles along $y = x$ in the $z = 0$ plane, while the lower row compares axial profiles along the $z$-axis at $x = y = 0$. These profiles assess agreement in both lateral contrast and axial image structure.

### D. Partial coherence in DIC microscopy

Eq. S95 describes the DIC image formation process for single $(\omega, \theta, \psi)$. For the extended LED source considered here, we adopt the same partial coherence model as in the brightfield case. The fields in O- and E- rays in each plane wave simulation are combined coherently according. However, the fields in different plane wave simulations are mutually incoherent and are therefore combined incoherently with intensity. Hence, the final DIC image at a given focal position $z_f$ is obtained by summing the contributions from all source realizations, weighted by the source spectrum and angular distribution:

$$I_{\text{tot}}(M\boldsymbol{\rho}, z_f) = P_0 \int_{-\infty}^{\infty} p_\omega(\omega) \int_0^{\theta_{\max}^{\text{Illu}}} \int_0^{2\pi} p_{\theta,\psi}(\theta, \psi|\omega) I(M\boldsymbol{\rho}, z_f|\omega, \theta, \psi) \mathrm{d}\psi \mathrm{d}\theta \mathrm{d}\omega. \tag{S96}$$

### E. Line Profiles under different Illumination NAs

Figure 4 of the main text compares simulated and experimentally measured DIC image cross-sections at different illumination NAs. Figure S16 further evaluates the agreement between SPECTRAM simulations and experimental measurements using line profiles. The first row compares profiles along $y = x$ in the $z = 0$ plane, whereas the second row shows profiles along the $z$-axis at $x = y = 0$. The simulated and experimental profiles agree closely in both the lateral and axial directions, supporting the accuracy of the proposed model.

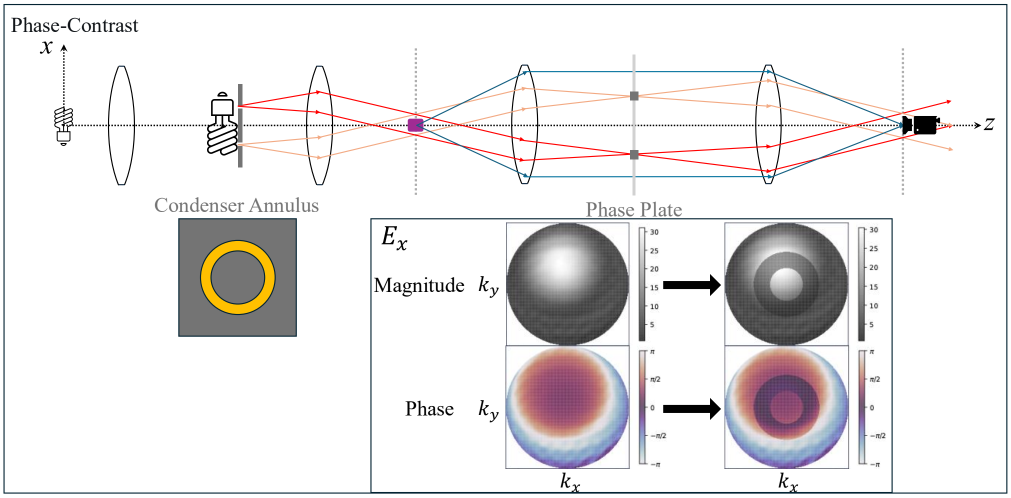


**Fig. S17. Annular illumination and phase-plate modulation in phase-contrast microscopy.** The condenser annulus restricts the illumination to an annular angular support, while the phase plate modifies the corresponding region of the objective pupil. The example spatial-frequency maps illustrate the amplitude attenuation and phase shift applied to $E_x$ within this annular region.

**Table S1.** Illumination NA ranges under different condenser annulus.

| Condenser Annulus | NA Range |
|---|---|
| Phase 1 | 0.15-0.20 |
| Phase 2 | 0.25-0.30 |
| Phase 3 | 0.45-0.55 |

### 3. THEORY OF PHASE-CONTRAST MICROSCOPY

Phase-contrast microscopy modifies the brightfield system through an annular illumination aperture and a phase plate in the conjugate objective back focal plane [9]. As shown in Fig. S17, the illumination annulus restricts the spatial frequency range of incident light, while the phase plate attenuates and phase-shifts the field within the corresponding pupil region. Interference between the modified reference field and the scattered field converts specimen-induced phase variations into intensity contrast. The full-wave specimen calculation remains unchanged; the additional optical modulation is applied to the collected angular spectrum before transformation to the camera plane.

#### A. Annular Illumination with Condenser Annulus

Unlike the spatially extended condenser apertures used in brightfield and DIC microscopy, phase-contrast microscopy introduces a condenser annulus at the condenser aperture plane. Consequently, the spatial-frequency content, or equivalently the propagation directions, of the incident illumination is restricted to a finite ring range.

The ring indicates that the polar angle of propagation direction is limited in an interval. Let $\mathrm{NA}_{\mathrm{in}}$ and $\mathrm{NA}_{\mathrm{out}}$ denote the inner and outer illumination numerical apertures of the condenser annulus. Then, we the range for polar angle $\theta$ is $[\theta_{\mathrm{in}}, \theta_{\mathrm{out}}]$, where

$$\theta_{\mathrm{in}} = \arcsin\left(\frac{\mathrm{NA}_{\mathrm{in}}}{n_{\mathrm{source}}}\right), \quad \theta_{\mathrm{out}} = \arcsin\left(\frac{\mathrm{NA}_{\mathrm{out}}}{n_{\mathrm{source}}}\right). \tag{S97}$$

However, the azimuthal angle $\psi$ remains equally distributed over $[0, 2\pi]$. Therefore, the normalized angular distribution of the incident illumination in Eq. S46 becomes

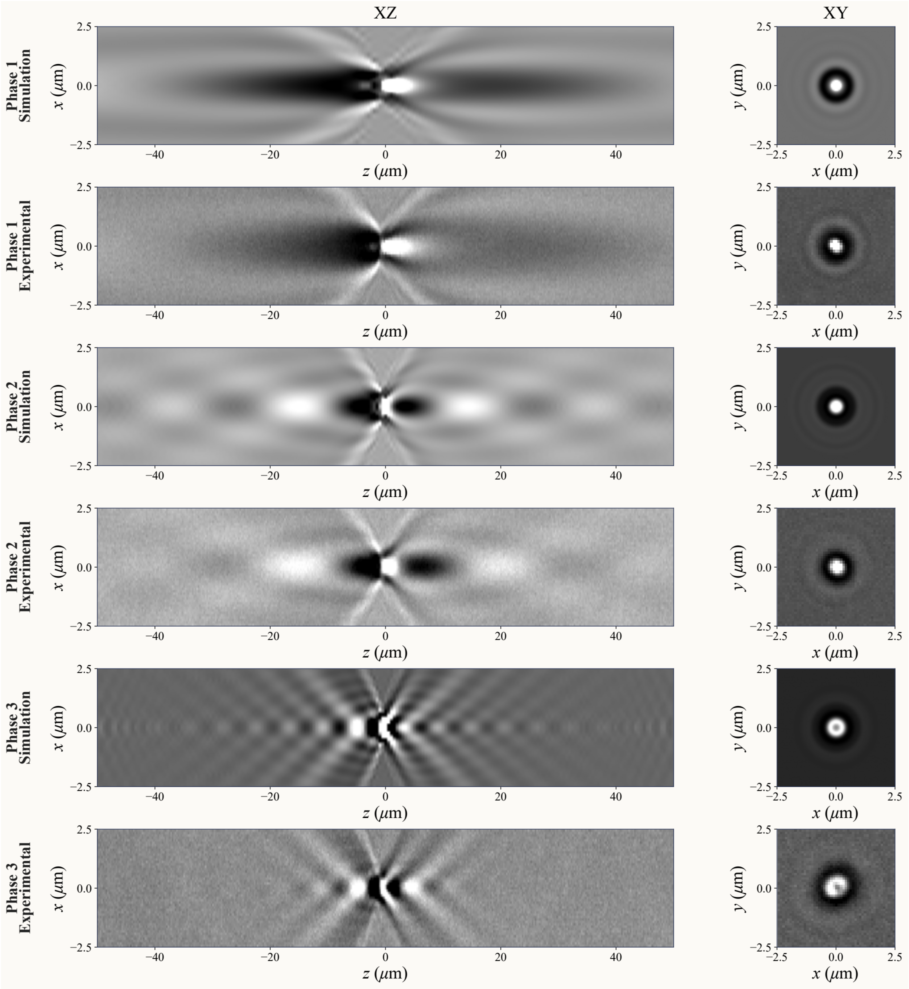


**Fig. S18. Comparison of simulated and experimental bead z-stacks under annular illumination.** A 1-µm-diameter polystyrene bead is imaged using the Ph1, Ph2, and Ph3 condenser annuli without a phase plate. For each annulus, simulated and experimental $xz$ cross-sections at $y = 0$ and $xy$ cross-sections at $z = 0$ are compared, revealing the extended axial image structure produced by annular illumination.

$$p_{\theta,\psi}^{\text{Annular}}(\theta, \psi|\omega) = \frac{p_{\theta,\psi}(\theta, \psi|\omega)}{\int_{\theta_{\text{in}}}^{\theta_{\text{out}}} \int_0^{2\pi} p_{\theta,\psi}(\theta', \psi'|\omega)\, \mathrm{d}\psi'\, \mathrm{d}\theta'}, \quad \theta_{\text{in}} \le \theta \le \theta_{\text{out}}, \quad 0 \le \psi < 2\pi. \tag{S98}$$

The Zeiss microscope provides three condenser annuli corresponding to different illumination-NA ranges. Table S1 summarizes the NA ranges provided by the three annuli evaluated in this study.

To evaluate the accuracy of SPECTRAM in modeling the condenser annulus, we first simulated brightfield images under annular illumination and compared them with experimental measurements. No phase plate was applied in this case. Figure S18 compares simulated and experimental z-stack images of a 1-µm-diameter polystyrene bead obtained using the Ph1, Ph2, and Ph3 condenser annuli. The comparison includes $xz$ cross-sections at $y = 0$ and $xy$ cross-sections at $z = 0$. The close agreement between the simulated and experimental results supports the accuracy of the annular-illumination model. Under annular illumination, the brightfield z-stack of a 1-µm-diameter bead exhibits detectable image features over an axial range approaching 100 $\mu$m. This observation indicates that the extended axial response in phase-contrast microscopy arises

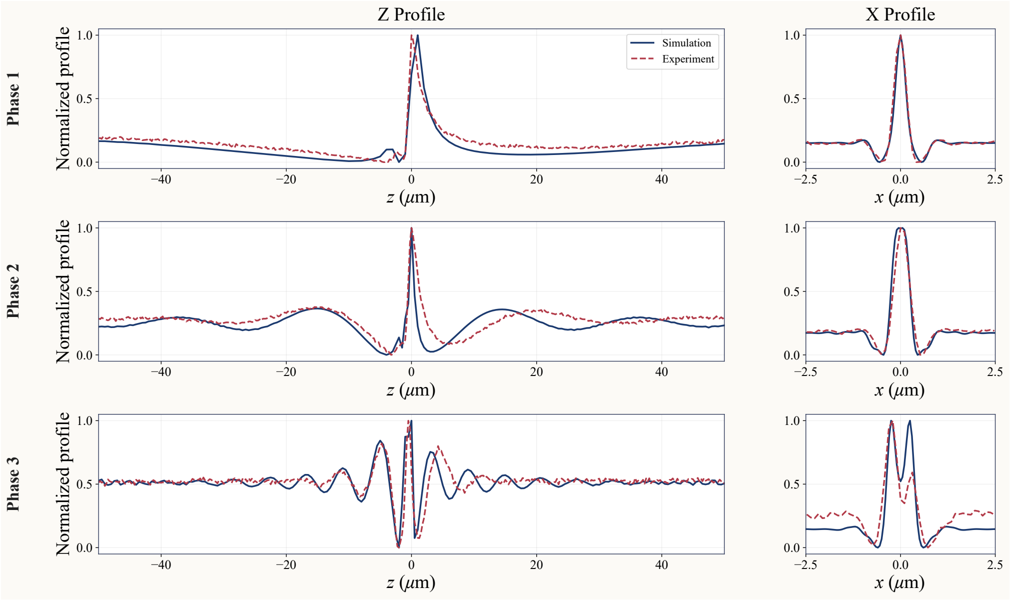


**Fig. S19. Quantitative comparison of simulated and experimental intensity profiles under annular illumination.** Rows correspond to the Ph1, Ph2, and Ph3 condenser annuli. The left column compares axial profiles along the $z$-axis at $x = y = 0$, and the right column compares lateral profiles along the $x$-axis at $y = z = 0$. All profiles are normalized for comparison.

largely from its annular illumination. We further observe a central dark region in the $xy$ images obtained with the Ph3 annulus, consistently reproduced in both simulation and experiment. This feature results from the high transverse spatial frequencies of the incident illumination, which cause the associated diffraction patterns to propagate obliquely rather than remain centered on the optical axis, as illustrated in Fig. S13.

Figure S19 further compares the simulated and experimental line profiles obtained with the three condenser annuli. The profiles are evaluated along the $z$-axis at $x = y = 0$ and along the $x$-axis at $y = z = 0$. Line profiles also show good agreement between simulation and experiment.

### B. Phase-ring modulation in the spatial-frequency domain

Another defining feature of phase-contrast microscopy is the phase plate positioned at the objective back focal plane. Its effect can be represented by a pupil-plane transmission function $T'(\boldsymbol{\rho}^b|z^b)$ acting on the spatial domain. Its definition is

$$T'(\boldsymbol{\rho}^b|z^b) = \begin{cases} \alpha e^{i\beta} & n_{\text{collection}} \sin\theta_{\text{in}} \, f_o \leq |\boldsymbol{\rho}^b| \leq n_{\text{collection}} \sin\theta_{\text{out}} \, f_o \\ 1 & \text{else} \end{cases}, \tag{S99}$$

where $\alpha$ and $\beta$ are real. They represent the amplitude attenuation and phase shift of the phase ring, respectively. The annular modulation region of the phase plate is spatially aligned with the region at the objective back focal plane occupied by the unscattered illumination generated by the condenser annulus. For the Zeiss microscope used in this study, comparison with experimental measurements suggests $\alpha \approx 0.5$ and $\beta \approx -0.3\pi$.

As in brightfield microscopy, the electromagnetic field collected from the specimen is mapped to the objective back focal plane according to Eq. S72. At this plane, the field is modified by this transmission function of the phase plate

$$\widetilde{\mathbf{E}}_{\text{pc}}(\mathbf{k}_\perp^c|z^c) = T'(\boldsymbol{\rho}^b|z^b) \cdot \widetilde{\mathbf{E}}_{\text{BF}}(\mathbf{k}_\perp^c|z^c), \quad \widetilde{\mathbf{H}}_{\text{pc}}(\mathbf{k}_\perp^c|z^c) = T'(\boldsymbol{\rho}^b|z^b) \cdot \widetilde{\mathbf{H}}_{\text{BF}}(\mathbf{k}_\perp^c|z^c). \tag{S100}$$

The modified fields are then mapped to the camera plane through the same tube-lens transformation used in the brightfield model, as described by Eq. S72, yielding the camera-side spatial-frequency-domain fields. Accordingly, Eqs. S74a and S74b are modified as follows:

$$\widetilde{\mathbf{E}}(\frac{\mathbf{k}_\perp}{M}|z^c) = M\left[\frac{(n_{\text{collection}}k_0)^2 - |\mathbf{k}_\perp|^2}{k_0^2 - \frac{1}{M^2}|\mathbf{k}_\perp|^2}\right]^{\frac{1}{4}} P(\mathbf{k}_\perp)e^{-ik_z z^t}\left(\overline{\overline{\mathbf{R}}}(\mathbf{k}_\perp)\cdot\widetilde{\mathbf{E}}_{\text{pc}}(\mathbf{k}_\perp|z^t)\right), \quad \text{(S101a)}$$

$$\widetilde{\mathbf{H}}(\frac{\mathbf{k}_\perp}{M}|z^c) = M\left[\frac{(n_{\text{collection}}k_0)^2 - |\mathbf{k}_\perp|^2}{k_0^2 - \frac{1}{M^2}|\mathbf{k}_\perp|^2}\right]^{\frac{1}{4}} P(\mathbf{k}_\perp)e^{-ik_z z^t}\left(\overline{\overline{\mathbf{R}}}(\mathbf{k}_\perp)\cdot\widetilde{\mathbf{H}}_{\text{pc}}(\mathbf{k}_\perp|z^t)\right), \quad \text{(S101b)}$$

Expressing the transmission function in spatial-frequency coordinates, we define it as follows:

$$T(\mathbf{k}_\perp) = \begin{cases} \alpha e^{i\beta} & k_0\frac{n_{\text{collection}}}{n_{\text{source}}}\text{NA}_{\text{in}} \le |\mathbf{k}_\perp| \le k_0\frac{n_{\text{collection}}}{n_{\text{source}}}\text{NA}_{\text{out}} \\ 1 & \text{else} \end{cases}. \quad \text{(S102)}$$

Then the frequency domain fields in Eqs. S101a and S101b can be expressed as

$$\widetilde{\mathbf{E}}(\frac{\mathbf{k}_\perp}{M}|z^c) = M\left[\frac{(n_{\text{collection}}k_0)^2 - |\mathbf{k}_\perp|^2}{k_0^2 - \frac{1}{M^2}|\mathbf{k}_\perp|^2}\right]^{\frac{1}{4}} T(\mathbf{k}_\perp)P(\mathbf{k}_\perp)e^{-ik_z z^t}\left(\overline{\overline{\mathbf{R}}}(\mathbf{k}_\perp)\cdot\widetilde{\mathbf{E}}(\mathbf{k}_\perp|z^t)\right), \quad \text{(S103a)}$$

$$\widetilde{\mathbf{H}}(\frac{\mathbf{k}_\perp}{M}|z^c) = M\left[\frac{(n_{\text{collection}}k_0)^2 - |\mathbf{k}_\perp|^2}{k_0^2 - \frac{1}{M^2}|\mathbf{k}_\perp|^2}\right]^{\frac{1}{4}} T(\mathbf{k}_\perp)P(\mathbf{k}_\perp)e^{-ik_z z^t}\left(\overline{\overline{\mathbf{R}}}(\mathbf{k}_\perp)\cdot\widetilde{\mathbf{H}}(\mathbf{k}_\perp|z^t)\right). \quad \text{(S103b)}$$

In the spatial-frequency domain, it is clear that the modulated annular region coincides with the support of the unscattered incident field. The phase plate attenuates this field and introduces a relative phase shift between the incident and scattered fields, thereby converting specimen-induced phase variations into stronger intensity contrast. However, as shown in Fig. S17, the transmission function acts not only on the incident field but also on scattered-field components falling within the same annular region. This unintended modulation of the scattered field contributes to the characteristic halo artifacts observed in phase-contrast microscopy.

As in brightfield microscopy, the Fourier transformation, Poynting-vector calculation, and free-space propagation described by Eqs. S78a-S78c and Eqs. S84a-S84c are then applied to calculate the phase-contrast image when the objective is focused at $z_f$ and, by repeating the calculation for different focal positions, to generate the complete z-stack.

### C. Control of Coherence between Different Realizations

The preceding formulation describes phase-contrast image formation for a single realization with parameters $(\omega, \theta, \psi)$. However, the condenser annulus produces an ensemble of plane waves spanning different values of $(\omega, \theta, \psi)$. Because the annulus confines the illumination to a relatively narrow region of transverse spatial-frequency space, its effective spatial-coherence properties differ from those of the broadly extended illumination supports used in brightfield and DIC microscopy. In the latter modalities, fields generated by distinct source realizations are assumed to be mutually incoherent because the spatially extended source spans a broad angular range. We therefore adopt the absolutely incoherent limit in Eq. S46, in which the final image is obtained by summing the intensity contributions from different $(\omega, \theta, \psi)$ realizations. The absolutely incoherent treatment is itself an approximation that neglects residual coherence between different illumination realizations. For brightfield and DIC microscopy, this approximation is generally sufficient.

In phase-contrast microscopy, however, the condenser annulus confines the illumination to a relatively narrow angular range, resulting in stronger effective coherence among the incident plane-wave components. Consequently, the fully incoherent model may not accurately reproduce phase-contrast images. Coherence is an ensemble-level statistical property that is conventionally characterized by averaging over a large number of realizations with randomly distributed initial phases. In SPECTRAM, however, each FDTD realization is computationally expensive, making direct estimation of coherence through a sufficiently large ensemble impractical. We therefore introduce an effective strategy to approximate the inter-realization coherence in phase-contrast image formation.

We consider two limiting cases. The first is the fully incoherent limit between different realizations, as described by Eq. S46. In this case, when the objective is focused at $z_f$, the resulting image is given by

$$I_{\text{incoh}}(M\boldsymbol{\rho}, z_f) = P_0 \int_{-\infty}^{\infty} p_\omega(\omega) \int_0^{\theta_{\max}^{\text{Illu}}} \int_0^{2\pi} p_{\theta,\psi}^{\text{Annular}}(\theta, \psi|\omega) I(M\boldsymbol{\rho}, z_f|\omega, \theta, \psi) \mathrm{d}\psi \mathrm{d}\theta \mathrm{d}\omega. \quad \text{(S104)}$$

The other limiting case assumes complete mutual coherence between realizations with identical initial phases. Following Eq. S40, the complex electromagnetic fields from the different realizations are coherently summed before the image intensity is calculated:

$$\hat{\mathbf{E}}_{\text{coh}}(M\boldsymbol{\rho}, z_f|\omega) = B(\omega) \int_0^{\theta_{\max}^{\text{Illu}}} \int_0^{2\pi} \sqrt{p_{\theta,\psi}^{\text{Annular}}(\theta, \psi|\omega)} \hat{\mathbf{E}}(M\boldsymbol{\rho}, z_f|\omega, \theta, \psi) \mathrm{d}\psi \mathrm{d}\theta, \quad \text{(S105a)}$$

$$\hat{\mathbf{H}}_{\text{coh}}(M\boldsymbol{\rho}, z_f|\omega) = B(\omega) \int_0^{\theta_{\max}^{\text{Illu}}} \int_0^{2\pi} \sqrt{p_{\theta,\psi}^{\text{Annular}}(\theta, \psi|\omega)} \hat{\mathbf{H}}(M\boldsymbol{\rho}, z_f|\omega, \theta, \psi) \mathrm{d}\psi \mathrm{d}\theta, \quad \text{(S105b)}$$

where $B(\omega)$ is a normalization factor such that $B(\omega) = \left[\int_0^{\theta_{\max}^{\text{Illu}}} \int_0^{2\pi} p_{\theta,\psi}^{\text{Annular}}(\theta, \psi|\omega) \mathrm{d}\psi \mathrm{d}\theta\right]^{-1}$. The image is then calculated as

$$I_{\text{coh}}(M\boldsymbol{\rho}, z_f) = P_0 \int_{-\infty}^{\infty} p_\omega(\omega) \left[\frac{1}{2} \mathrm{Re}\{\hat{\mathbf{E}}_{\text{coh}}(M\boldsymbol{\rho}, z_f|\omega) \times \hat{\mathbf{H}}_{\text{coh}}^*(M\boldsymbol{\rho}, z_f|\omega)\} \cdot \hat{\mathbf{n}}\right] \mathrm{d}\omega. \quad \text{(S106)}$$

Realizations with different temporal frequencies are still treated as mutually incoherent, and their contributions are therefore combined through intensity-based summation.

The effective coherence associated with the condenser annulus lies between these two limiting cases. We therefore model the resulting image as a weighted combination of the images predicted under the fully coherent and fully incoherent limits:

$$I_{\text{tot}}(M\boldsymbol{\rho}, z_f) = w\bar{I}_{\text{coh}}(M\boldsymbol{\rho}, z_f) + (1 - w)\bar{I}\text{incoh}(M\boldsymbol{\rho}, z_f), \quad \text{(S107)}$$

where $w$ is a weighting factor that controls the coherence between realizations. The two images are normalized before the weighted summation. This approach allows us to effectively capture the partial coherence effects in phase-contrast microscopy without the need for extensive ensemble averaging, thereby reducing computational complexity while maintaining accuracy in the simulated images.

Similarly, the z-stack is generated by calculating the image at each focal position $z_f$ and stacking the resulting images along the axial direction.

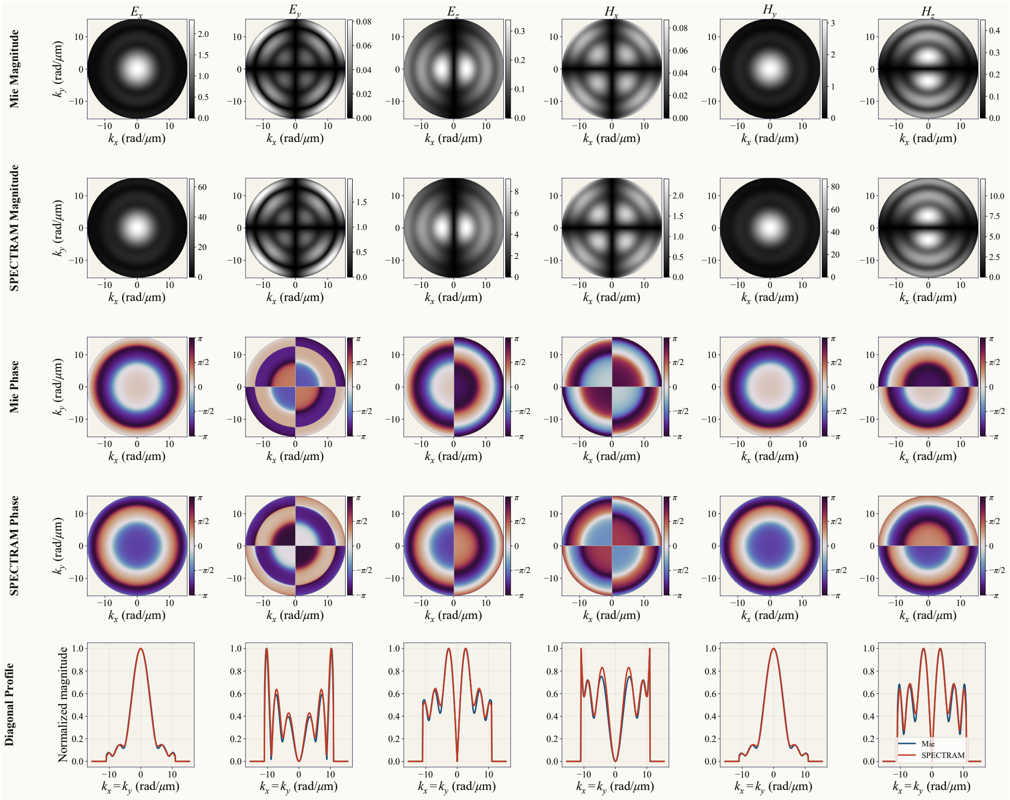

**Fig. S20. Validation of the simulated scattered-field spectra against Mie theory.** The magnitude and phase of all six electromagnetic-field components are compared in the transverse spatial-frequency domain for a 1-µm-diameter polystyrene bead under normally incident, $x$-polarized illumination. The bottom row compares normalized magnitude profiles along $k_x = k_y$. The phase maps are shown with the global phase offset between the two calculations retained.

### 4. MIE THEORY VALIDATION

As described in the main text, we used SPECTRAM to calculate the scattered field generated by a 1-µm-diameter polystyrene bead under plane-wave illumination and compared the result with Mie theory [10]. Specifically, the bead was illuminated by an $x$-polarized plane wave propagating along the optical axis. For this homogeneous spherical structure, Mie theory provides an exact analytical solution for the scattered field, enabling direct validation of the SPECTRAM simulation.

The validation was performed in both the spatial-frequency and spatial domains. Figure S20 compares the magnitude and phase obtained using SPECTRAM and Mie theory over the two-dimensional transverse spatial-frequency domain $(k_x, k_y)$. The two results exhibit close agreement in magnitude. Their phase distributions also agree apart from a global phase offset, which prevents the absolute phase values from being directly identical.

We further validated the results in the spatial domain, as shown in Fig. S21. Using Eqs. S84a-S84c, we calculated the scattered field in the two-dimensional $xy$-plane and propagated it through three-dimensional space to obtain the corresponding $xz$ cross-section. The SPECTRAM and Mie-theory results remain in close agreement, except for a modest discrepancy in the negative-$z$ region of the axial line profile.

### 5. MEASUREMENTS OF SPECTRUM AND ANGULAR DISTRIBUTION

We measured both the temporal spectrum and angular distribution of the illumination produced by the Zeiss microscope. The incident light was filtered using a green filter, and the illumination NA of the condenser aperture could be precisely adjusted from 0.09 to 0.55.

Figure S22 shows the spectrum measured using a visible-light spectrometer. The spectrum

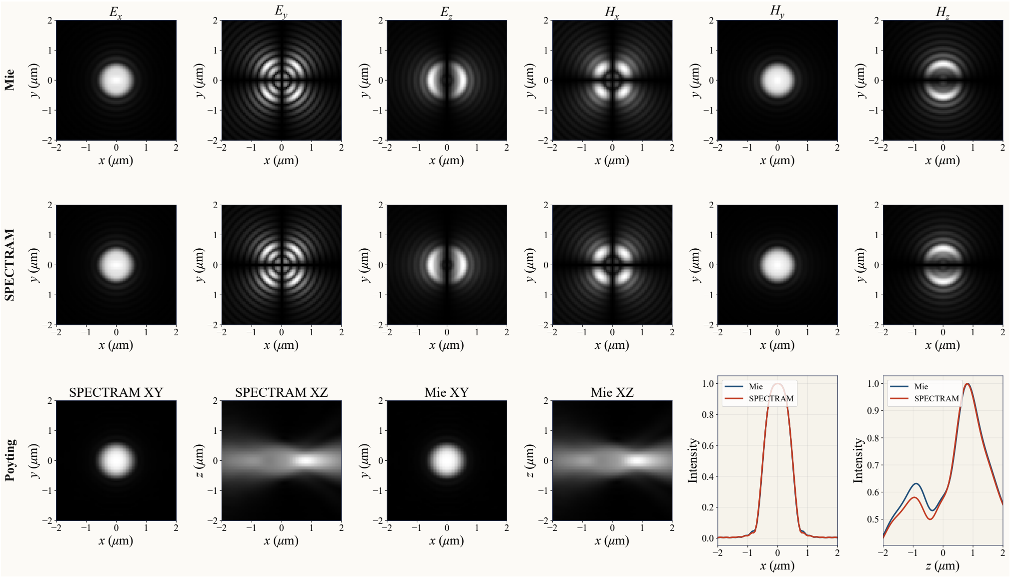


**Fig. S21. Spatial-domain comparison of SPECTRAM and Mie-theory predictions.** The upper two rows compare the magnitudes of the six scattered electromagnetic-field components in the $xy$-plane. The lower row compares the corresponding Poynting-vector-based intensity distributions in the $xy$ and $xz$ planes, together with lateral and axial line profiles.

peaks at 509 nm, with most of its intensity concentrated between 490 and 540 nm. The corresponding angular-frequency range is

$$\omega \in \left[\frac{2\pi c}{540\ \mathrm{nm}}, \frac{2\pi c}{490\ \mathrm{nm}}\right] = [3.49,\, 3.84] \times 10^{15}\ \mathrm{rad\,s^{-1}}. \tag{S108}$$

We approximate the temporal frequency spectrum $p_\omega(\omega)$ as uniform within this interval:

$$p_\omega(\omega) = \begin{cases} \dfrac{1}{\omega_{\mathrm{max}} - \omega_{\mathrm{min}}}, & \omega_{\mathrm{min}} \le \omega \le \omega_{\mathrm{max}} \\ 0, & \text{else} \end{cases}, \tag{S109}$$

where $\omega_{\mathrm{min}} = 3.49 \times 10^{15}\ \mathrm{rad\,s^{-1}}$ and $\omega_{\mathrm{max}} = 3.84 \times 10^{15}\ \mathrm{rad\,s^{-1}}$.

We also measured the angular distribution $p_{\theta,\psi}(\theta, \psi|\omega)$ of the illumination. Under otherwise identical conditions, the illumination NA was varied from 0.09 to 0.55, and the corresponding background pixel values recorded by the camera were measured. The contribution from each polar-angle interval was then estimated through finite differencing and interpolation. Assuming azimuthal symmetry, the distribution over $\psi$ was taken to be uniform.

Figure S23 shows the normalized histogram of the measured angular distribution together with the theoretical distribution derived in Eq. S87:

$$p_{\mathrm{theory}}(\theta, \psi|\omega) = \frac{\sin\theta\cos\theta}{\int_0^{\theta_{\mathrm{max}}^{\mathrm{Illu}}}\int_0^{2\pi} \sin\theta\cos\theta\,\mathrm{d}\psi\,\mathrm{d}\theta}. \tag{S110}$$

Although the theoretical prediction broadly agrees with the measurement, it assumes uniform illumination intensity across the condenser aperture plane. The measured intensity at large polar angles is lower than predicted; therefore, directly using the theoretical distribution in SPECTRAM would overemphasize the contributions from high-angle illumination. To balance the theoretical prediction with the experimentally observed behavior, we use the following modified angular distribution in SPECTRAM:

$$p_{\mathrm{sim}}(\theta, \psi|\omega) = \frac{\sqrt{\sin\theta\cos\theta}}{\int_0^{\theta_{\mathrm{max}}^{\mathrm{Illu}}}\int_0^{2\pi} \sqrt{\sin\theta\cos\theta}\,\mathrm{d}\psi\,\mathrm{d}\theta}. \tag{S111}$$

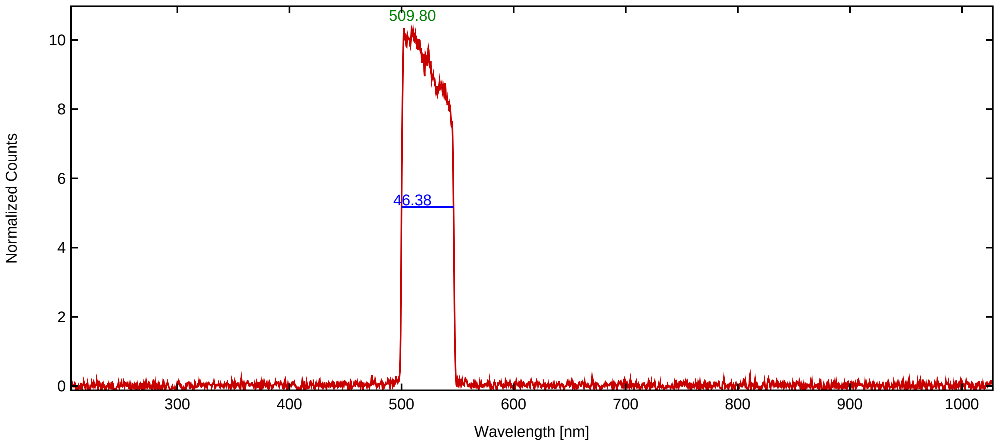


**Fig. S22. Measured illumination spectrum after green filtering.** The spectrum recorded using a visible-light spectrometer exhibits a peak at approximately 509.8 nm and an indicated bandwidth of approximately 46.4 nm. The measurement informs the spectral interval adopted in SPECTRAM, corresponding to wavelengths between 490 and 540 nm.

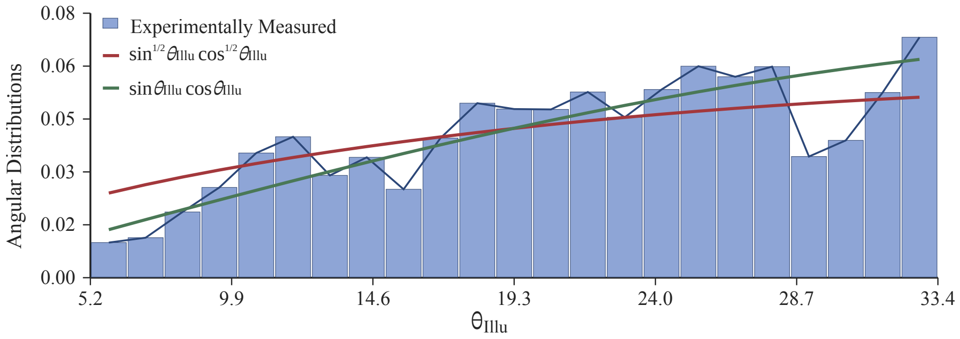


**Fig. S23. Measured and modeled angular distributions of the microscope illumination.** The normalized histogram is estimated from changes in camera background intensity as the illumination NA is varied. The curves show the theoretical distribution proportional to $\sin\theta\cos\theta$ and the modified distribution proportional to $\sqrt{\sin\theta\cos\theta}$ used in SPECTRAM, with a uniform azimuthal distribution assumed in both cases.

### REFERENCES


1. J. W. Goodman, *Introduction to Fourier Optics* (American Institute of Physics, 1969).
2. E. Dusch, T. Dorval, N. Vincent, *et al.*, "Three-dimensional point spread function model for line-scanning confocal microscope with high-aperture objective," J. microscopy **228**, 132–138 (2007).
3. K. Yee, "Numerical solution of initial boundary value problems involving maxwell's equations in isotropic media," IEEE Transactions on antennas propagation **14**, 302–307 (1966).
4. J.-P. Berenger, "A perfectly matched layer for the absorption of electromagnetic waves," J. computational physics **114**, 185–200 (1994).
5. A. F. Oskooi, D. Roundy, M. Ibanescu, *et al.*, "Meep: A flexible free-software package for electromagnetic simulations by the fdtd method," Comput. Phys. Commun. **181**, 687–702 (2010).
6. J. A. Stratton and L. Chu, "Diffraction theory of electromagnetic waves," Phys. review **56**, 99 (1939).
7. B. Richards and E. Wolf, "Electromagnetic diffraction in optical systems. ii. structure of the image field in an aplanatic system," Proc. Royal Soc. London. Ser. A, Math. Phys. Sci. pp. 358–379 (1959).
8. S. B. Mehta and C. J. R. Sheppard, "Partially coherent image formation in differential interference contrast (DIC) microscope," Opt. Express **16**, 19462–19479 (2008).
9. F. Zernike, "Phase contrast, a new method for the microscopic observation of transparent objects," Physica **9**, 686–698 (1942).
10. H. Du, "Mie-scattering calculation," Appl. optics **43**, 1951–1956 (2004).
